# A lifespan program of mouse synaptome architecture

**DOI:** 10.1101/838458

**Authors:** Mélissa Cizeron, Zhen Qiu, Babis Koniaris, Ragini Gokhale, Noboru H. Komiyama, Erik Fransén, Seth G.N. Grant

**Author notes:** these authors contributed equally (Mélissa Cizeron, Zhen Qiu).

## Abstract

How synapses change molecularly during the lifespan and across all brain circuits is unknown. We analyzed the protein composition of billions of individual synapses from birth to old age on a brain-wide scale in the mouse, revealing a program of changes in the lifespan synaptome architecture spanning individual dendrites to the systems level. Three major phases were uncovered, corresponding to human childhood, adulthood and old age. An arching trajectory of synaptome architecture drives the differentiation and specialization of brain regions to a peak in young adults before dedifferentiation returns the brain to a juvenile state. This trajectory underscores changing network organization and hippocampal physiology that may account for lifespan transitions in intellectual ability and memory, and the onset of behavioral disorders.

**One sentence summary:** The synaptome architecture of the mouse brain undergoes continuous changes that organize brain circuitry across the lifespan.

## Introduction

All animals undergo a stereotypical progression of behavioral changes across their lifespan(*1–3*). In humans, intellectual abilities are initially undifferentiated in early childhood, then differentiate to produce a multifaceted ability structure reaching a peak at ∼25 years of age(*4–7*). Intellectual abilities then slowly diminish and become undifferentiated during old age, returning to levels seen in younger individuals(*4–7*). This arching lifespan trajectory of psychological functions is known as the Differentiation-Dedifferentiation hypothesis(*5, 6*). Although these psychological trajectories suggest the existence of a biological program that controls and modifies brain circuits at every age, the identification of such programs has remained elusive. The importance of understanding the mechanisms that produce and modify the behavioral repertoire across the lifespan is underscored by studies indicating that the pathological behaviors that emerge at different ages reflect an aberration in the normal lifespan program(*8*).

In recent years it has become firmly established that the proteome composition of synapses controls the physiological mechanisms of behavior. The postsynaptic proteome of excitatory synapses (the main class of brain synapse) is highly complex(*9–13*) and genetic studies have shown that the individual proteins regulate many innate and learned behaviors in the behavioral repertoire(*14–17*). Moreover, mutations in these proteins cause over 130 brain diseases(*10*), including common disorders that characteristically arise in childhood, adolescence, young or elderly adults. How gene mutations result in synaptic pathology in particular brain areas and ages is poorly understood.

A major technical limitation to the comprehensive spatiotemporal study of synapse proteins is that current methods have only been suitable for assessing small numbers of synapses in selected brain regions. The recent development of synaptome mapping overcomes these limitations, and it is now possible to map the molecular and morphological features of billions of individual synapses across all mouse brain regions(*18*). The synaptome of the adult mouse comprises a high diversity of excitatory synapse types distributed into different brain regions, creating a three-dimensional synaptome architecture(*18*). This work has led to a new model of how the brain stores and retrieves information: the information is stored in molecularly diverse synapses and can be retrieved by their response to patterns of nerve cell activity(*18, 19*).

Here we report the first systematic analysis of the molecular and morphological properties of individual synapses across the brain and lifespan of any species. We mapped excitatory synapse diversity and the spatial and temporal synaptome architecture in >100 brain regions from birth until 18 months of age in the mouse (Fig. S1). Synapses were labelled using fluorescent tags on the endogenous PSD95 (PSD95-eGFP) and SAP102 (SAP102-mKO2) proteins(*18*). These two postsynaptic scaffold proteins assemble signaling complexes containing neurotransmitter receptors, structural proteins and signaling enzymes(*13, 20, 21*) that play a key role in synaptic plasticity and innate and learned behaviors(*15, 16, 22-24*). Using the synaptome mapping image analysis pipeline (SYNMAP)(*18*), we quantified the molecular and morphological parameters of billions of individual synapses, classified synapse types and subtypes and determined their lifespan trajectories.

Our findings reveal a remarkable spatiotemporal program that underpins the organization of synapse diversity across the brain, which we call the lifespan synaptome architecture (LSA). The LSA reveals how synapse diversity is generated, brain regions differentiate, and how the global synaptome architecture is built during development and then changes throughout adulthood and old age. Furthermore, the LSA provides a new mechanism for understanding lifespan trajectories of psychological functions and has important implications for understanding the onset and progression of brain disorders. The Mouse Lifespan Synaptome Atlas and the interactive visualization and analysis tools that we have developed provide an important new community resource for investigation of synapse function across all brain regions and the lifespan.

### The lifespan synaptome mapping pipeline and data resource

Para-sagittal brain sections from cohorts of PSD95-eGFP/SAP102-mKO2 male mice were collected at ten postnatal ages: one day (1D), one week (1W), two weeks (2W), three weeks (3W), one month (1M), two months (2M), three months (3M), six months (6M), 12 months (12M) and 18 months (18M) (Figs. 1, S2, S3). Whole brain sections were imaged at single-synapse resolution (optical resolution ∼260 nm in *xy*, pixel size 84 x 84 nm) on a spinning disc confocal microscope and the density, intensity, size and shape parameters of individual puncta were acquired using computer vision methods as previously described(*18*). These parameters were used to classify synapses into three molecular types (type 1 express PSD95 only, type 2 express SAP102 only, and type 3 express both PSD95 and SAP102) and 37 subtypes based on molecular and morphological features(*18*). Supervised synaptome maps were generated by registering the data to the Allen Reference Atlas(*25*) delineated into 109 anatomical subregions within 12 overarching regions, including isocortex, olfactory areas (OLF), hippocampal formation (HPF), cortical subplate (CTXsp), striatum (STR), pallidum (PAL), thalamus (TH), hypothalamus (HY), midbrain (MB), pons (P), medulla (MY) and cerebellum (CB).

All data and analysis tools are available in the Mouse Lifespan Synaptome Atlas (www.brain-synaptome.org). Synaptome Explorer enables in-depth exploration of raw and processed image data in single sections at individual punctum resolution, and the Synaptome Homology Viewer enables comparison of brain regions within and between mice of different ages.

### The synaptome continuously changes across the lifespan

Raw images at low and high magnification reveal that each synaptic protein has a distinct spatiotemporal pattern and that the synaptome differs at every age examined (Figs. 1A, S2, S3). To quantify the spatiotemporal differences in the synaptome, the lifespan trajectories of PSD95 and SAP102 puncta density, intensity and size were plotted as graphs and heatmaps for the whole brain, 12 regions and 109 subregions (Figs. 1B, C, S4, S5, Table S1), revealing a number of characteristic patterns. First, each parameter was continuously changing across the lifespan. For example, synapse density rapidly increases during the first month in all brain areas, then varies before declining in old age (while brain size remained unchanged, Fig. S6). Second, trajectories differ between brain areas, with each brain area undergoing a specific program of synapse development, maturation and ageing (Figs. 1B, C, S4, S5). For example, the density of synapses in the brainstem peaked before it peaked in cerebrum structures, potentially reflecting the requirement for the brainstem in early postnatal functions (Fig. 1D). Third, the two synapse proteins showed differing trajectories (Fig. 1B, C). For example, SAP102 puncta density peaks before that of PSD95 in most brain areas (Fig. 1D).

**Figure 1.**
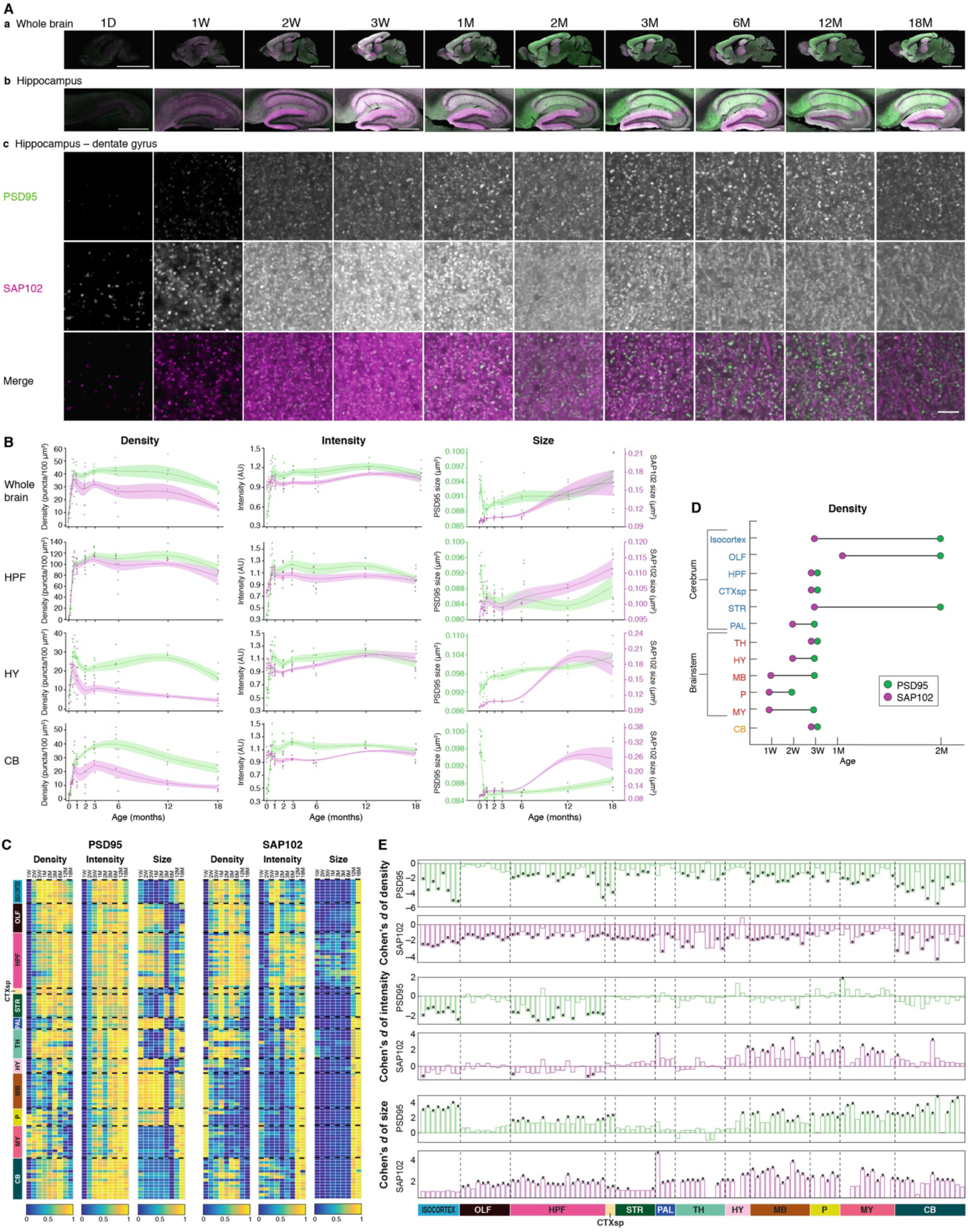
Lifespan trajectories of synapse parameters. A. PSD95-eGFP (green) and SAP102-mKO2 (magenta) expression acquired at low (20X, a and b) and high (100X, c) magnification in the whole brain (a), hippocampus (b), and molecular layer of the dentate gyrus (c) at ten ages across the mouse postnatal lifespan. Scale bars: a, 4 mm; b, 500 µm; c, 3.5 µm. D, day; W, week; M, month. B. Lifespan trajectories of synapse density (left), intensity (middle, normalized to the mean intensity, arbitrary units: AU) and size (right) in the whole brain, HPF, HY and CB. PSD95-eGFP (green) and SAP102-mKO2 (magenta). Points represent individual mice, with beta-spline smoothed curve of mean values and standard error of the mean. C. PSD95 (left) and SAP102 (right) density, intensity and size normalized from 0 to 1 for each brain subregion (see Table S1) across the lifespan. D. Age of the first peak value for the density parameter. PSD95-eGFP (green) and SAP102-mKO2 (magenta). Regions: isocortex, olfactory areas (OLF), hippocampal formation (HPF), cortical subplate (CTXsp), striatum (STR), pallidum (PAL), thalamus (TH), hypothalamus (HY), midbrain (MB), pons (P), medulla (MY), cerebellum (CB). E. Differences (Cohen’s *d*) in synapse parameters between 3M and 18M in brain subregions (see Table S1). Statistical significance calculated using Bayesian analysis(*18*) with Benjamini-Hochberg multiple comparison correction: corrected \**P* < 0.05.

Although the synaptome architecture is changing at all ages, landmarks in the trajectories indicate that the lifespan can be divided into three broad phases. The first transition point is around 1M, when the initially rapid increase in puncta density finishes. This first phase from birth to 1M corresponds to the period when the mouse is dependent on its mother and can be considered as the equivalent of childhood in humans. This is followed by a phase of relative stability until 6M (adulthood), and then there is a decline in puncta density and increase in synapse size, which is a phase equivalent to late adult life or ageing. We refer to these three phases as LSA-I, -II and - III.

The fact that there are synaptome changes throughout adult life led us to examine the synapse characteristics that distinguish the young and old adult brain and whether brain areas age in different ways. There was a loss of synapse density and an increase in their average size in almost all brain regions from 3M to 18M (Figs. 1E, S7). However, there were differences between the two synaptic proteins and between brain areas. For example, in old mice PSD95 punctum size selectively increased in isocortex whereas SAP102 punctum size increased in olfactory cortex. Examination of the size distribution of the synapse populations shows a shift toward larger synapses (effect size >0.25 with p<0.01, Kolmogorov-Smirnov test). We also found a decline in PSD95 punctum intensity in certain regions and contrasting increases in SAP102 intensity in others. Thus, trajectories in the molecular composition of individual synapses generate a changing synaptome architecture across the lifespan.

### Synapse diversity across the lifespan

To explore how the molecular trajectories result in changing populations of synapses we classified every synapse into one of three molecular types and 37 molecular/morphological subtypes using previously described methods(*18*). Dramatic changes in synapse populations occur in the first three weeks (1D – 3W, LSA-I), as shown in the stacked bar plots of synapse type (Figs. 2A, S8) and subtype density (Figs. 2B, C, S9). The synaptome of the very young brain is initially dominated by a small subset of synapse types/subtypes, and these are overtaken by expanding populations of other types/subtypes. For example, type 2 and subtype 16 synapses dominate in the first postnatal week but are a minor population by 1M.

To examine the lifespan trajectories of each synapse type we plotted heatmaps for each brain region (Fig. 2D) and subregion (Fig. 2E) and found that each synapse type within a given region/subregion has a distinct trajectory with maximal values at different ages. Similarly, heatmaps of each of the 37 subtypes showed distinct trajectories across the lifespan (Figs. 2F, S10). Importantly, these results show that the synapse composition of brain regions continues to change throughout the lifespan and is not restricted to the first phase in which synapse density increases dramatically. Moreover, the presence of alternating trends with more than one peak suggests ongoing regulatory processes that shape synapse composition with age (e.g. subtype 17 and 18, p ≤ 0.05 paired t-test and Kolmogorov-Smirnov test, Cohen’s *d* ≥1.2, Fig. S10). In old age (LSA-III), some subtypes are reduced while others increased, with differing specificity to brain regions (Fig. S11). For example, subtypes 2, 27 and 34 increased in many brain regions (Figs. 2F, S11) whereas subtypes 12-16 were lost in olfactory areas and thalamus in the old brain (Fig. S10, S11). Importantly, subtypes 2, 27 and 34 are large and subtypes 12 and 14-18 are small synapses (Fig. S9 in (*18*)). Crucially, this reveals the subtypes of excitatory synapses that are selectively gained or lost with ageing and how regions of the brain age in different ways.

**Figure 2.**
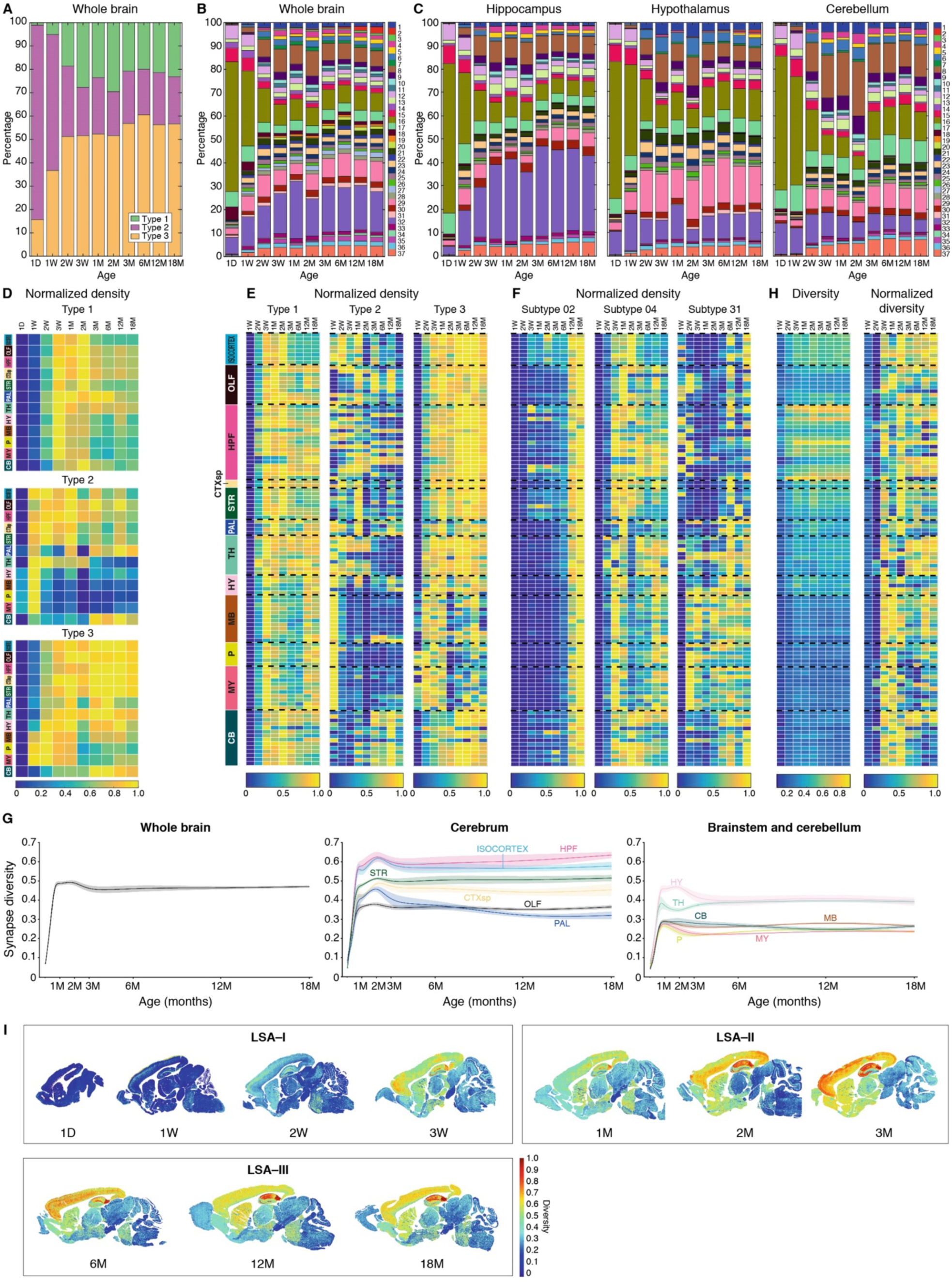
Lifespan trajectories of synapse types, subtypes and diversity. A. Stacked bar plot of percentage of synapse type density (type 1, PSD95 only; type 2, SAP102 only; type 3, colocalized PSD95+SAP102) in the whole brain across the lifespan. B. Percentage of synapse subtype density in the whole brain across the lifespan. Key: synapse subtypes (1–37). C. Percentage of synapse subtype density in hippocampus, hypothalamus and cerebellum across the lifespan. D. Lifespan trajectories of synapse type density for each of 12 main brain regions (rows). Density in each region was normalized (0-1) to its maximal density across the lifespan (columns). E. Lifespan trajectories of synapse type (normalized) density in each of 109 subregions (rows, see Table S1). Density in each subregion was normalized (0-1) to its maximal density across the lifespan (columns). F. Lifespan trajectories of synapse subtype 2, 4 and 31 (normalized) density in each of 109 subregions (rows) (see Table S1). Density in each subregion was normalized (0-1) to its maximal density across the lifespan (columns). G. Lifespan trajectories of synapse diversity (Shannon entropy) for whole brain (left) and main regions from the cerebrum (middle) and brainstem and cerebellum (right). Beta-spline smoothed curve of mean and standard error of the mean are shown. H. Raw (left) and normalized (right) values of diversity (Shannon entropy) for each of 109 subregions (rows) across the lifespan (columns) (see Table S1). The normalized diversity was obtained by min-max normalization of raw values in each subregion across the lifespan. I. Unsupervised synaptome maps showing the spatial patterning of synapse diversity (Shannon entropy) per area (pixel size 21.5 µm x 21.5 µm) in representative para-sagittal sections. LSA phases indicated.

We next quantified synapse diversity and found that all regions and subregions show a rapid initial increase in the first three postnatal weeks (Figs. 2G, H, S12). Surprisingly, brain areas responsible for higher cognitive functions (isocortex, HPF, CTXsp, STR) continued to expand their diversity, reaching a peak at 2M, whereas brain areas serving basal neurophysiological functions (MB, P, MY) peaked at 3W - 1M (Figs. 2G, H, S12). Synapse diversity declined after the peak and from 3M onwards was stable throughout the rest of the lifespan in most brain areas (Fig. 2G, H, S12). To visualize the anatomical distribution of synapse diversity, we plotted unsupervised synaptome maps of the mouse brain (Fig. 2I). The expansion in diversity in the first 3 weeks (LSA-I) is particularly dramatic, with the progressive emergence of layers in the isocortex and subregional differentiation in the HPF.

### A brain-wide trajectory of differentiation/specialization then dedifferentiation characterizes synaptome architecture across the lifespan

How the expansion in synapse diversity in LSA-I and compositional changes across the lifespan influence the higher-order organization of brain regions is unknown. To reveal how changing synapse composition might contribute to the differentiation of brain regions, we plotted the similarity matrices of brain subergions at each age (Figs. 3A, S13). The highest degree of similarity is found in the first postnatal week and rapidly diminishes during the first three weeks (when synapse diversity is expanding) and continues to diminish until 3M, when brain regions show their least similarity. Strikingly, as the brain ages beyond 3M there is a progressive increase in the similarity between brain areas (Figs. 3A, S13). We quantified this trajectory (Fig. 3B) and the topology of the synaptome network using an index of small-worldness (which reflects the patterns of the similarity matrix and was previously shown to correlate with the functional connectivity measured using resting-state functional magnetic resonance imaging of brain regions(*18*))(Fig. 3C). These analyses show that during the first three months there is a progressive differentiation and specialization of brain regions which is followed by their dedifferentiation into old age.

**Figure 3.**
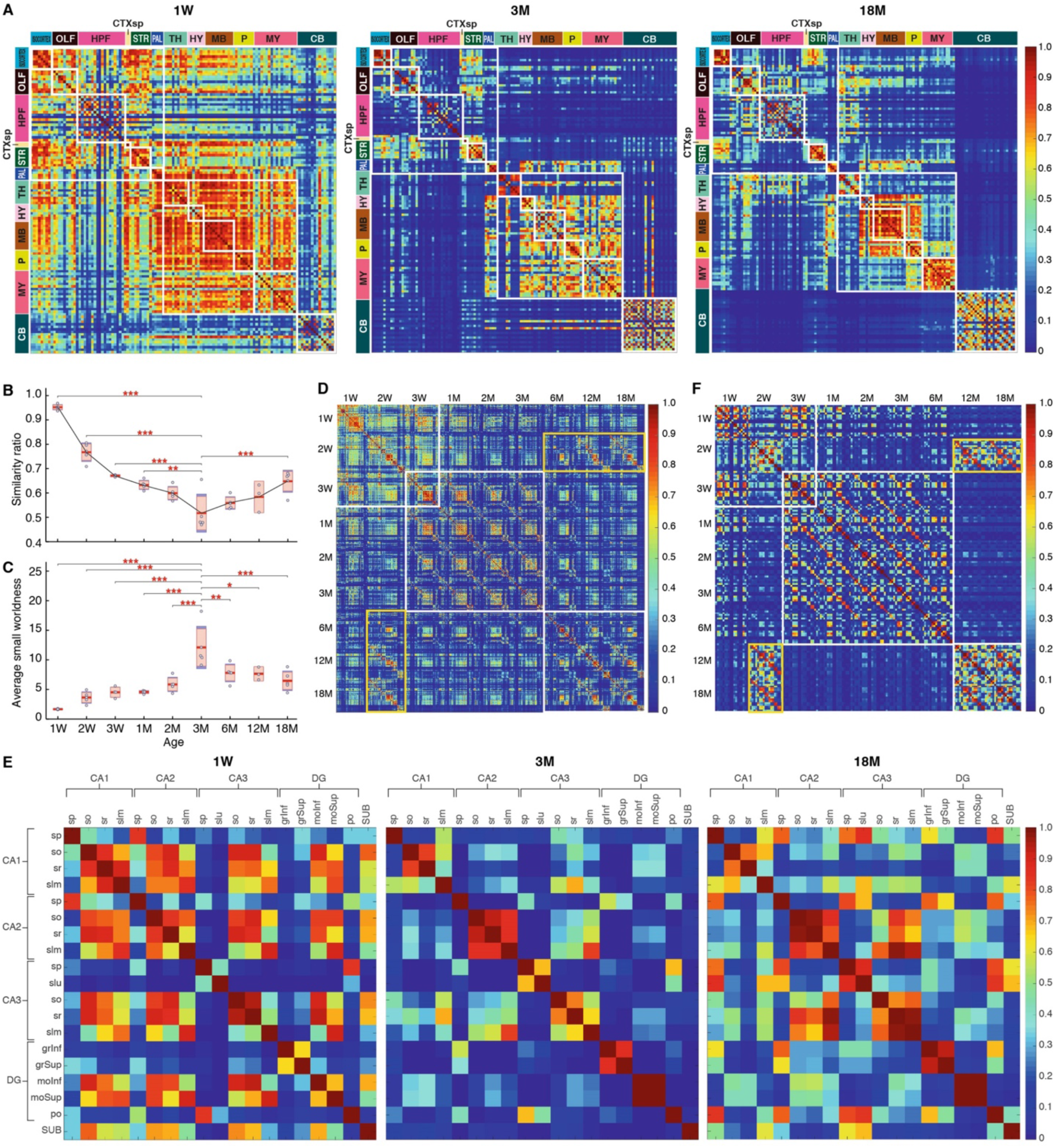
Lifespan synaptome architecture. A. Matrix of similarities between pairs of subregions (rows and columns) at 1W, 3M and 18M. Small white boxes indicate the subregions that belong to the same main brain region (see color code on the left and at top) and larger white boxes indicate main clusters: cerebrum, brainstem, cerebellum. Note reduction in similarity from 1W to 3M and increase to 18M. B. Similarity ratio indicates the relative similarity of the synaptome in each subregion with every other subregion(*18*). See Materials and Methods for details. Significant differences of ratio between 3M and other ages: **p<0.01, ***p < 0.001. C. Average small worldness across the lifespan. Scatter plots indicate the average small worldness per mouse brain section at different ages. Since small worldness can vary by different network density, a single-value average small worldness per mouse is generated by averaging a set of small-worldness values of the same animal at network densities ranging from 0 to 30%. See Materials and Methods for details. Significant differences of small worldness between 3M and other ages: *p <0.05, **p<0.01,***p < 0.001. D. Whole brain hypersimilarity matrix showing the similarity between pairs of subregions at all ages. White boxes indicate main three clusters corresponding to LSA-I, -II and -III. Yellow box shows increased similarity of the old brain with young brain. E. Matrix of similarities between pairs of hippocampal subregions (see Table S1) at 1W, 3M and 18M. Note reduction in similarity between CA1, CA2, CA3 and DG from 1W to 3M and increase to 18M. F. Hippocampus hypersimilarity matrix showing the similarity of pairs of hippocampal subregions at all ages. White boxes indicate main three clusters corresponding to LSA-I, -II and -III. Yellow box shows increased similarity of the old brain with young brain.

These findings raise the tantalizing question of whether this dedifferentiation represents a return to a synaptome resembling that of a young brain or to a distinct, elderly-specific synaptome architecture. To address this, we compared the synaptome architecture of all subregions at all ages using a hypersimilarity matrix (Figs. 3D, S14). Remarkably, the synaptome architecture of the ageing brain (6M – 18M) increasingly resembles that of the 2W mouse brain (Fig. 3D, S14 yellow boxes). This hypersimilarity matrix also effectively reveals the three lifespan phases, LSA-I, -II and -III (Fig. 3D, S14 white boxes). Interestingly, LSA-I and LSA-II overlap in week three, which coincides with the crucial behavioral transition from dependence on maternal care to independent living.

Having identified this arching trajectory in the synaptome architecture across the lifespan at the brain-wide level, we reasoned that these same mechanisms might extend to the organization of individual brain structures, such as the hippocampal formation. We generated similarity matrices between hippocampal subregions at each age (Figs. 3E, S15) and a hypersimilarity matrix comparing all ages and subregions (Figs. 3F, S16). Similar to the brain-wide analysis, the similarity matrices showed differentiation of hippocampal subregions from LSA-I to LSA-II and dedifferentiation in LSA-III. The hypersimilarity matrix also showed the three LSA phases and the similarity in synaptome architecture of the 12M −18M brain with the 2W brain.

### Lifespan synaptome architecture underwrites changes in brain functional outputs

To explore how the age-dependent changes in synaptome architecture may cause changes in cognitive functions we focused on the hippocampal formation, which plays a key role in spatial navigation, learning and memory. We previously demonstrated that in the CA1 stratum radiatum of the adult mouse there are orthogonal (radial and tangential) spatial gradients in PSD95 and SAP102 synaptic parameters that produce a local architecture of molecularly diverse synapses(*18*). The radial gradient mostly corresponds to the linear distance of synapses from the soma on apical dendrites, and the tangential gradient is formed by differences between adjacent neurons along the pyramidal cell body layer(*18*). These diverse synapses produce postsynaptic potentials of varying amplitudes in response to incoming patterns of activity and thereby generate a functional output from the synaptome map(*18*) (Fig. 4A).

**Figure 4.**
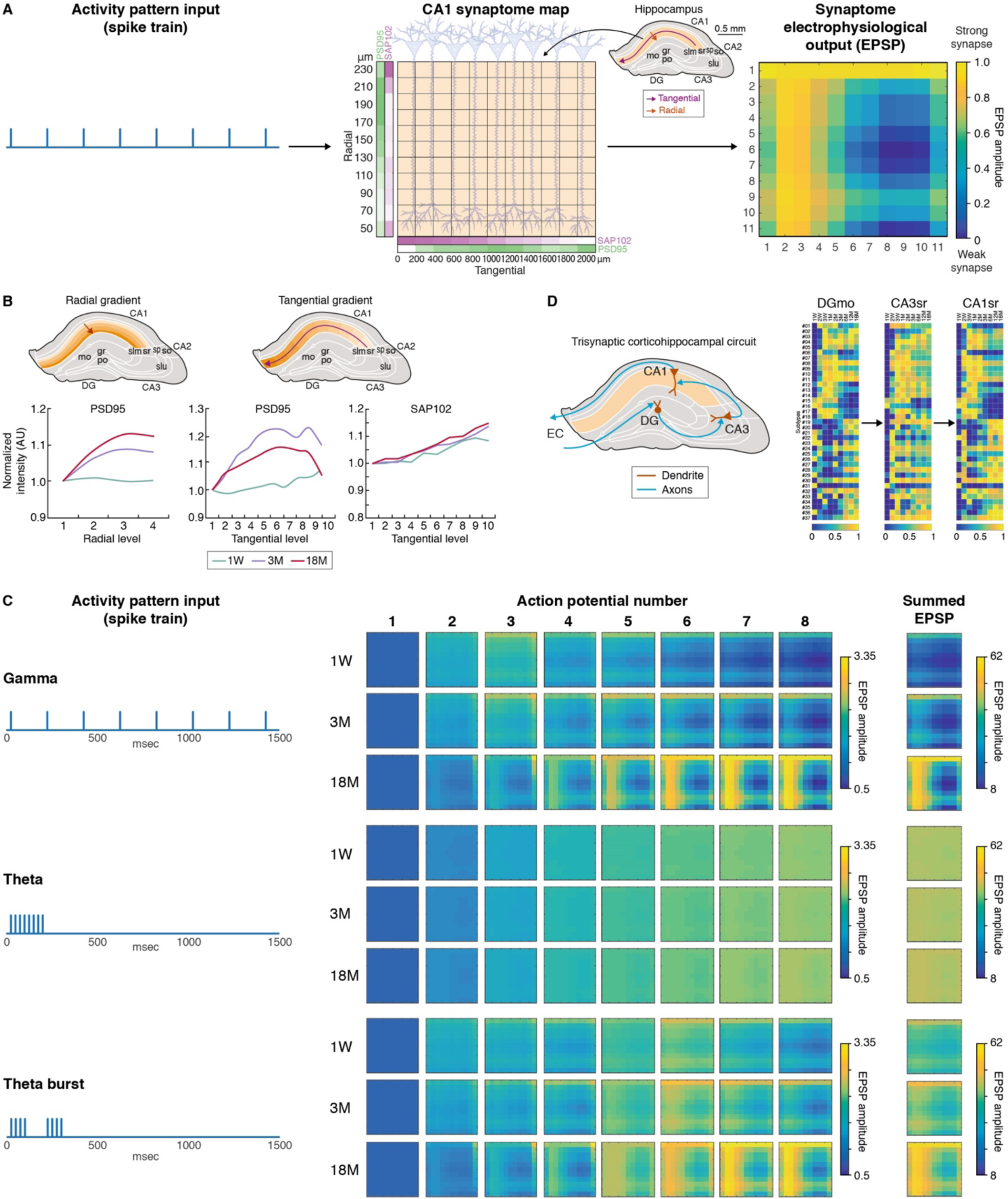
Lifespan changes in hippocampus architecture and electrophysiological properties. A. Schematic of neuronal firing pattern driving a matrix of 11 x 11 synapses expressing different amounts of PSD95 and SAP102 in the hippocampal region (adapted from(*18*)) and the spatial output of excitatory postsynaptic potentials (summed EPSP). B. Schematics of hippocampus showing radial (left) and tangential (right) gradients in CA1sr subfields. Graphs show gradients of normalized synapse intensity (AU) of PSD95 and SAP102 at 1W, 3M and 18M. C. The response (EPSP amplitude) to three patterns of eight action potentials of the 11 x 11 matrix of hippocampus synapses (see A) at three different ages (1W, 3M, 18M). Responses to each action potential and summed response are shown. D. Schematic of the trisynaptic hippocampal circuit connecting DG molecular layer (DGmo), CA3 stratum radiatum (CA3sr) and CA1 stratum radiatum (CA1sr) and the lifespan trajectory of synapse subtype density (normalized) in each region. EC, entorhinal cortex.

We quantified these tangential and radial gradients at 1W, 3M and 18M (Figs. 4B, S17). The PSD95 intensity radial and tangential gradients underwent age-dependent changes, in contrast to the SAP102 intensity tangential gradient, which did not change with age (Supplementary Text, Fig. S17). This shows that these two closely related synaptic proteins undergo distinct spatiotemporal changes within the dendrites of CA1 pyramidal cells, producing a changing two-dimensional synaptome map across the lifespan.

We next asked whether these age-dependent changes in CA1 synaptome architecture produce different functional outputs in response to patterns of neural activity. Using established methods(*18*), we plotted the amplitude of the postsynaptic response (excitatory postsynaptic potential, EPSP) of 11 x 11 synapses from the hippocampal region to trains of eight action potentials (Fig. 4A, C). The response to each action potential and the summed response to all eight action potentials is shown for three patterns of activity – theta burst, gamma and theta train patterns (Fig. 4C). This clearly shows that the functional outputs of the CA1 synaptome maps vary with age and with the pattern of activity. Consistent with the limited capacity to process behavioral information in the very young animal, the spatial responses for all three patterns of activity were less complex at 1W than at adult ages.

Different subregions of the hippocampal formation contribute distinct cognitive functions, which together produce an integrated behavioral output(*26–28*). This integrated function is exemplified by the classical trisynaptic circuit that connects the dentate gyrus (DG) to CA3, which in turn drives the CA1 region(*29*) (Fig. 4D). To examine the possibility that the integrated functions of hippocampal subregions may change with age, we generated synapse subtype lifespan trajectories for all regions comprising the trisynaptic circuit (Fig. 4D) and all hippocampal subregions (Fig. S18). Each subregion in the trisynaptic circuit undergoes a different lifespan trajectory of synaptic subtype composition, indicating that the memory functions controlled by these hippocampal subregions will change with the LSA.

## Discussion

We have systematically mapped the molecular and morphological features of individual excitatory synapses across the brain and throughout the lifespan in the mouse, creating the first lifespan synaptome atlas. A striking finding is that there are continuous changes in the synapse composition of all brain areas across the lifespan. The dynamic temporal trajectories of synapse number, protein composition, morphology, types and subtypes in over 100 brain areas reveal landmarks and phases that define a lifespan synaptome architecture (LSA) that can be divided into three phases (Fig. 5).

**Figure 5.**
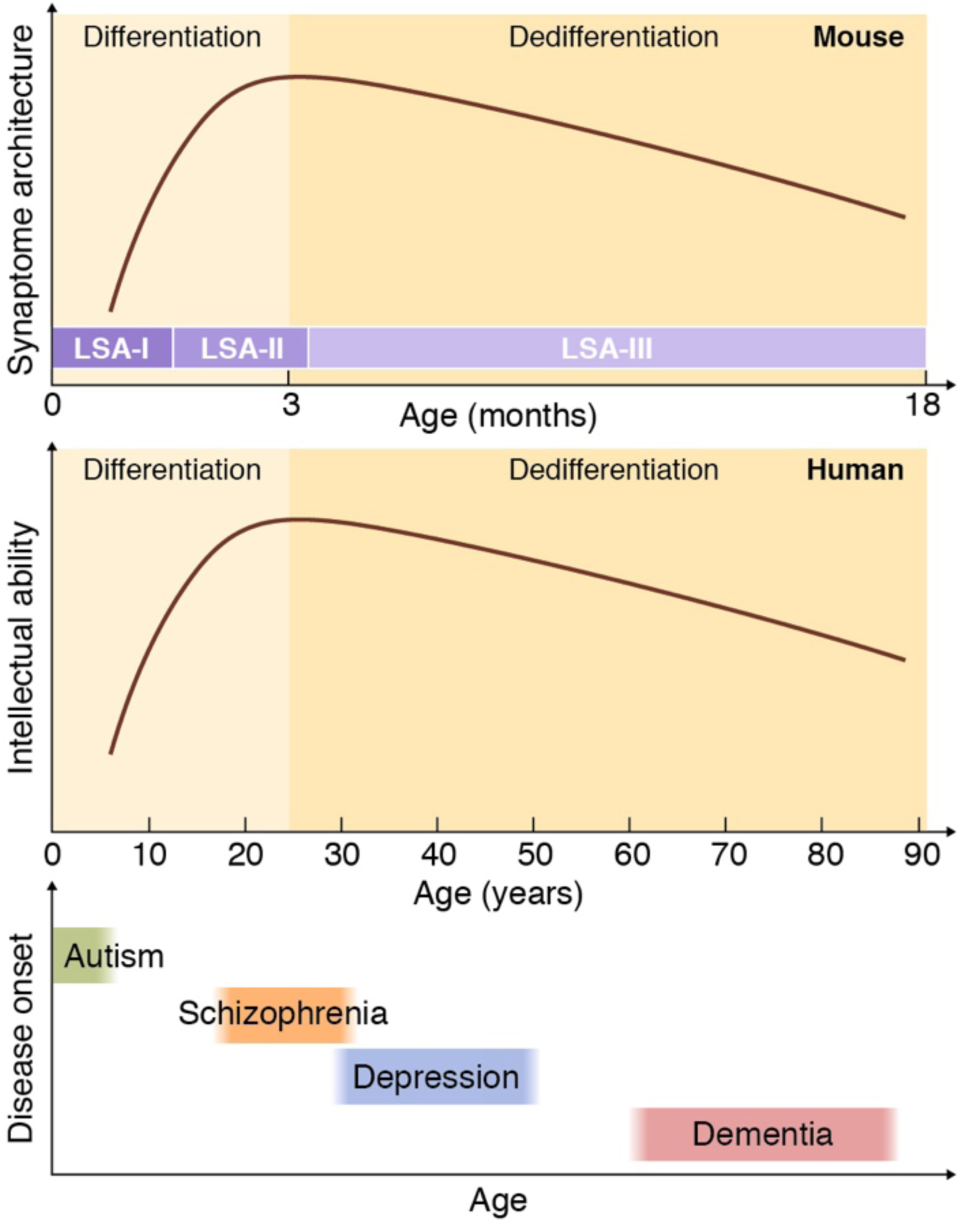
Model of the correspondence of systems level lifespan synaptome architecture, psychology and brain disorders. (Top) The trajectory of LSA in the mouse, showing the differentiation of brain areas peaking in young adults followed by gradual dedifferentiation into old age. The three phases are illustrated, with LSA-I, LSA-II and LSA-III corresponding to childhood, young adulthood and ageing in humans. (Middle) The lifespan trajectory of intellectual abilities (adapted from(*4*)) in humans with ages scaled to mouse in accordance with transcriptome data(*30*). The differentiation, peak age and dedifferentiation of intellectual abilities correlate with the trajectory of synaptome architecture. (Bottom) The age windows of onset of autism, schizophrenia, depression and dementia in humans (*38*). We propose that the programmed changes in brain synapse composition that characterize the LSA render the brain vulnerable to the genetic and pathological mechanisms causing psychiatric and neurodegenerative disorders at different ages.

The first phase (LSA-I) spans from birth to 1M and is characterized by a dramatic increase in synapse number and diversity. This is the most dynamic of all phases and coincides with the animal’s initial dependence on its mother before weaning at 3W. The transition to the second phase (LSA-II) is marked by an abrupt cessation of synapse proliferation in most areas. Despite this, the synapse composition continues to change in all areas, with an increase in diversity especially in higher brain areas. These changes continue to drive the differentiation and specialization of brain areas and the organization of the brain-wide synaptome network architecture to its zenith at 3M. At this age, mice have developed a complete and complex behavioral repertoire, enabling them to respond to diverse environmental challenges. Importantly, brain transcriptome analyses show that 3M in mice corresponds to ∼25 years of age in humans(*30*), which is also the peak for many intellectual abilities and cognitive processes(*4*). The rapid expansion of synapse diversity and changing synapse composition that characterizes the adolescent brain are likely to contribute to the complex behavioral changes occurring during this phase of life(*31*). From this age onward (LSA-III) there is a gradual erosion of the synaptome architecture, which begins with a decline in synaptome network topology and accelerates at 6M with reductions in synapse number, changing synapse parameters, types and subtypes. The net effect is a change in the synapse composition of brain areas that causes them to dedifferentiate from each other. Thus, there is a loss of the specialized functions of brain areas and the global network properties of the brain in old age.

A remarkable finding is that the synaptome architecture of the elderly brain becomes more similar to that of the juvenile brain. Thus, ageing degrades the highly specialized synaptome of the young adult animal to one found in an animal that is dependent on its mother. This may explain the preservation of basal physiological functions (e.g. feeding) relative to the loss of higher cognitive functions in the elderly. We were also struck by the changing populations of synapse subtypes with age, with some subtypes clearly being more resilient to ageing whereas others decline in proportion and may indicate age-dependent vulnerability. The general increase in synapse size with age could be accounted for by the decrease in relative abundance of small synapses, which is consistent with electron microscopy studies in the ageing macaque dorsolateral prefrontal cortex(*32–34*). With these new findings in mind, it should be possible to identify genes that enhance or slow synaptome ageing and potentially develop interventions that may slow intellectual decline in old age.

We have shown that the LSA impacts on synaptic physiological mechanisms within brain subregions, between subregions in the hippocampus, and between regions on the global systems level. The changes in the capacity of the hippocampal region to generate representations in response to neural activity and the differing trajectories of hippocampal subregion synapse composition could account for changes in memory, spatial navigation(*35, 36*) and cognitive functions that occur with age(*37*). The progression of regional differentiation and the increase in synaptome network smallworldness during LSA-I and LSA-II are consistent with brain imaging studies showing that the integrated roles of multiple areas contribute to the development of language, social behaviors, emotion, empathy, anxiety, moral judgement and other attributes from birth to young adulthood in humans(*8*). Furthermore, the time course of the arching lifespan trajectory of the LSA corresponds well with the arching lifespan trajectory of intellectual ability(*4–7*) and the integrated functions of the brain that contribute to measures of intelligence(*7*) (Fig. 5).

An extensive literature describes the relationships between the acquisition of behaviors in childhood, adolescence and young adults (corresponding to LSA-I and LSA-II) and the disorders that manifest as pathological behaviors(*8*) (Fig. 5). These include autism, language impairments, social anxieties, psychopathy and schizophrenia, among others(*8*). At later adult ages that correspond to LSA-III, depression and dementia are increasingly common. Importantly, it is known that there are mutations and variants that directly target synaptic proteins or cellular mechanisms that damage synapses in many of these disorders(*10*). Yet, one of the most important and unanswered questions is why the pathology of these disorders targets particular brain areas and ages. Our findings indicate that the changing molecular composition of synapses will render them vulnerable to genetic disorders at different ages. Our results also predict that altering gene expression will shift the distributions of synapse types/subtypes in particular circuits at different ages, opening windows of vulnerability and producing behavioral phenotypes.

The LSA has the hallmarks of a genetically regulated biological program. Although we expect numerous cell biological, developmental and ageing mechanisms to play a role in the phases of the LSA, a key mechanism that could explain the regulation of synapse proteins and synaptome architecture across the whole lifespan is the Genetic Lifespan Calendar(*30*). This program regulates synaptic gene expression across all ages and, moreover, produces a crucial turning point in the expression of synaptic proteins that overlaps with the turning point at the zenith of the LSA. Our highly scalable synaptomic methods and the Lifespan Synaptome Atlas delivered here provide valuable new tools for studying genetic models of diseases and their impact on synaptome architecture throughout development and aging.

## Acknowledgements

C. McLaughlin and K. Elsegood for mouse colony and lab management, D. Kerrigan and D. Fricker for genotyping; N. G. Skene for statistical advice; S. Munni, O. Kealy, H. Taczynski for image calibration; D. Maizels for artwork; C. Davey for editing.

## Author Contributions

MC, design of animal cohort; collection, preparation and imaging of brain samples; delineation of brain regions/subregions; calibration of synapse detection; analysis of synapse parameters; analysis of hippocampal gradients; data interpretation; ZQ, methodology development and optimization of lifespan SYNMAP pipeline; data analysis of the whole lifespan mouse cohort: image segmentation and puncta quantification, classification, unsupervised and supervised mapping of synapse parameters, types and subtypes, diversity and network topology analysis; EF, statistical analysis and computational modelling of synaptome physiology; BK, construction of Synaptome Explorer and Synaptome Homology Viewer; RG, construction of website; NHK, advice and supervision; SGNG, conception, supervision and writing.

## Data and materials availability

all data are available at the Mouse Synaptome Atlas (www.brain-synaptome.org).

## List of Supplementary Materials

**Materials and Methods**

**Figures S1-S18**

**Table** S1

**External Database** https://doi.org/10.7488/ds/2711

**References** (38-44)

## Other supplementary Materials for this manuscript include the following

Mouse Lifespan Synaptome Atlas (www.brain-synaptome.org)

Synaptome Explorer V2 (www.brain-synaptome.org)

Synaptome Homology Viewer (www.brain-synaptome.org)

External Database: https://doi.org/10.7488/ds/2711

## Materials and Methods

### Animals

Animal procedures were performed in accordance with UK Home Office regulations and approved by Edinburgh University Director of Biological Services. Generation and characterization of PSD95^eGFP/eGFP^/SAP102^mKO2/mKO2^ knock-in mouse line was described previously(*18*). Para-sagittal brain sections from cohorts of 49 PSD95^eGFP/+^;SAP102^mKO2/y^ male mice were collected at ten postnatal ages: one day (1D, N=5), one week (1W, N=5), two weeks (2W, N=5), three weeks (3W, N=4), one month (1M, N=6), two months (2M, N=6), three months (3M, N=5), six months (6M, N=5), 12 months (12M, N=3) and 18 months (18M, N=5). From 12 months of age, mice were monitored monthly by weight and scoring their body condition as previously described (*39*). No change in mouse weight or body condition was observed in ageing mice.

### Tissue collection

Tissue collection was performed at ten ages as described previously(*18*). Briefly, mice were anesthetized by intraperitoneal injection of 20% pentobarbital sodium (Pentoject, Animalcare Ltd.; 0.03 ml for 1D, 0.05 ml for 1W-1M, and 0.1 ml for 2M-18M animals). Upon complete anesthesia, transcardiac perfusion of phosphate buffered saline (PBS; Oxoid) was performed and followed by transcardiac perfusion of 4% (v/v) paraformaldehyde (PFA; Alfa Aesar). PBS and PFA volumes used were: 2 ml (1D), 3 ml (1W), 5 ml (2W), 7ml (3W), 8 ml (1M) and 10 ml (2M-18M). Whole brain was then dissected and post-fixed in 4% PFA for 1h (1D), 1.5h (1W), 2h (2W), 2.5h (3W), 3h (1M) or 3.5h (2M-18M), followed by cryoprotection in 30% sucrose solution (w/v in PBS; VWR Chemicals) for 24h (1D-1W), 48h (2W-3W) or 72h (1M-18M). Brains were then placed in a cryomold, embedded in optimal cutting temperature (OCT, CellPath) and frozen in isopentane cooled with liquid nitrogen. Para-sagittal brain sections were then cut at 18 μm thickness using a Thermo Fisher NX70 cryostat. Cryosections were placed on a Superfrost Plus glass slides (Thermo Scientific) and stored at −80°C.

### Tissue preparation

Para-sagittal sections from left hemisphere (1.2 mm laterally from the midline in Franklin and Paxinos sagittal atlas, corresponding to sections 12-13/24 from sagittal Allen Brain Reference Atlas) were washed for 5 min in PBS, incubated for 15 min in 1 mg/ml DAPI (Sigma), washed with PBS, mounted in home-made MOWIOL (Calbiochem) containing 2.5% anti-fading agent DABCO

(Sigma-Aldrich), covered with a coverslip (thickness #1.5, VWR international) and imaged the following day.

### Slide scanner widefield microscopy

Whole sections were imaged using the Zeiss Axio Scan.Z1 system with a Zeiss Plan-Apochromat 20X lens with a numerical aperture (NA) of 0.8. Samples were illuminated using Colibri.2 at 365 nm for DAPI, 470 nm for EGFP and 555 nm for mKO2. Excitation filters were 365 nm for DAPI, 470/40 nm for EGFP and 546/12 nm for mKO2. Emission filters were 445/50 nm for DAPI, 525/50 nm for eGFP and 607/80 nm for mKO2. Light detection was achieved using a Hamamatsu Orca-flash 4.0 monochrome camera.

### Spinning disc confocal microscopy

Fast high-resolution imaging was achieved using the Andor Revolution XDi system, equipped with an Olympus UPlanSAPO 100X oil immersion lens (NA 1.4), a CSU-X1 spinning-disc (Yokogawa), an Andor iXon Ultra monochrome back-illuminated EMCCD camera, a 2X post-magnification lens and a Borealis Perfect Illumination Delivery™. Images have a pixel dimension of 84 x 84 nm and a depth of 16 bits. A single mosaic grid, with no overlap between adjacent tiles, was set up in the Andor iQ2 software to cover each entire brain section, using an adaptive *z* focus to follow the unevenness of the tissue. eGFP was excited using a 488 nm laser and mKO2 with a 561 nm laser. Emitted light was filtered with a Quad filter (BP 440/40, BP 521/21, BP 607/34 and BP 700/45).

### Acquisition parameters in developmental and adult time points

To optimize imaging between time points where synapse intensity was low (development) or high (adulthood), two sets of spinning disc acquisition parameters were used for 1D-3M and 3M-12M time points. Acquisition of the 3M time point was performed with both sets of parameters using adjacent brain sections from the same mice (N=3) and synapse parameters were compared. As no significant difference was found between the two sets of data, no correction was applied and the two sets of time points were merged.

### Detection of synaptic puncta

Synaptic puncta were detected using the developed machine learning-based Ensemble detection method (*18*). A training set of the 552 images (10.8 × 10.8 µm) for PSD95 and SAP102 (1104 images in total), respectively, across various mouse brain regions was randomly sampled using bootstrapping(*40*) from a data cohort of different age groups. Synaptic puncta were manually annotated by their centroid location in the images by four experts independently. A weighting factor (0–1) was then given as the measurement of the annotation quality for each human expert. The ground truth was finally generated by aggregating the weighting factors of the four annotations: puncta annotated with an average weight greater than 0.7 would be considered as true puncta.

To avoid any overfitting problem, a K-fold cross-validation strategy was adopted to train the puncta detector using ensemble learning as described previously (*18*): half of the training set is randomly selected to train the detector and the other half to validate and test the performance of the detector. This procedure is repeated 150 times, each using a different random division of the training set for cross validation. An overall 95% detection rate for PSD95 and 92% for SAP102 was achieved on the validation dataset, both of which outperform conventional puncta/particle detection algorithms(*41*) on fluorescence microscopy images. The detector after training is finally applied to other images for puncta detection.

### Measurement of synaptic parameters

Once detected and localized, all puncta were segmented by thresholding their intensities adaptively: a threshold for each punctum is set as 10% of the height of the intensity profile. Six punctum parameters were then quantified: mean pixel intensity, size, skewness, kurtosis, circularity and aspect ratio, the latter four of which were used for shape quantification. For details, see previous work (*18*).

### Classification of synaptic puncta

Puncta were classified in 3 type classes depending on their expression of either or both synaptic protein: type 1 for PSD95 only, type 2 for SAP102 only and type 3 for colocalized PSD95+SAP102(*18*). Using the synapse subtype catalogues built from previous work(*18*), we can further classify each punctum detected and quantified from the last two steps into one of the 37 subtypes from the catalogue or some other new subtypes: Bayesian classifiers are first built for each of all subtypes. For each punctum, the likelihood values (0–1) of all 37 subtypes are calculated based on the punctum parameters and the 37 Bayesian classifiers. Finally, the punctum is labelled as the subtype with highest likelihood value, if the value is higher than a given threshold (set as 0.5). Otherwise the punctum is labelled as a new ‘other subtypes’. We find that overall the ‘other subtypes’ only accounts for ∼0.002% of the whole population of individual synapses quantified from the whole data cohort, and is thus considered not to affect the composition and diversity of synapses across the lifespan.

### Measurement of synapse diversity

Synapse diversity is measured based on the Shannon entropy to quantify the population differences across 37 subtypes in a given unit brain area: 0 if all puncta belong to a single subtype and 1 if all subtypes occur equally in the population. For details see previous work(*18*).

### Segmentation of brain regions and subregions

After high-resolution imaging, a downsized stitched image of the whole section was generated and used for manual delineations of brain subregions in ImageJ, according to the online reference atlas of the Allen Mouse Brain Atlas (http://mouse.brain-map.org). Delineation masks from these delineations were then used to generate mean values of synaptic punctum parameters over the corresponding subregion. For region analysis, the corresponding mask was generated by combining all the masks from subregions that belong to this brain region. For 1D mice, region masks were delineated directly as no subregion masks were generated. For estimation of the brain size, the area covered by the masks resulting from all main regions (whole section area) was measured at each time point.

### Similarity matrices, hypersimilarity matrix and network analysis

Each row/column in the similarity and hypersimilarity matrix represents one delineated brain subregion at one age (similarity matrix) or different ages (hypersimilarity matrix). Elements in the matrix are the synaptome similarities between two subregions quantified by differences in standardized synaptome parameters, details of which can be found in previous work(*18*).

The network analysis in Fig. 3B, C was based on the similarity matrices of individual brain sections quantified in a similar way to those in Fig. 3A and S13. Nodes in the network are representations of the delineated subregion. The small worldness is the topology quantification of the network where the whole set of nodes are divided into small and clustered groups: nodes within same groups are highly connected/similar, whereas those between groups are disconnected/dissimilar(*42*). For a given network the small worldness is a function of network density and is usually plotted as a curve against the network density(*18*). To facilitate comparison of small worldness between different age groups, we convert the small-worldness curve of each brain section into a single-value small worldness by averaging the curve between 0% and 30% network densities. We refer to this single-value measurement as the average small worldness, which allows us to present the small worldness of age groups using scatter plots in Fig. 3C and perform the hypothesis test.

Within-region similarity is defined as the average similarities between any two subregions belonging to the same main overarching brain regions. This corresponds to areas marked by the 12 small white boxes that are distributed diagonally in the similarity matrix (Fig. 3A and S13). In contrast, the between-region similarity is the average similarities between any two subregions coming from different overarching brain regions. This is the areas in the similarity matrix that are not bound by the 12 white boxes lying on the diagonal. The similarity ratio at each of the ages in Fig. 3B is then calculated as the between-region similarity divided by the within-region similarity.

### Hippocampal gradient analysis

To analyse the lifespan trajectories of intraregional synapse diversity, we focused on hippocampal CA1sr gradients. For mice aged 1W-18M, the CA1sr subregion was subdivided in two directions: four sublayers of equal thickness were delineated in the radial direction, starting from deep (close to CA1 stratum pyramidale) to superficial CA1sr (close to CA1 stratum lacunosum-moleculare); and 10 subfields of equal width were delineated in the tangential direction, from proximal (close to the CA2 field) to distal (close to the subiculum, SUB). Average synaptic parameters were measured in these different sublayers/subfields. Pearson coefficients were calculated between parameter values in a given sublayer/subfield and its sublayer/subfield level (i.e. 1 to 4 and 1 to 10, respectively). The slope value was calculated as the slope of the linear regression curve between parameter values in a given sublayer/subfield and its sublayer/subfield level (i.e. 1 to 4 and 1 to 10, respectively).

### Computational modelling of synaptic responses

Computational modeling of synaptic responses was based on our previously described model(*18*) representing physiology at 3M of age. To model synaptic physiology corresponding to 1W and 18M, differences in synaptome gradients between 3M and 1W and 18M, respectively, were estimated from PSD95 and SAP102 size data obtained in this study. More specifically, for each protein a size gradient was estimated in the tangential direction from the four regions CA1, CA2, CA3 and dentate gyrus and in the radial direction from the layers stratum oriens, pyramidale, radiatum and lacunosum moleculare of CA1. Differences in gradients were used to scale the amplitude of short-term depression and facilitation (PSD95 and SAP102 tangential gradient decrement factor, respectively), as well as time constant of short-term depression and facilitation (PSD95 and SAP102 radial gradient decrement factor, respectively).

### Lifespan Synaptome Explorer and Homology Viewer

To visualize the generated datasets interactively, we developed the tools Synaptome Explorer v2 and Homology Viewer. Synaptome Explorer v2 is an improved version of existing software(*18*) that is used for in-depth exploration of the synaptome of a single brain section. It enables interactive visualization of the brain region in full resolution, as captured by the microscope, and uses overlays to display the synaptic puncta and all their parameters, providing users with extensive parameter range filters to display subsets of puncta accordingly. The granularity of the data visualization is at the level of individual puncta, as users can click on a single punctum and see its parameters. An additional feature is region-based and tile-based filtered puncta statistics, where users can set up parameter range filters, select a combination of regions or subregions and interactively calculate statistics, such as mean intensity, mean size and density, visualized over the corresponding regions or tiles, respectively.

Homology Viewer is a tool that enables interactive visualization of similarity matrices whose values correspond to brain regions/subregions, puts the data in a spatial context and allows simultaneous visualization of different brain section data that are delineated similarly. Users can hover over a brain region or a similarity matrix entry and immediately see how that region is similar to all other regions using a heatmap. Likewise, when using multiple brain regions, users can visualize how a hovered-over region in a brain section is similar to all other regions in all other brain sections. Thus, the tool makes the results of similarity matrices visually accessible owing to the direct mapping of the data on the brain section(s).

## Supplementary text

### Distinct lifespan trajectories of radial and tangential hippocampal gradients

To study the lifespan synaptome trajectory of intraregional diversity, we focused on the CA1 stratum radiatum (CA1sr), which exhibits gradients for PSD95 intensity in radial and tangential directions and for SAP102 intensity in the tangential direction(*18*). The CA1sr subregion was subdivided into four sublayers in the radial direction from deep (close to CA1sp) to superficial CA1sr (close to CA1slm; Fig. S17A) and in 10 subfields in the tangential direction from proximal (close to CA2) to distal CA1sr (close to subiculum; Fig. S17B) for mice from 1W to 18M.

For PSD95 intensity radial gradient (Fig. S17C, F and I), we observe no gradient during the first 3 weeks (Pearson ≍ 0 from 1W-2W; Fig. S17F). At 1M, a weak gradient appears (Pearson = 0.55) and this gradient is fully established by 2M and maintained during all mouse adult life (Pearson > 0.8 from 2M-18M). The slope on the PSD95 intensity gradient follows a similar trajectory (Slope ≍ 0 from 1W-2W, Slope = 235 at 1M, Slope > 350 from 2M-18M; Fig. S17I). By contrast, PSD95 intensity shows a tangential gradient as early as 1W (Pearson = 0.91; Fig. S17D, G and J), which is maintained throughout the lifespan (Pearson > 0.5 from 1W-18M). Interestingly, the slope of this gradient has a different trajectory: it starts low at 1W and progressively increases, peaks at 3M and then gradually decreases 3M-18M (Fig. S17J). SAP102 intensity already forms a tangential gradient at 1W and remains relatively constant throughout life for both Pearson coefficient and slope value (Fig. S17E, H and K).

Thus, we show three types of gradient: a PSD95 intensity radial gradient is established between 3W and 2M and maintained throughout adulthood and ageing; a PSD95 intensity tangential gradient is present as early as 1W but the slope increases during development and then gradually plateaus during adulthood; and a SAP102 intensity tangential gradient is already established at 1W and is maintained throughout life. In adults, tangential gradients mostly derive from expression gradients between adjacent pyramidal neurons along CA1 (between-cell gradient(*43*)). We show here that tangential synaptic gradients are established very early in development, before 1W. By contrast, radial gradients represent synapse diversity arising from synapses within the dendritic tree of pyramidal cells (within-cell gradient(*18, 44*)), which receive layers of connections from different sources. Thus, the later development of a PSD95 intensity gradient during postnatal development (between 3W and 2M) may indicate that it is shaped during brain maturation by connectivity and activity, as opposed to the tangential gradient that could result from intrinsic, genetically programmed differences between pyramidal cells. Moreover, the different characteristics of the PSD95 and SAP102 tangential gradients (e.g. the inverted U shape of the slope of PSD95 intensity (Fig. S17J) compared with the steady slope of SAP102 intensity (Fig. S17K)) indicates that gradients vary between synaptic proteins across the lifespan. Considering that there are hundreds of other synaptic proteins that may also show differential gradients with age, then the synaptome map of dendrites and brain areas could show exquisite diversity and patterning across the lifespan.

**Fig. S1:**
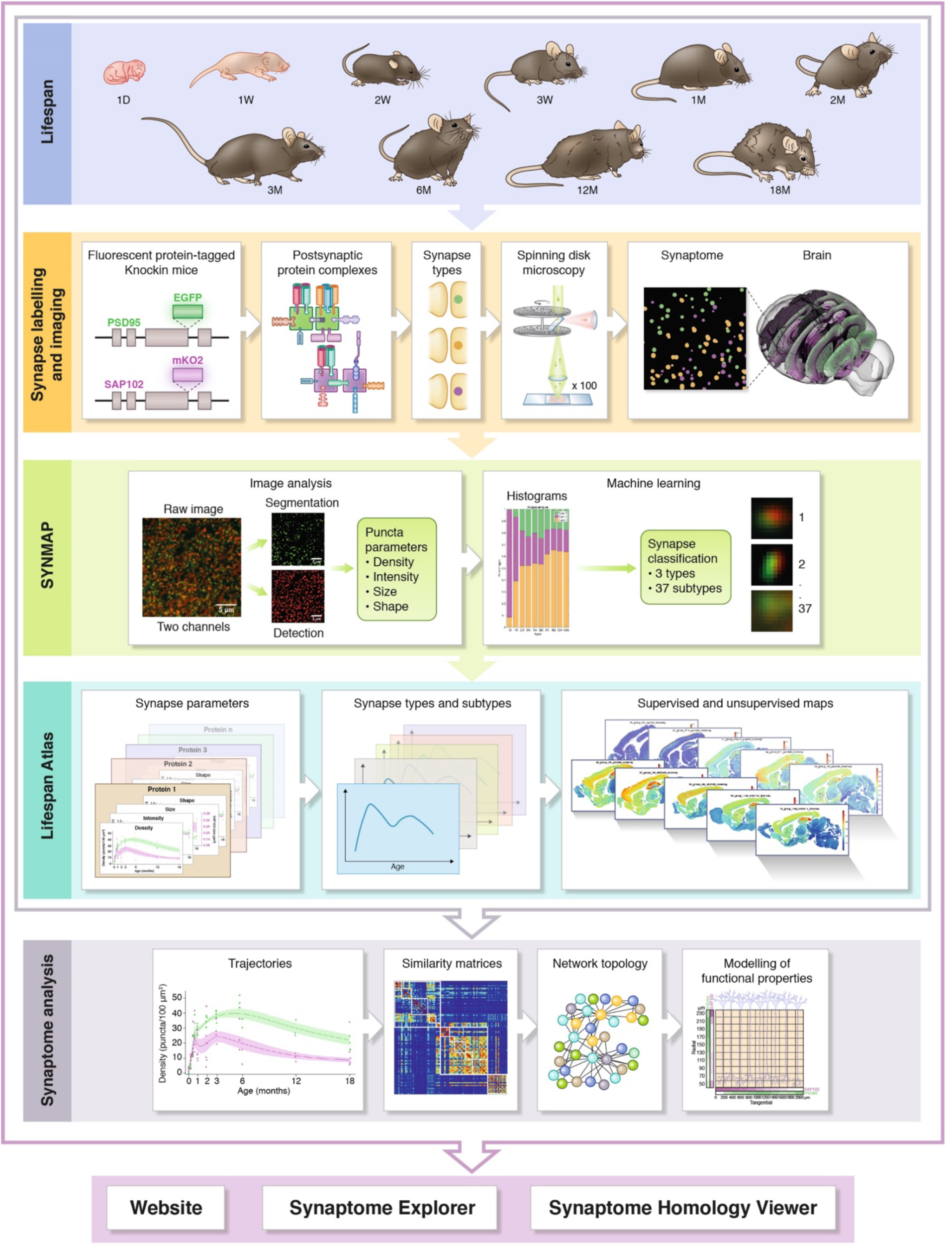
The Mouse Lifespan Synaptome Architecture project. Lifespan, indicates the ten ages from birth to 18M; Synapse labelling and imaging, shows genetic modification of PSD95 with eGFP and SAP102 with mKO2, which labels the proteins and their respective multiprotein complexes which are distributed into synapse types/subtypes that can be visualized in brain sections using confocal spinning disc microscopy; SYNMAP, image analysis pipeline that detects, segments and quantifies synapse puncta, which are categorized into types and subtypes by machine learning; Lifespan Atlas, describes the trajectories of synapse parameters, types and subtypes and houses supervised and unsupervised maps; Synaptome analysis, graphs and heatmaps of trajectories, similarity and hypersimilarity matrices of brain regions, synaptome network topology and modelling of physiological properties. The outputs are disseminated on the Mouse Synaptome Atlas website, Synaptome Explorer and Synaptome Homology Viewer.

**Fig. S2:**
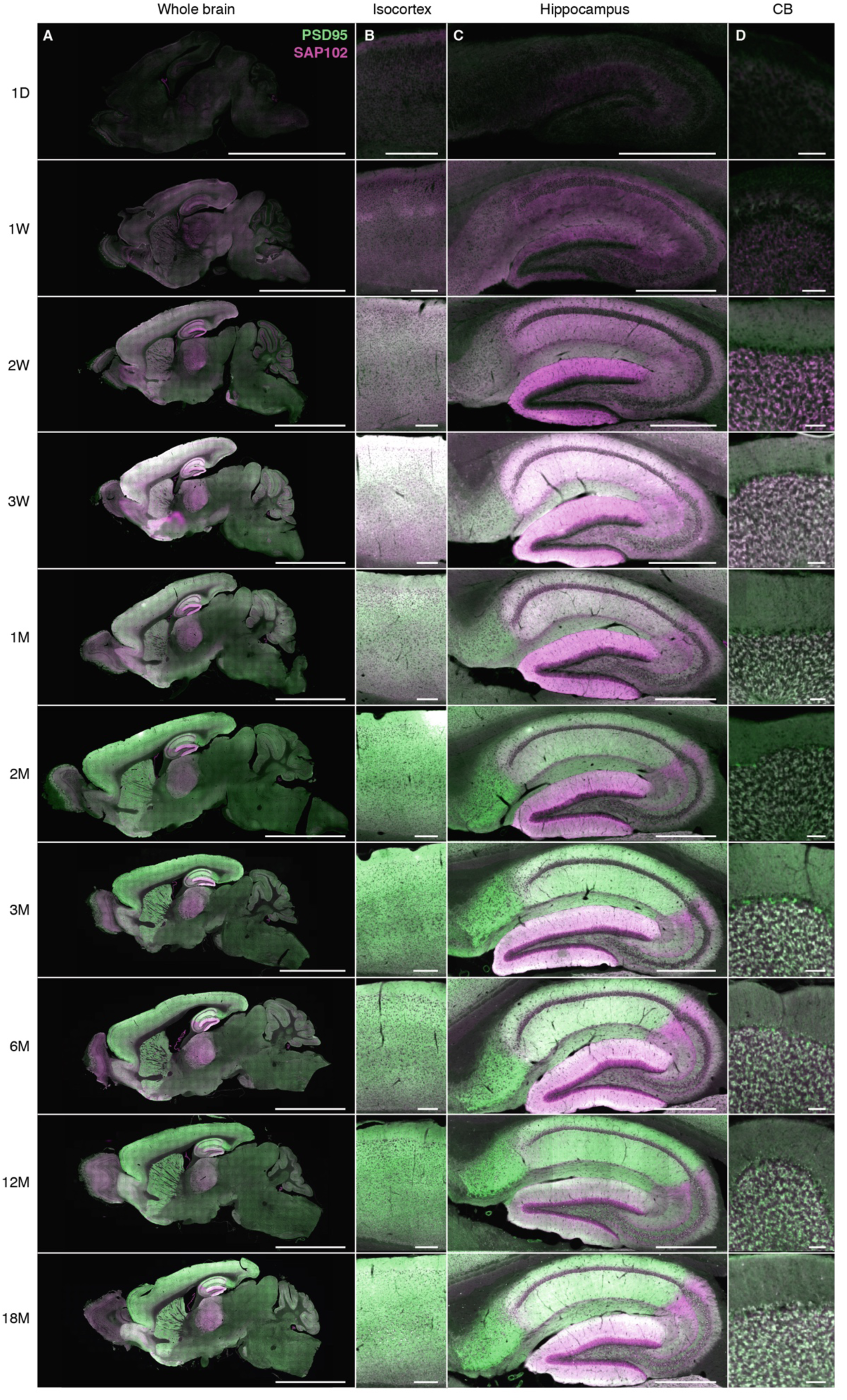
Lifespan expression of PSD95-eGFP and SAP102-mKO2 in the mouse brain. Low-magnification images (20X) showing expression of PSD95-eGFP (green) and SAP102-mKO2 (magenta) in the whole brain (A), isocortex (B), hippocampus (C) and cerebellum (CB) (D) at 1D, 1W, 2W, 3W, 1M, 2M, 3M, 6M, 12M and 18M. Scale bars: 4 mm (A); 200 µm (B); 500 µm (C); 50 µm (D).

**Fig. S3:**
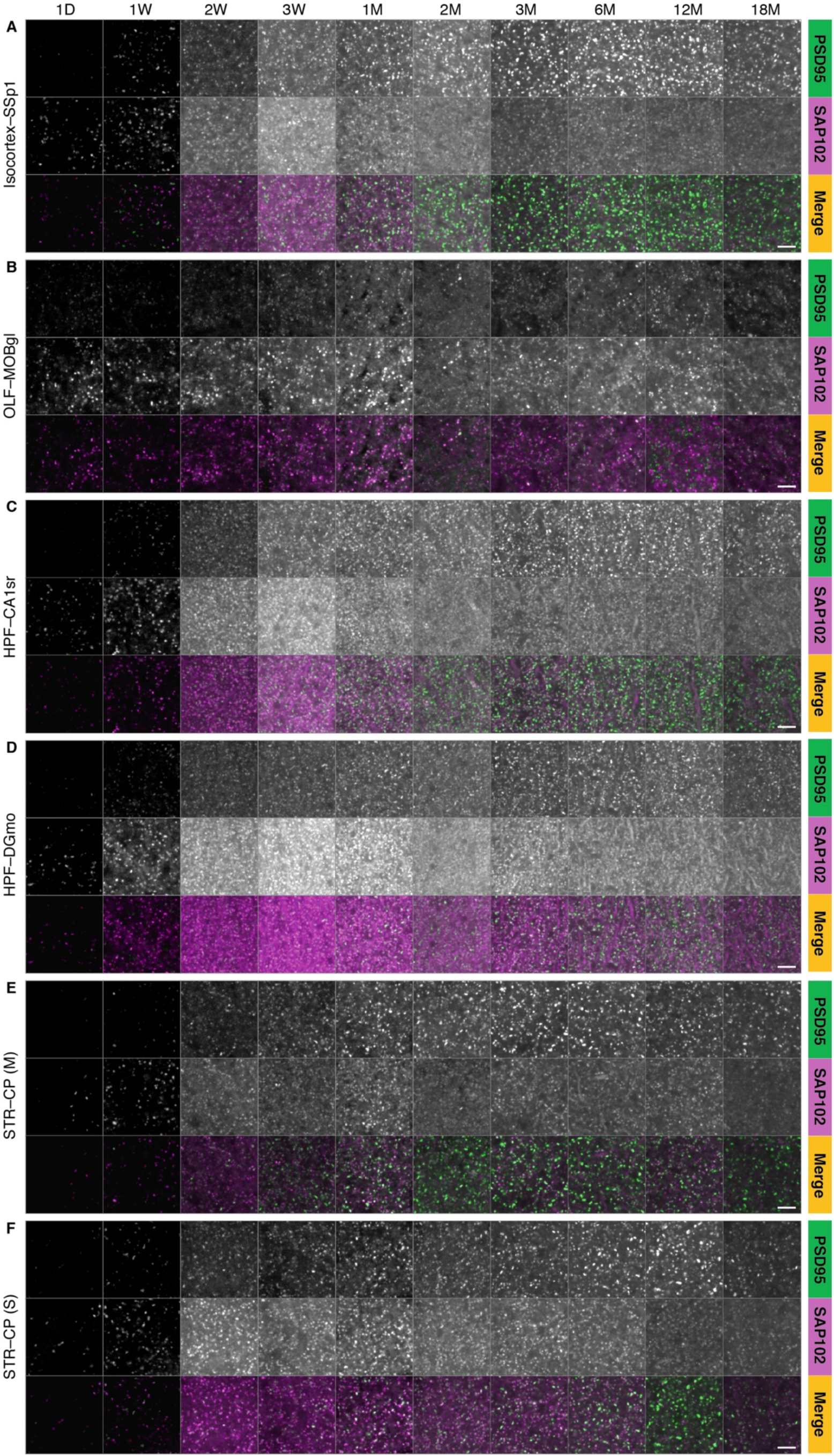
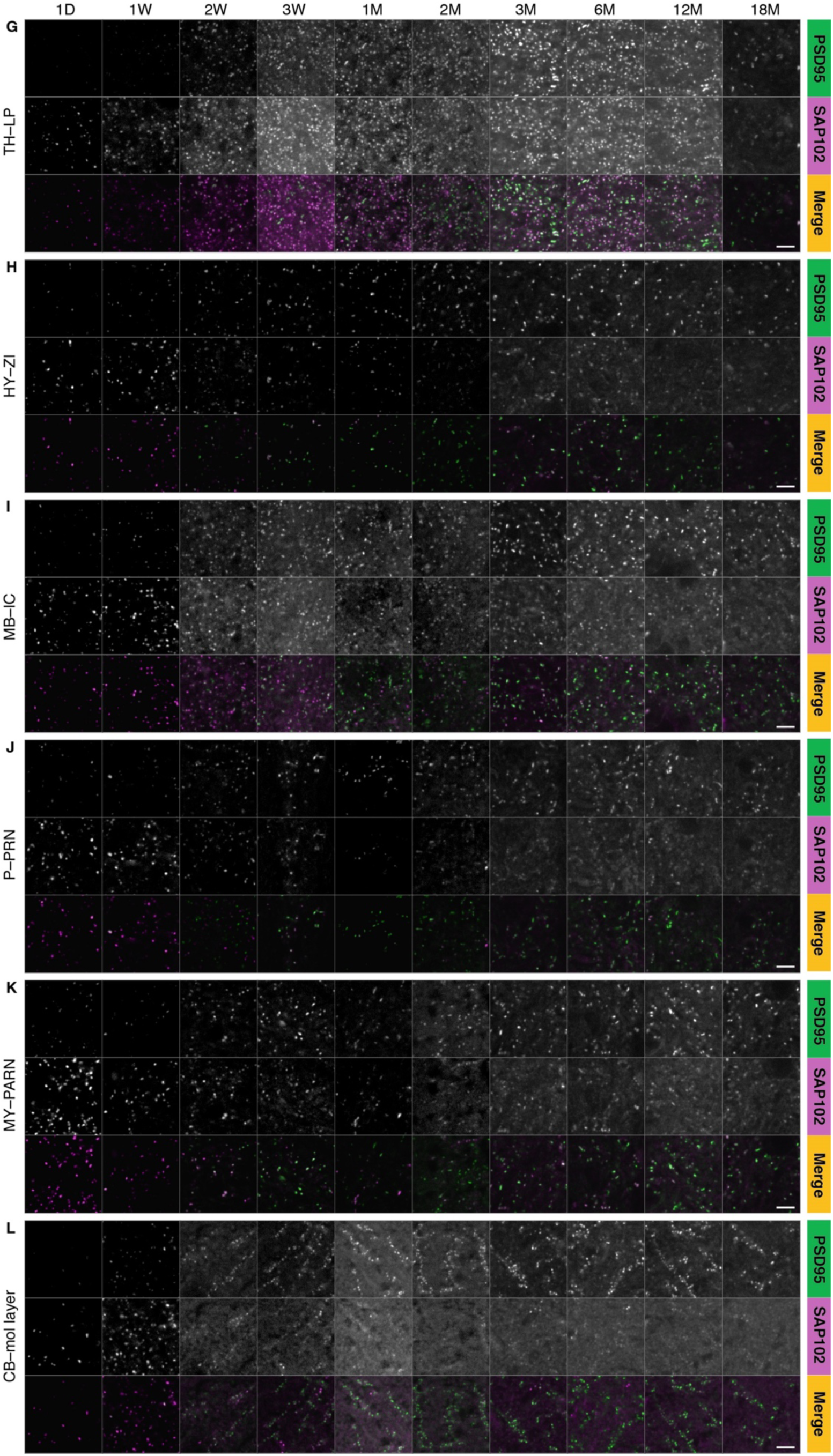
High-resolution images of PSD95-eGFP and SAP102-mKO2 expression across the mouse lifespan. High-resolution images (100X) showing expression of PSD95-eGFP (green, top panels), SAP102-mKO2 (magenta, middle panels) and merge (bottom panels) at the indicated time points in the following brain regions: isocortex, primary somatosensory area (SSp1) (A); OLF, glomerular layer of the main olfactory bulb (MOBgl) (B); HPF, stratum radiatum of CA1 field (CA1sr) (C) and dentate gyrus molecular layer (DGmo) (D); STR, caudate putamen (CP): matrix compartment (E) and striosomes (F); TH, lateral posterior nucleus (LP) (G); HY, zona incerta (ZI) (H); MB, inferior colliculus (IC) (I); P, pontine reticular nucleus (PRN) (J); MY, parvicellular reticular nucleus (PARN) (K); CB, molecular (mol) layer (L). Scale bars: 3.5 µm.

**Fig. S4:**
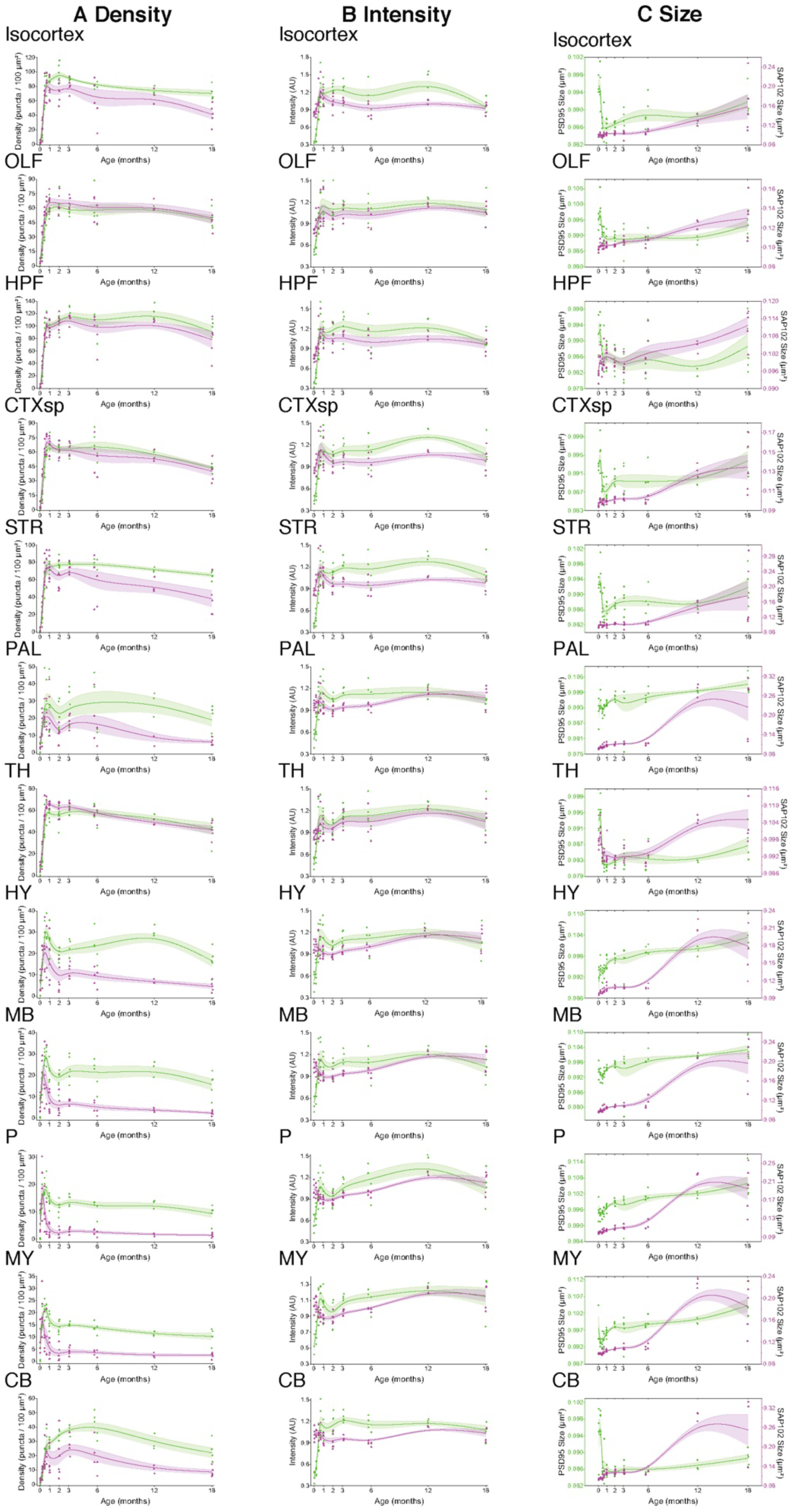
Lifespan trajectories of PSD95 and SAP102 synapse parameters in brain regions. Lifespan trajectories of PSD95 (green) and SAP102 (magenta) synapse density (A), intensity (B) and size (C) from 1D to 18M in 12 main brain regions. Scatter points indicate the mean value of the parameter for individual mice, lines are the β-spline curves of mean values across mice (1D, N=5; 1W, N=5; 2W, N=5; 3W, N=4; 1M, N=6; 2M, N=6; 3M, N=5; 6M, N=5; 12M, N=3; 18M, N=5) and grayed areas represent the standard error of the mean.

**Fig. S5:**
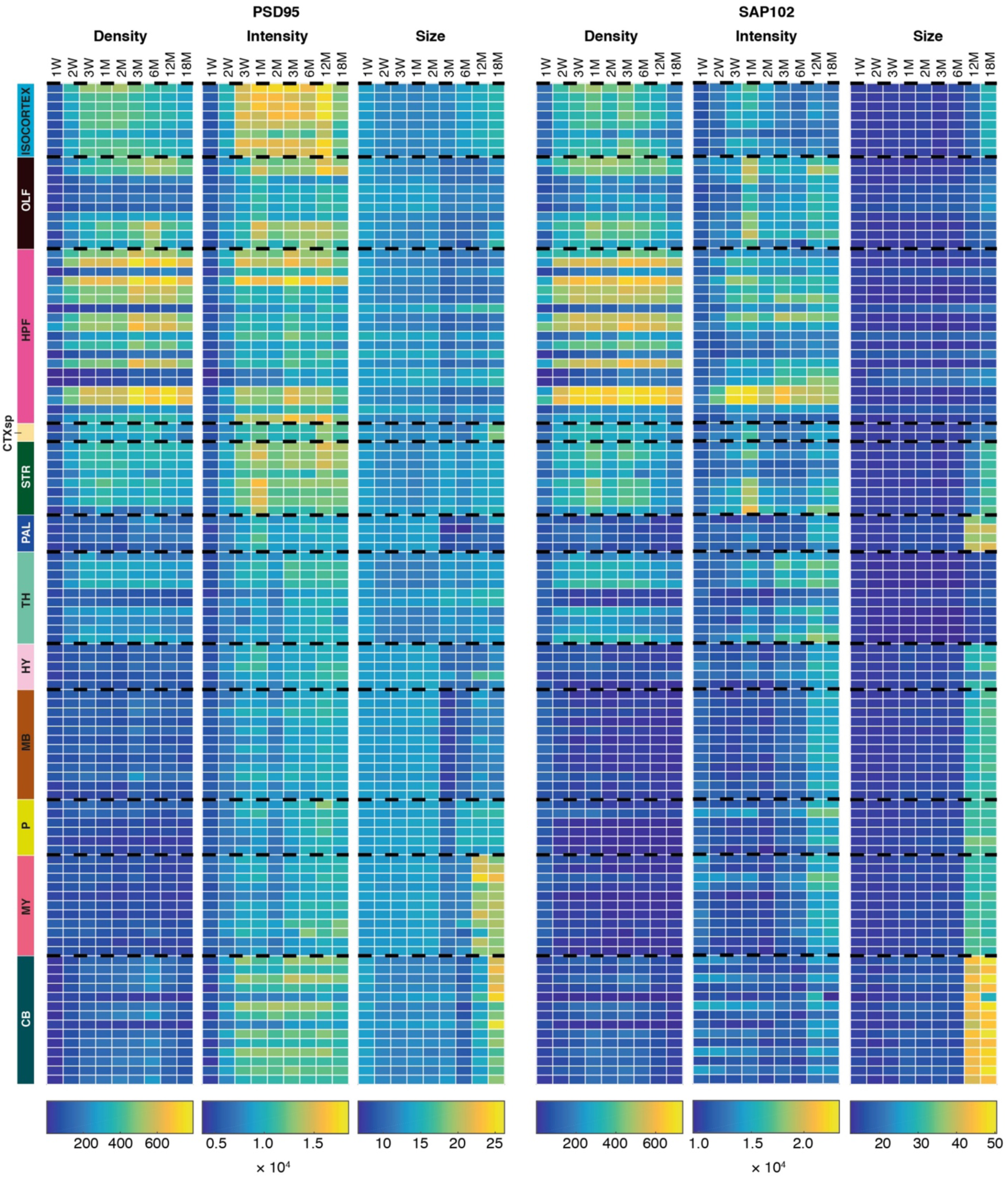
Heatmaps of lifespan trajectories of PSD95 and SAP102 synapse parameters in brain subregions. Lifespan trajectories of raw (mean) values of density, intensity and size of PSD95 and SAP102 puncta in brain subregions (see Table S1 for subregion names). Scale: puncta number per 21.5 × 21.5µm unit area for density, 16-bit grayscale values for intensity, pixel number for size.

**Figure S6.**
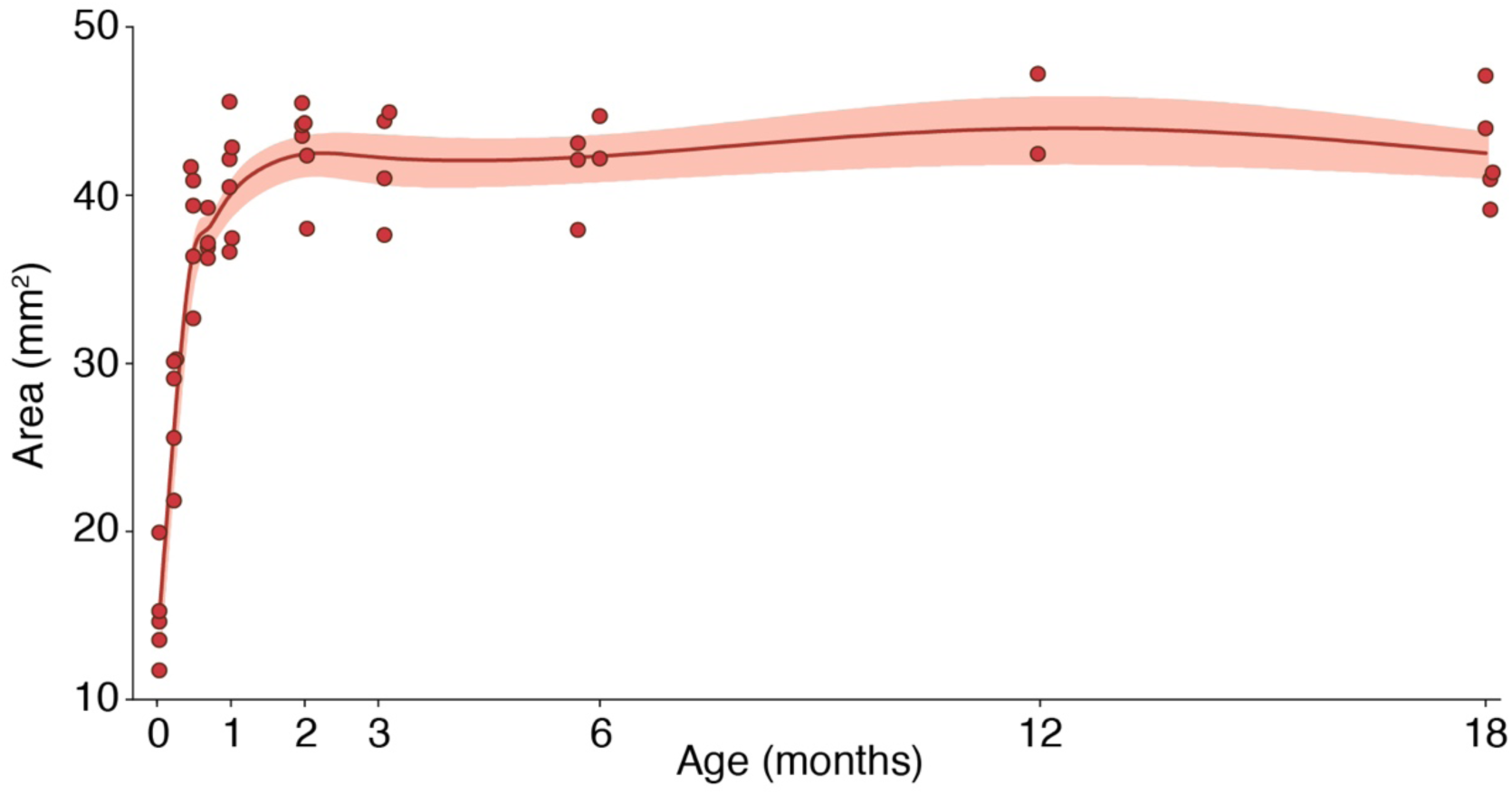
Trajectory of brain size across the lifespan. Area of the whole brain section was measured at each time point (1D, N=5; 1W, N=5; 2W, N=5; 3W, N=4; 1M, N=6; 2M, N=6; 3M, N=4; 6M, N=5; 12M, N=2; 18M, N=5). Scatter points indicate the mean value for individual mice, lines are the β-spline curves of mean values across mice and grayed areas represent the standard error of the mean. Brain size does not significantly vary between 3M (41.98 ± 1.69 mm^2^) and 18M (42.50 ± 1.39 mm^2^) (Cohen’s d = 0.29; t-test p = 0.82).

**Fig. S7:**
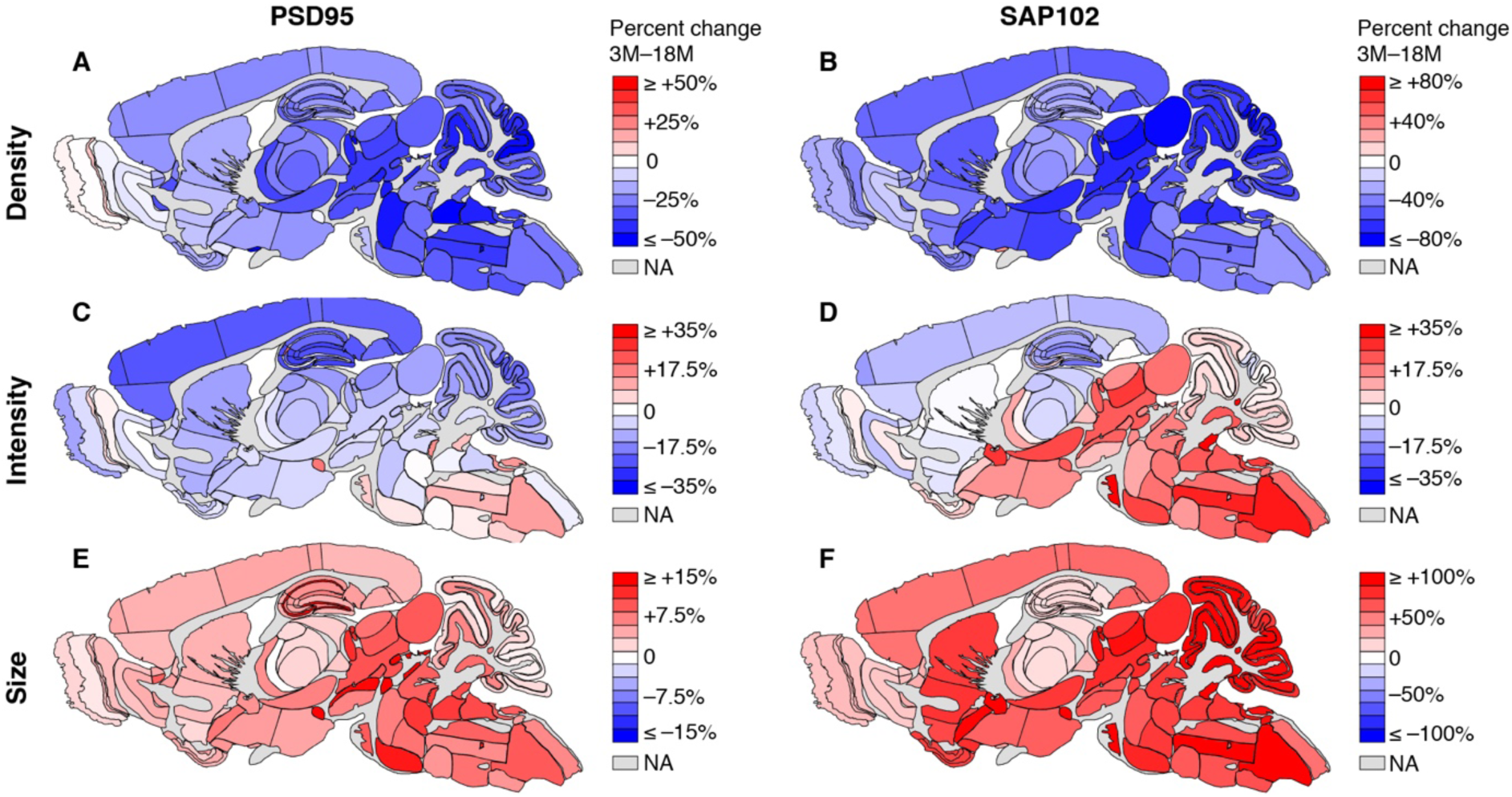
Spatial maps of synapse changes between 3M and 18M. Percentage change between 3M and 18M is color coded in each subregion for PSD95 density (A), SAP102 density (B), PSD95 intensity (C), SAP102 intensity (D), PSD95 size (E) and SAP102 size (F). Increase is indicated in red and decrease in blue.

**Fig. S8:**
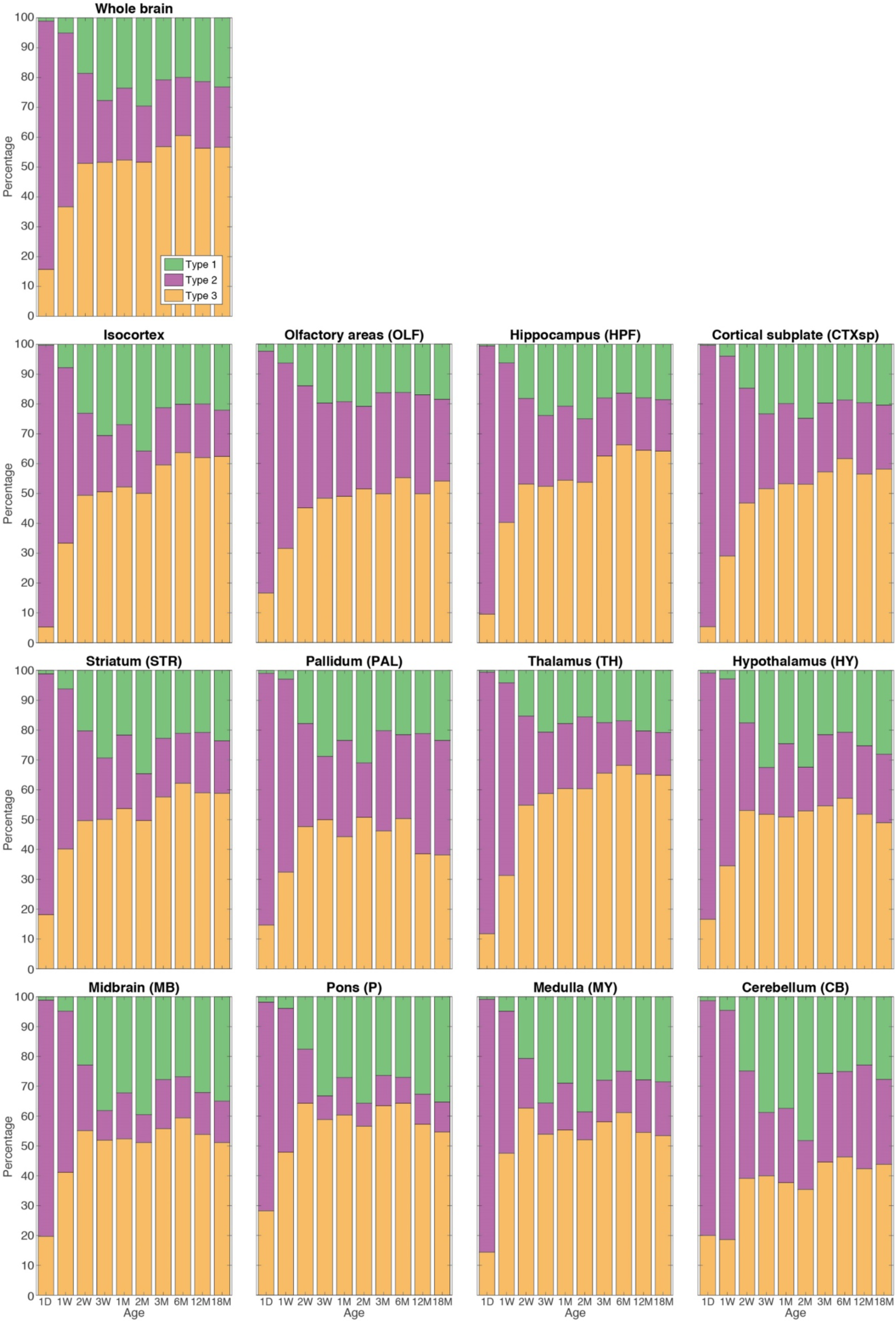
Relative abundance of synapse types in whole brain and brain regions. Stacked bar plots of the percentage of type 1 (PSD95 only), type 2 (SAP102 only) and type 3 (PSD95+SAP102) synapses in the whole brain and 12 main regions across the lifespan.

**Fig. S9:**
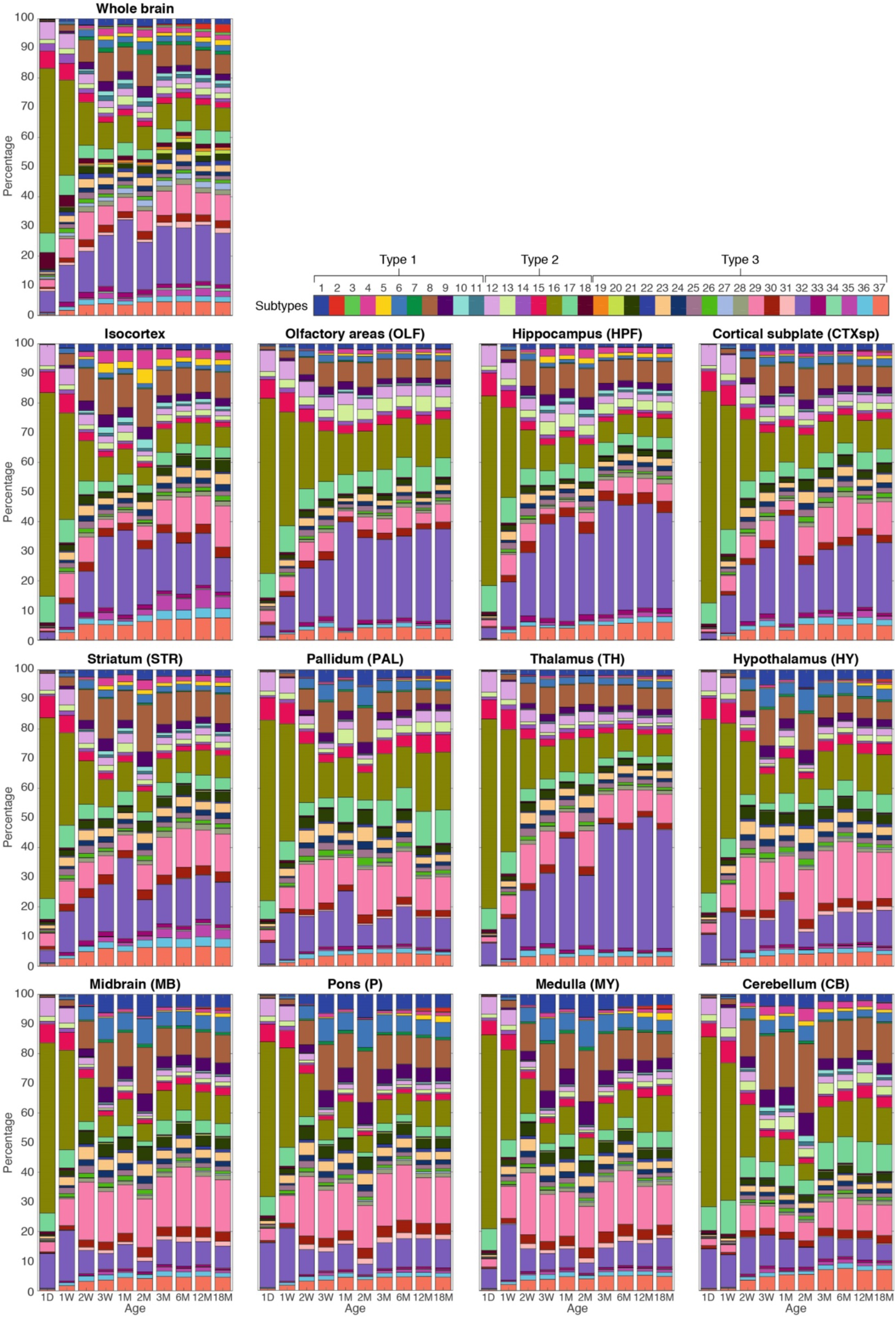
Relative abundance of synapse subtypes in whole brain and brain regions. Stacked bar plots of the percentage of 37 synapse subtypes in the whole brain and 12 main regions across the lifespan. Key: color codes for subtypes.

**Fig. S10:**
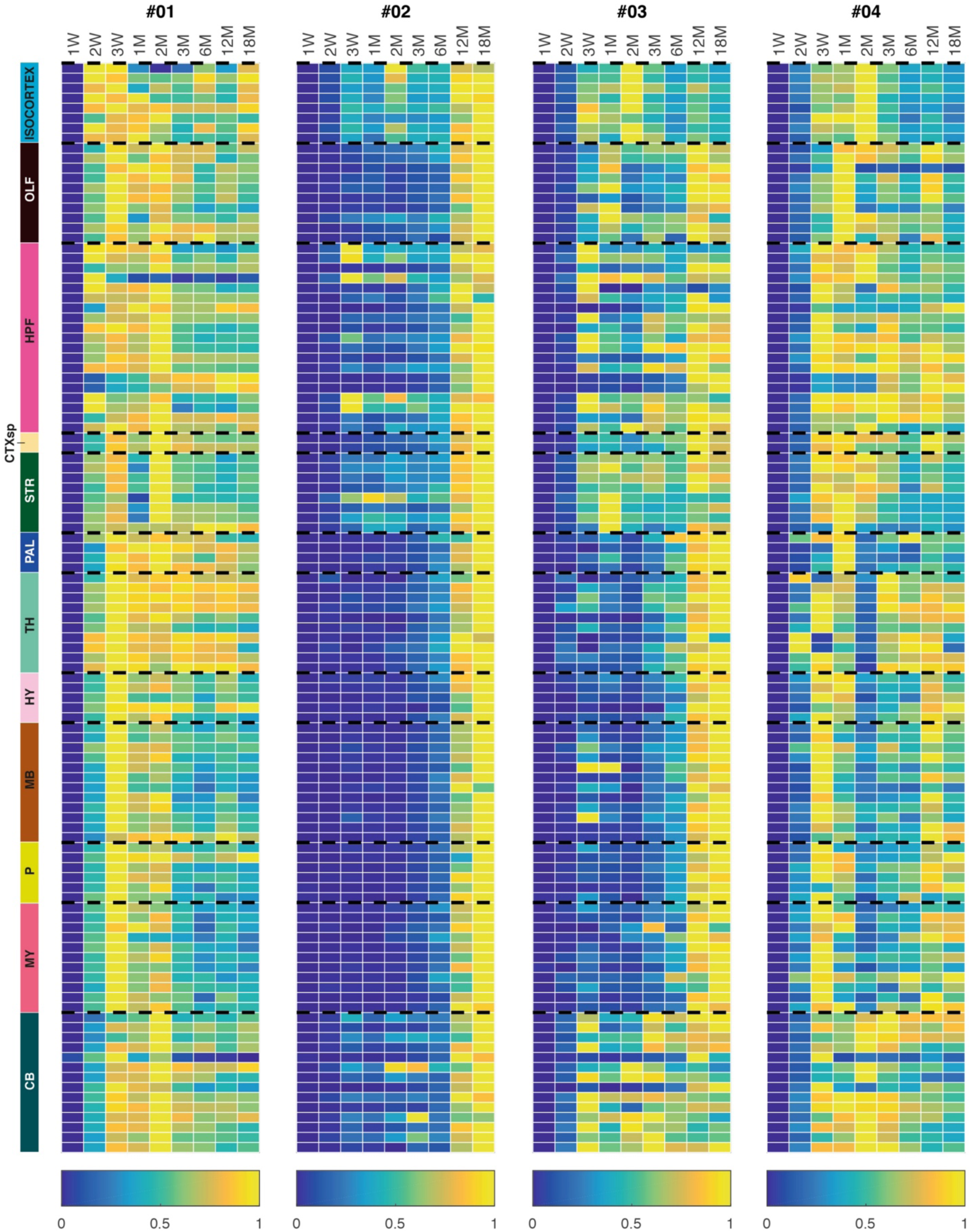
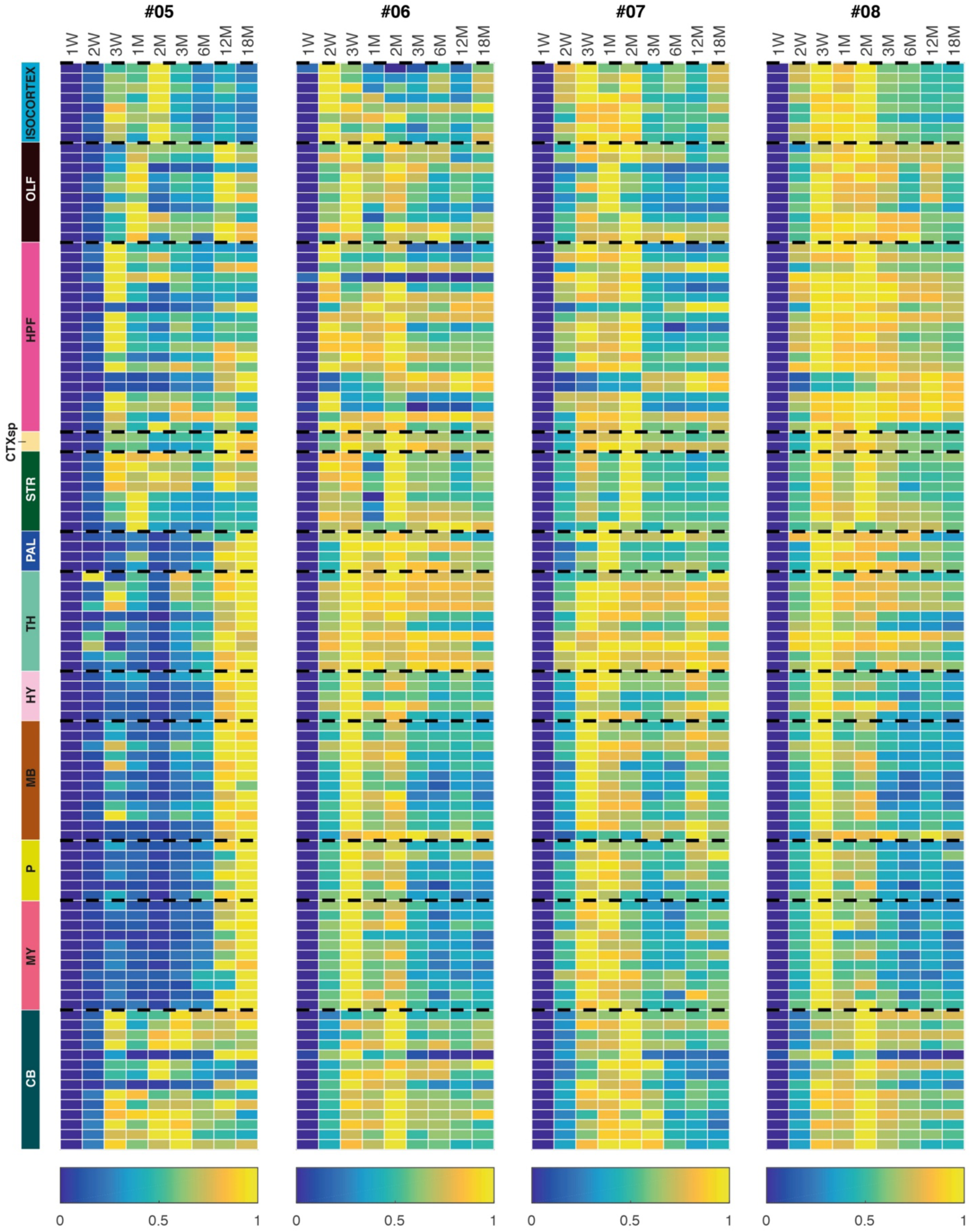
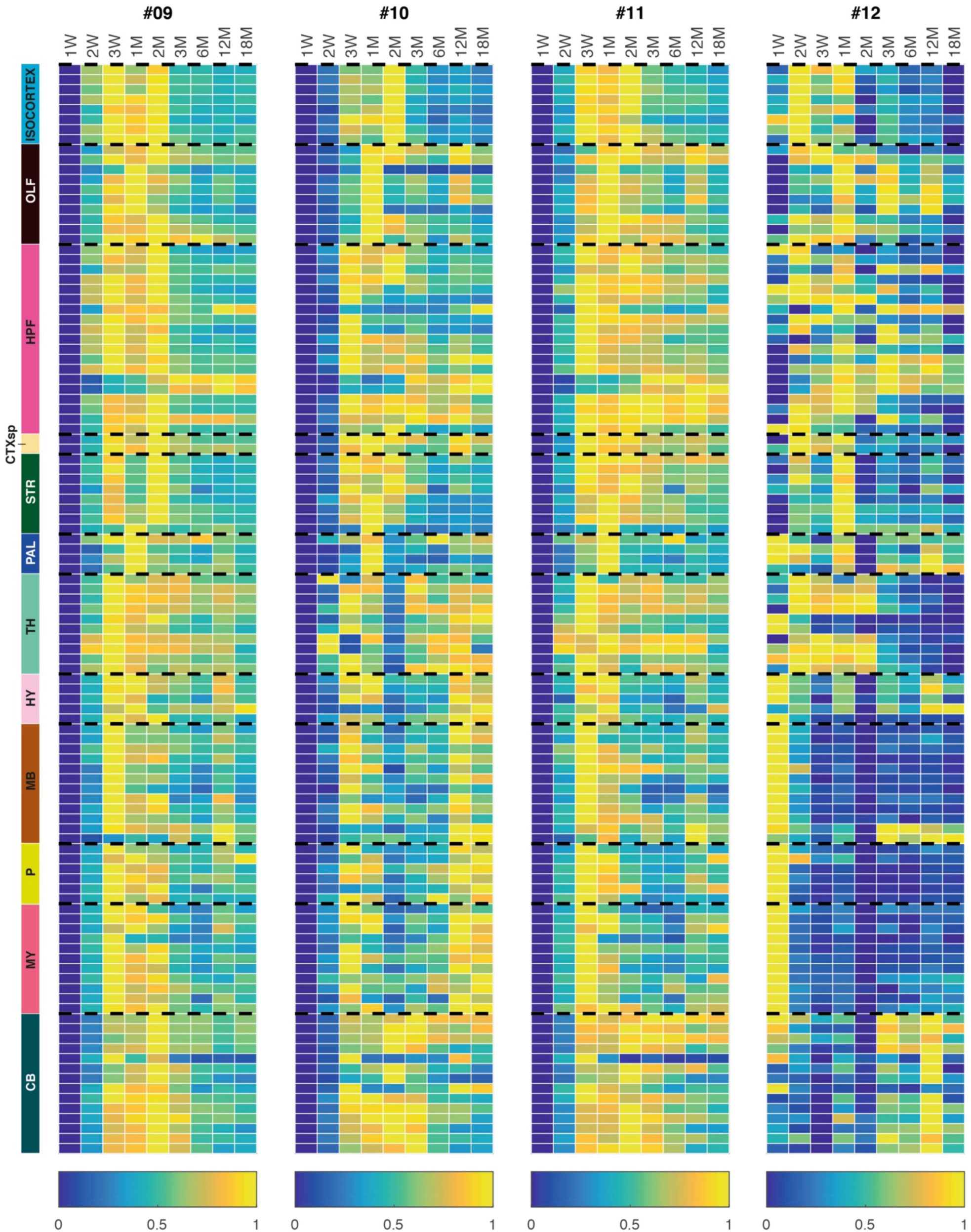
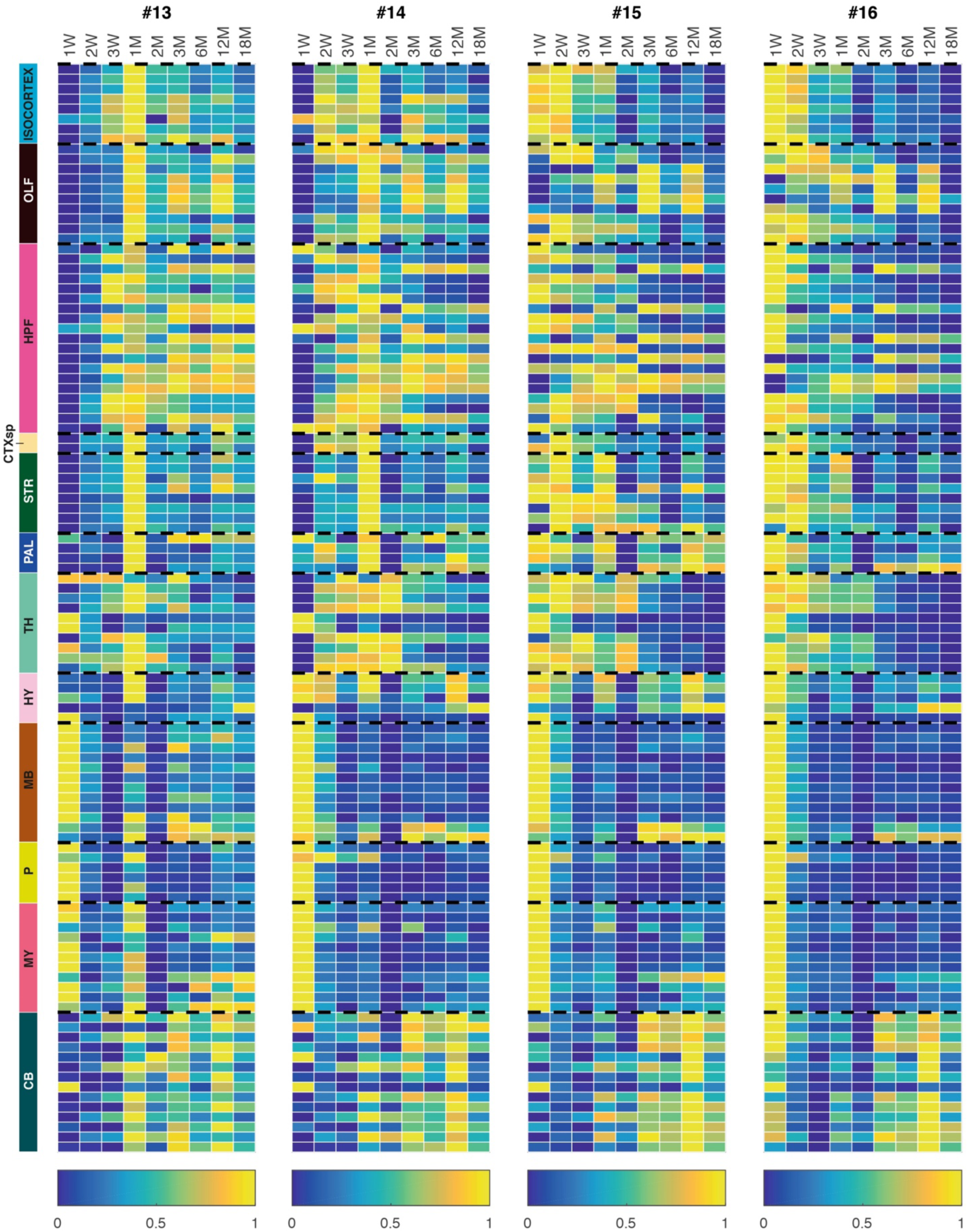
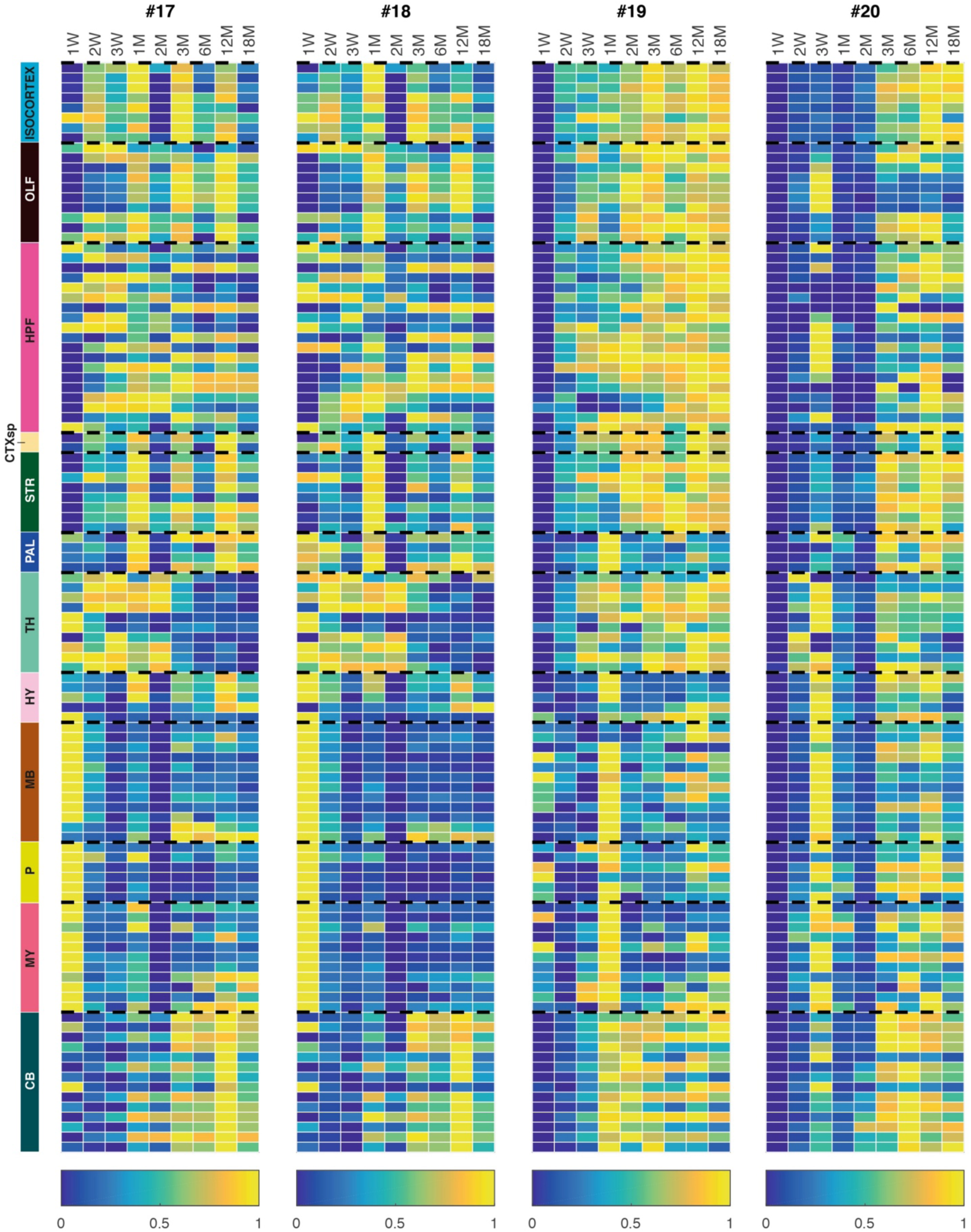
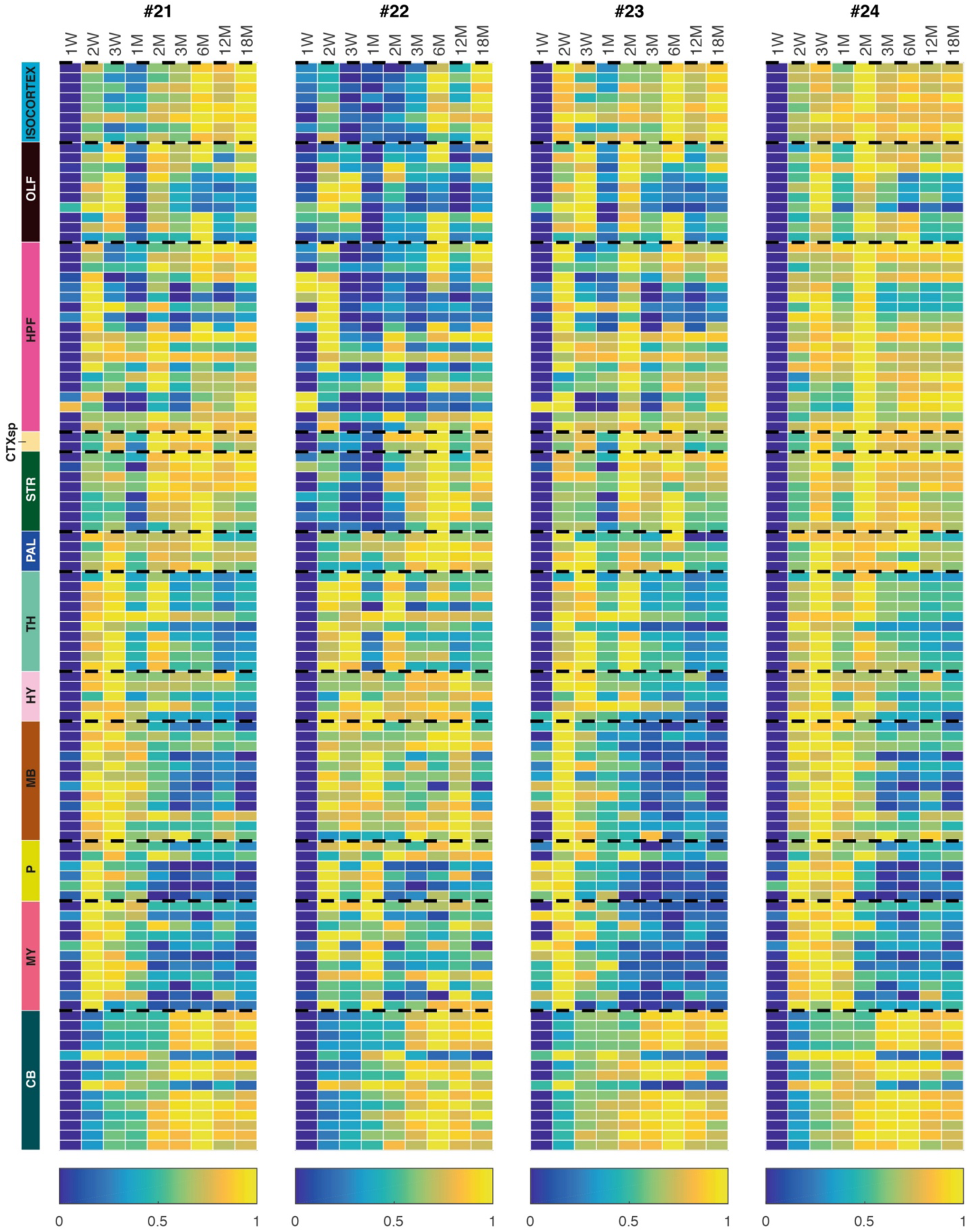
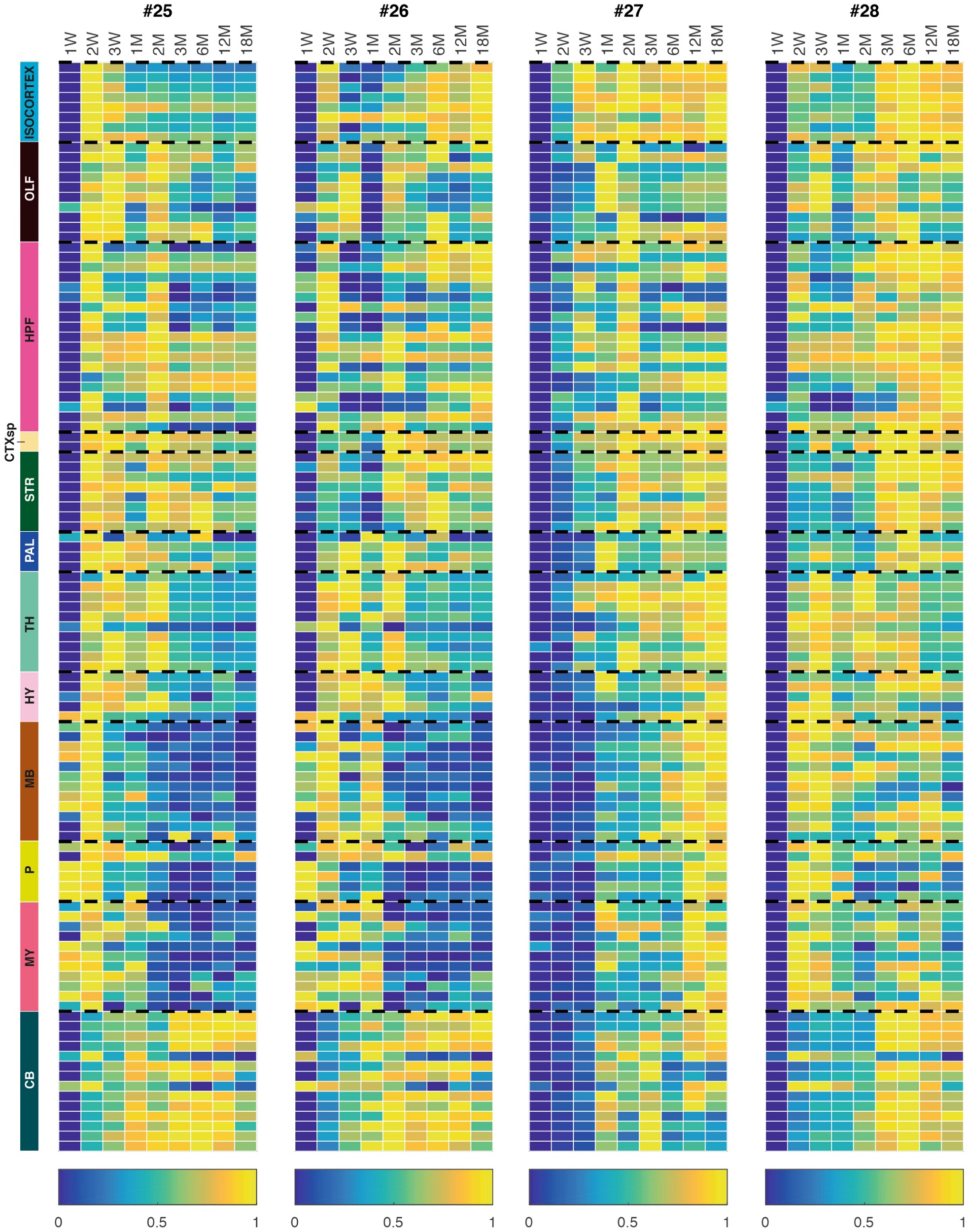
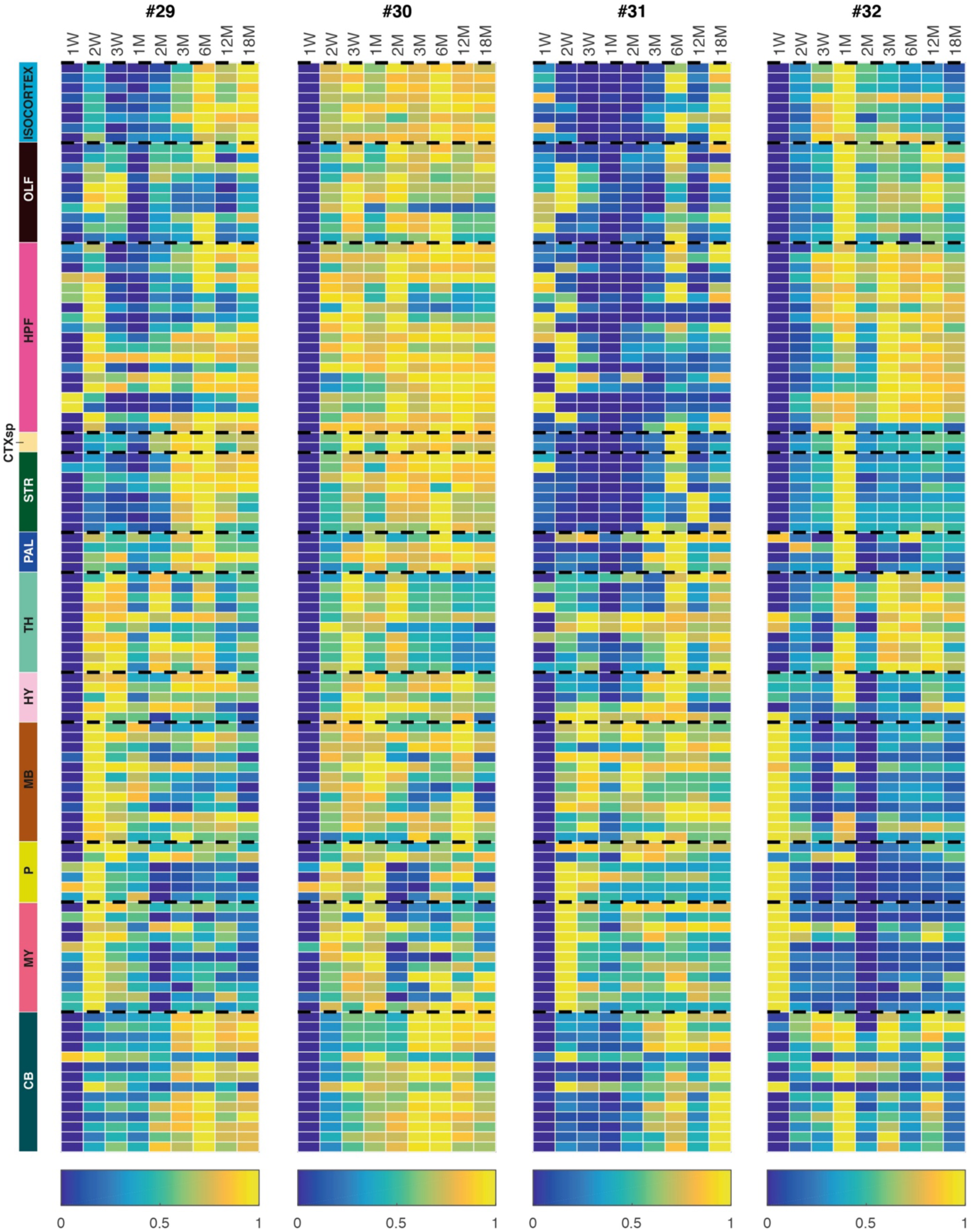
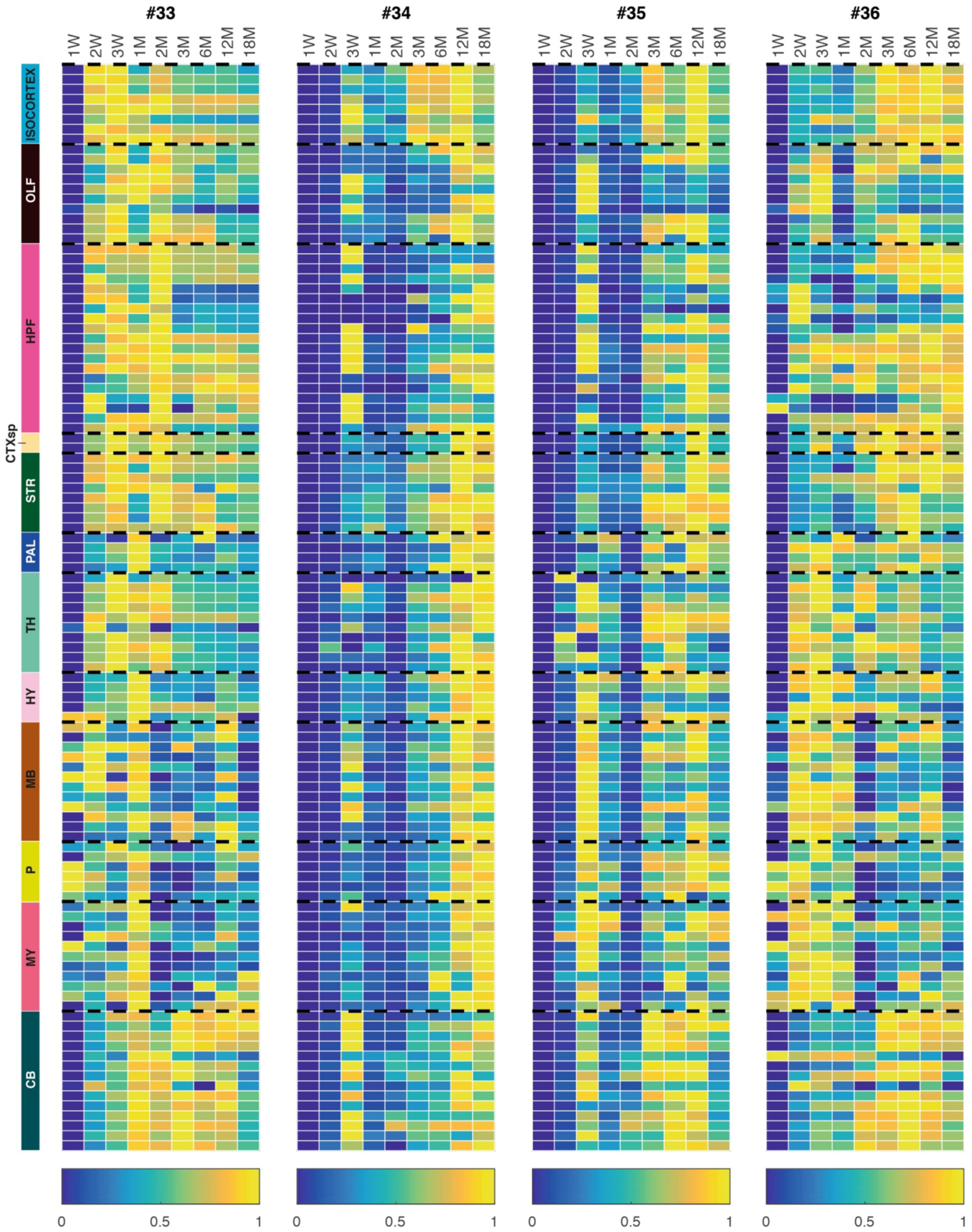
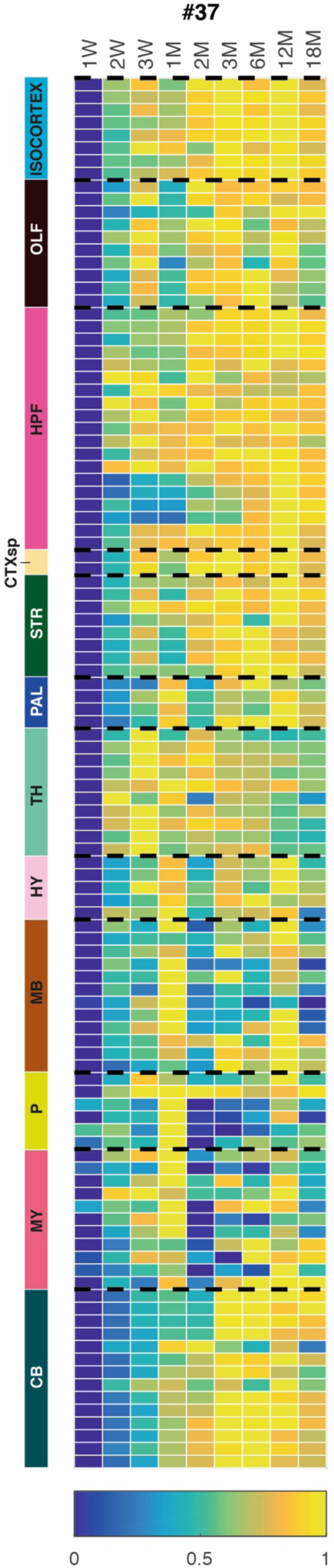
Heatmaps of lifespan trajectories of 37 synapse subtypes. Lifespan trajectories of 37 synapse subtypes by (normalized) density in each of 109 subregions (rows, see Table S1). Density in each subregion was normalized (0–1) to its maximal and minimal densities across the lifespan (rows).

**Fig. S11:**
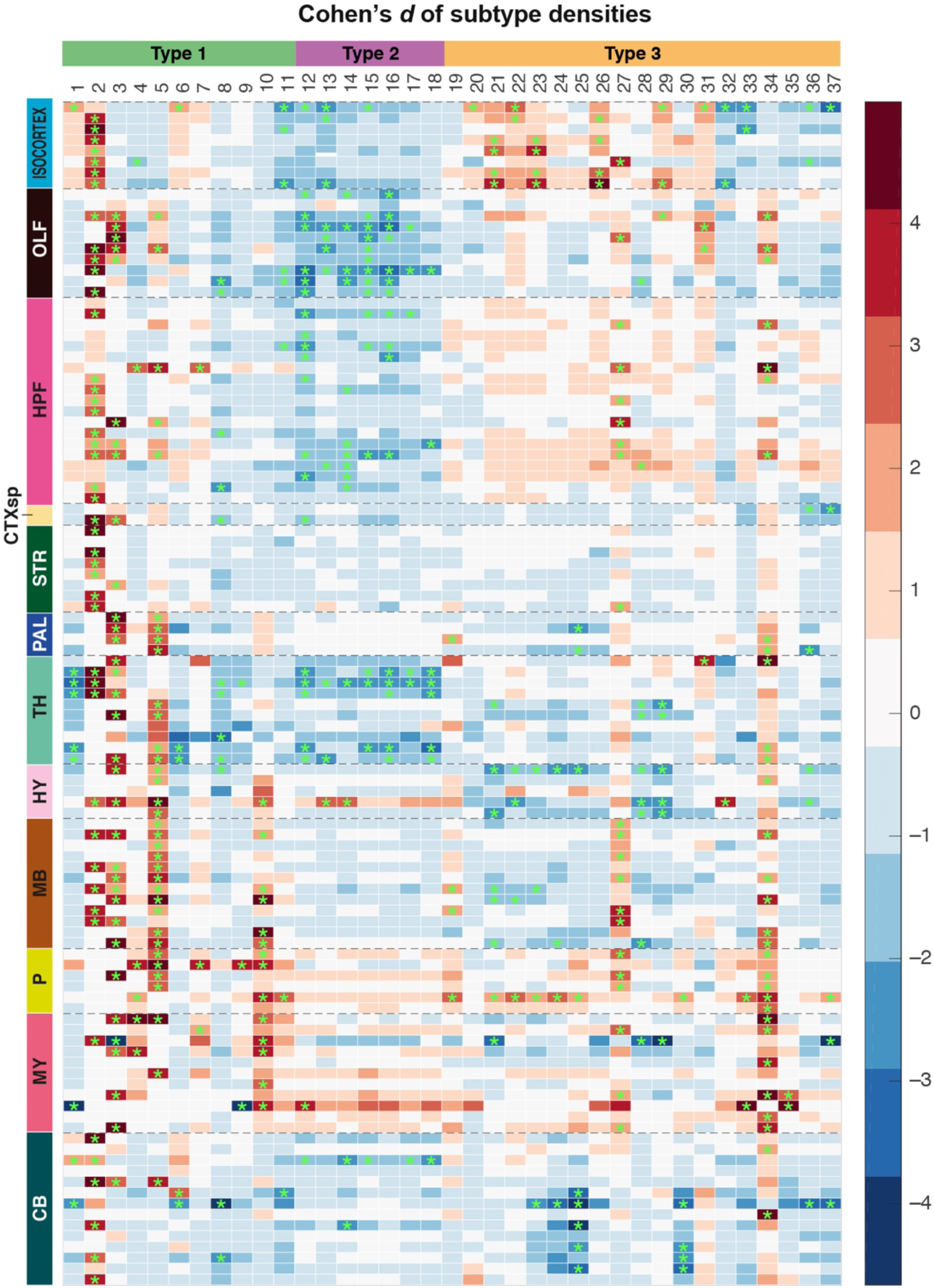
Changes in synapse subtypes in brain subregions between 3M and 18M. Heatmap showing the change (Cohen’s *d*, scale bar indicates increase in red and decrease in blue) in the density of 37 synapse subtypes in brain subregions between 3M and 18M. *P < 0.05 adjusted by the Benjamini-Hochberg multiple comparison correction after the Bayesian-based significance test method (*18*).

**Fig. S12:**
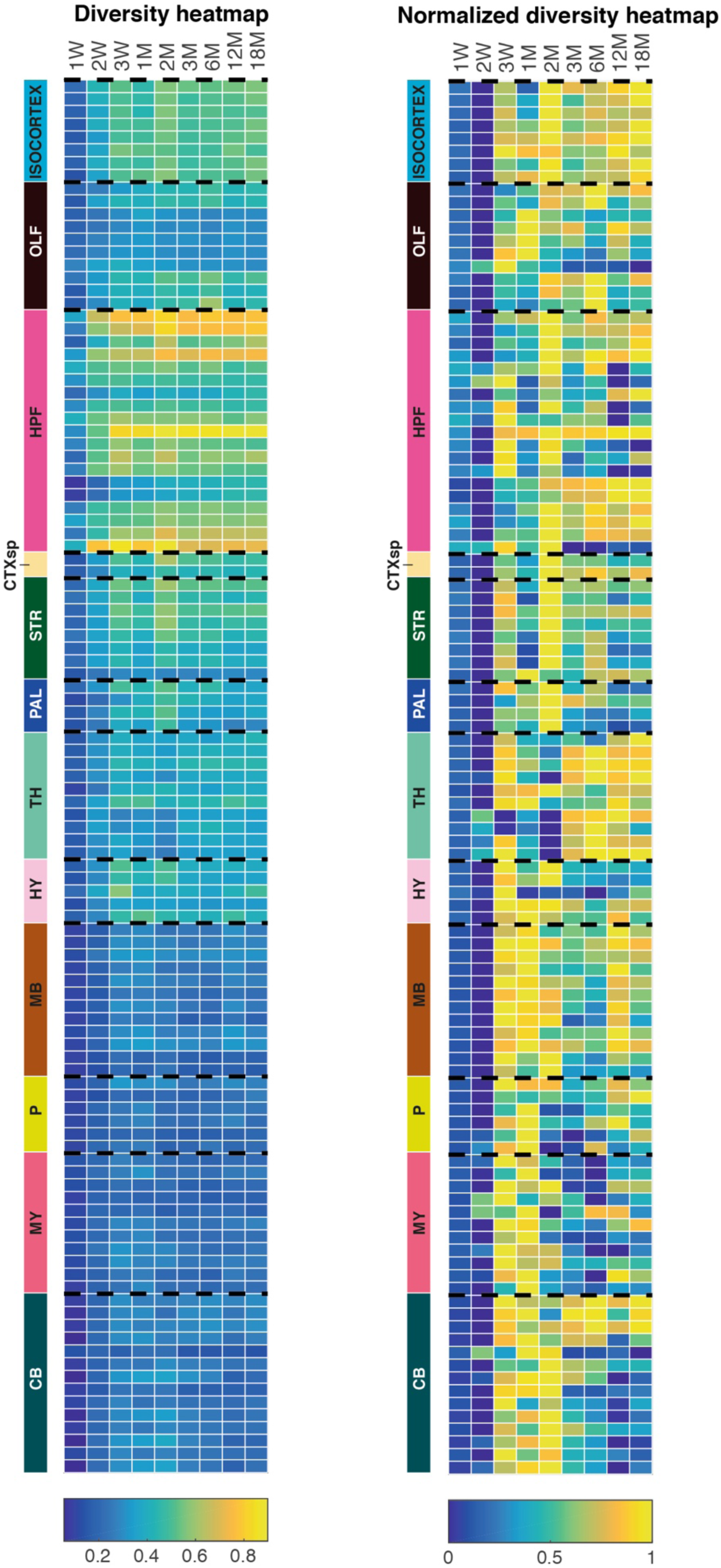
Lifespan trajectories of synapse diversity in brain subregions. (Left) Lifespan trajectories of raw (mean) values of synapse diversity in brain subregions (see Table S1 for subregion names). Scale, Shannon entropy (0 – 1). (Right) Lifespan trajectories of subregion-normalized values of synapse diversity in brain subregions (see Table S1 for subregion names). Scale, normalized Shannon entropy (0 – 1).

**Fig. S13:**
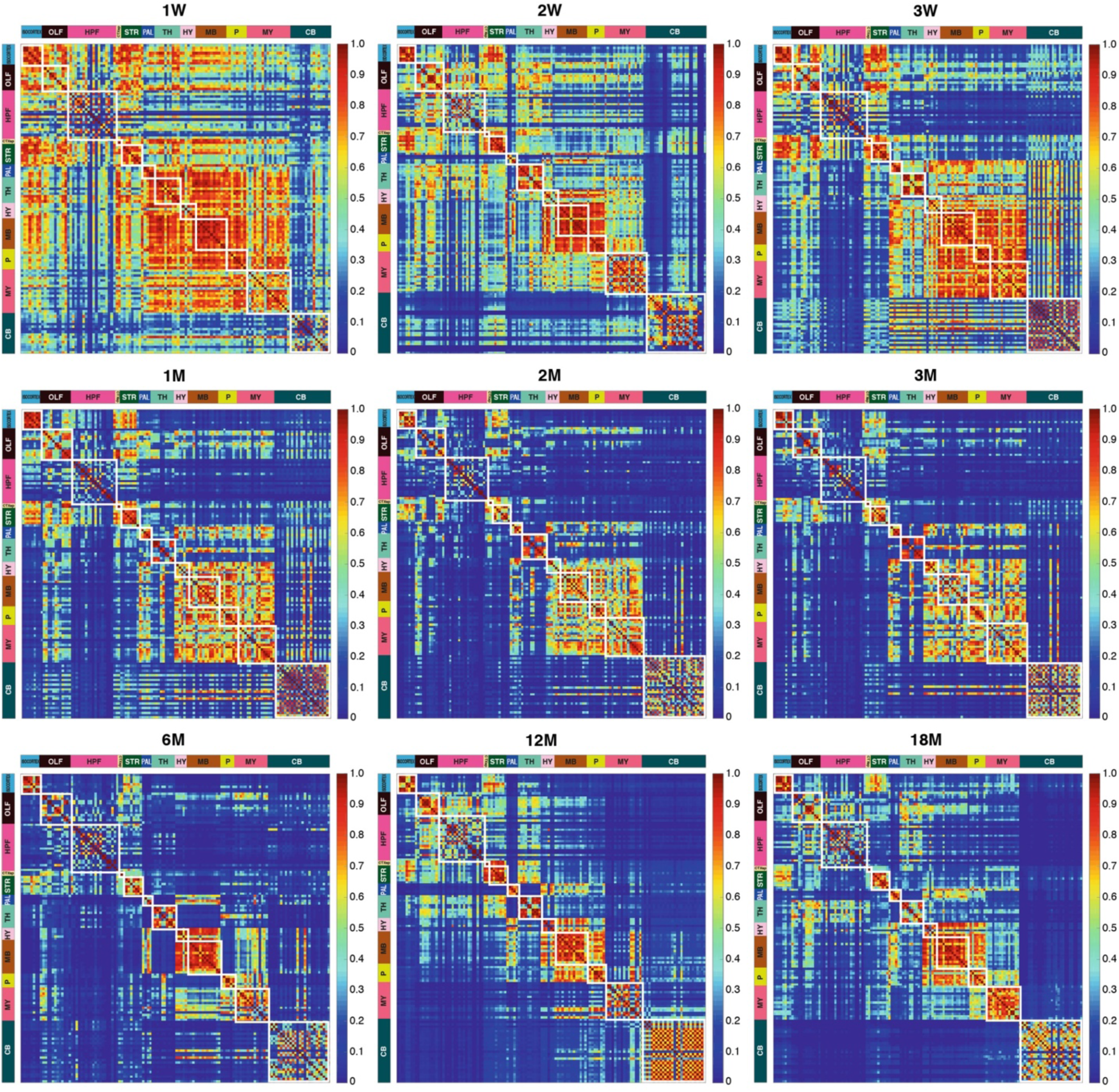
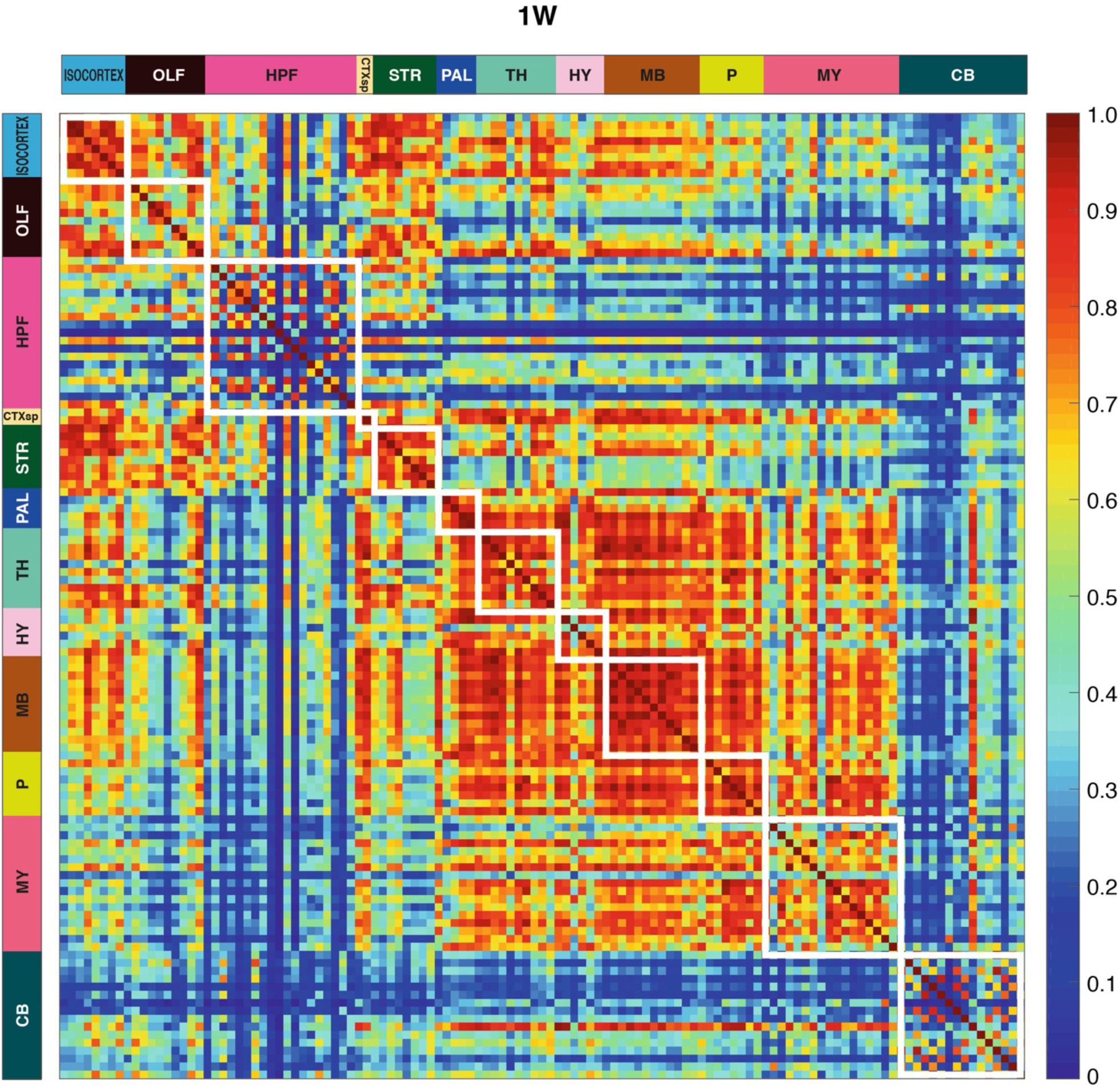
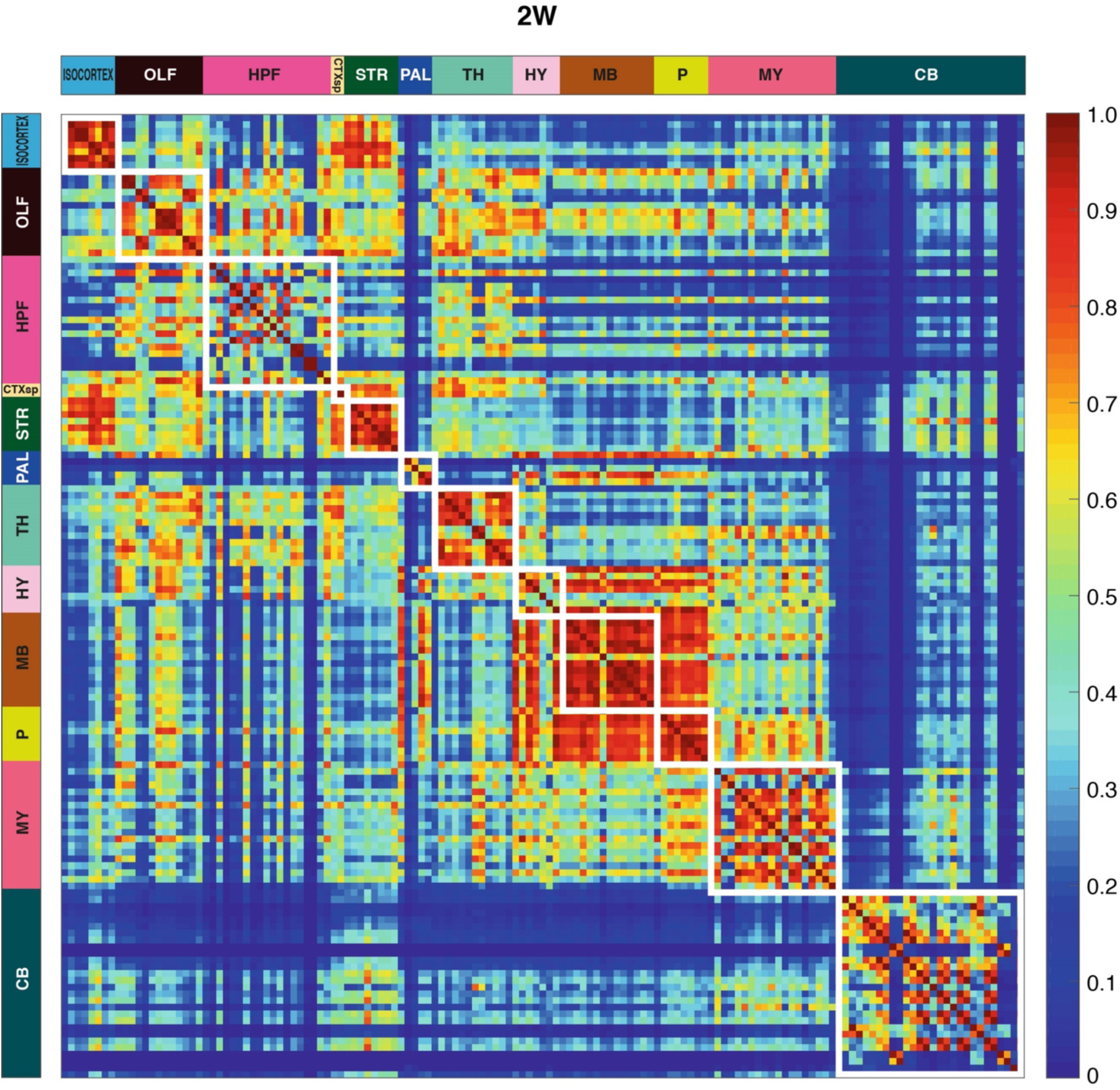
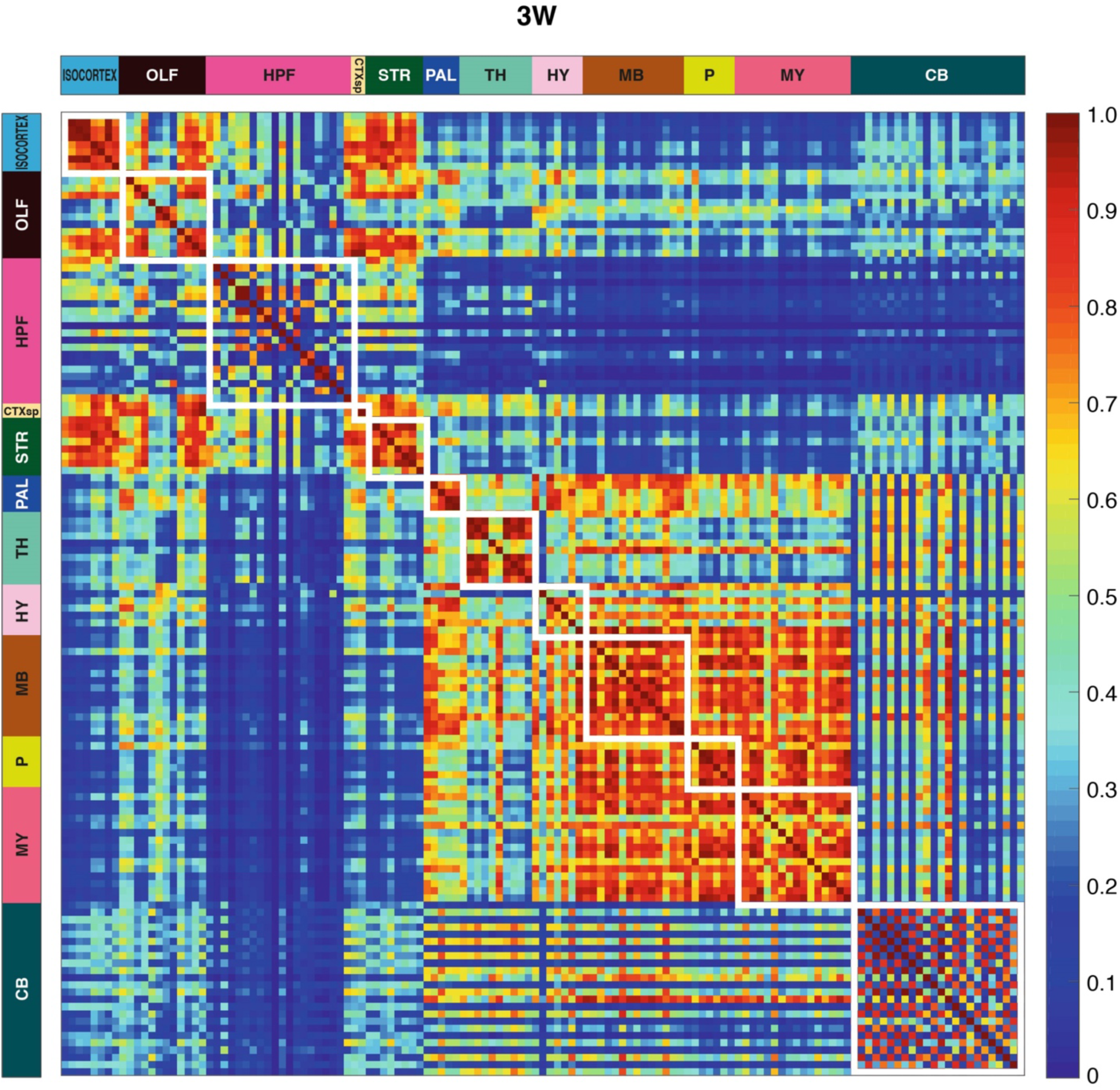
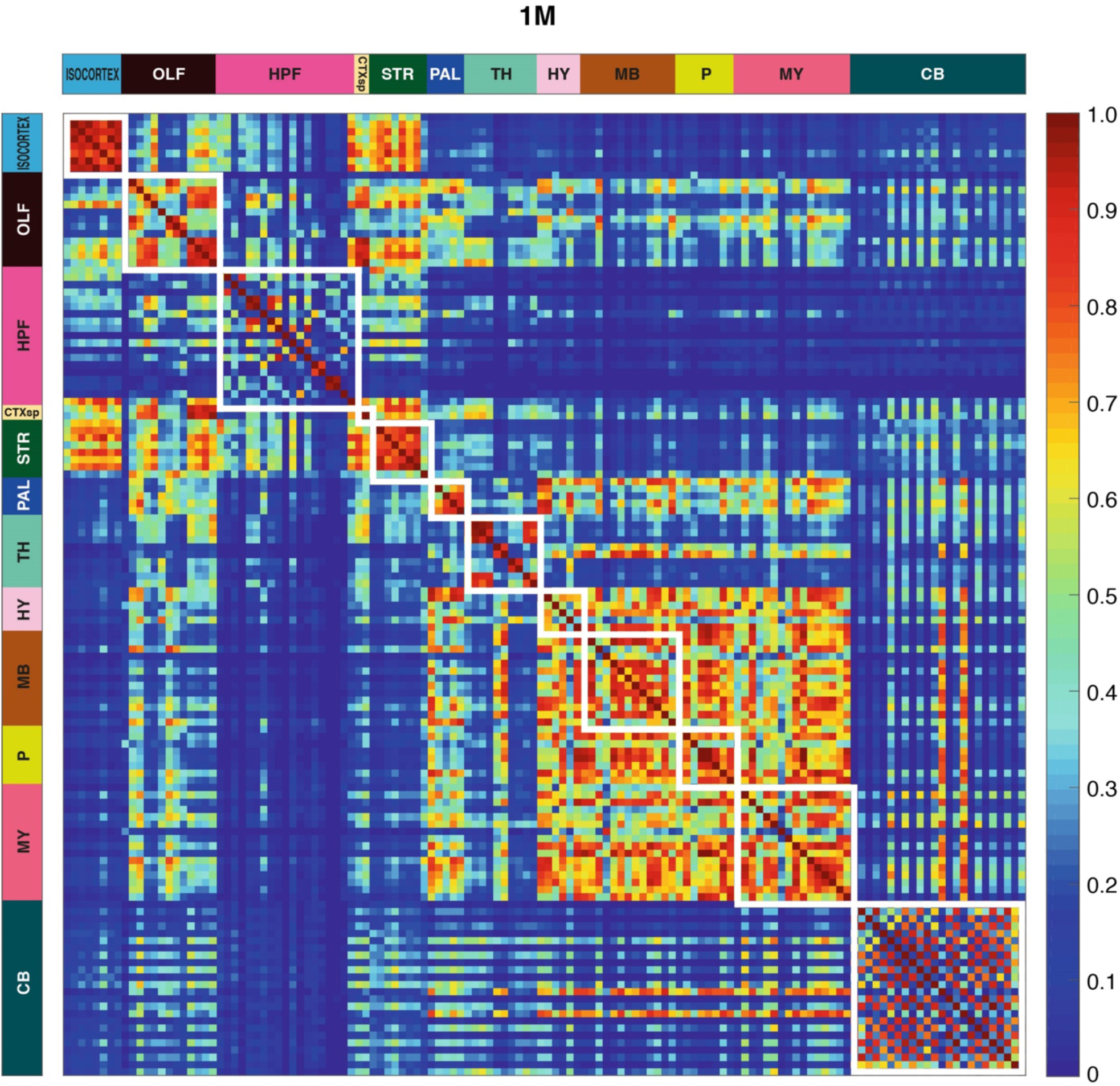
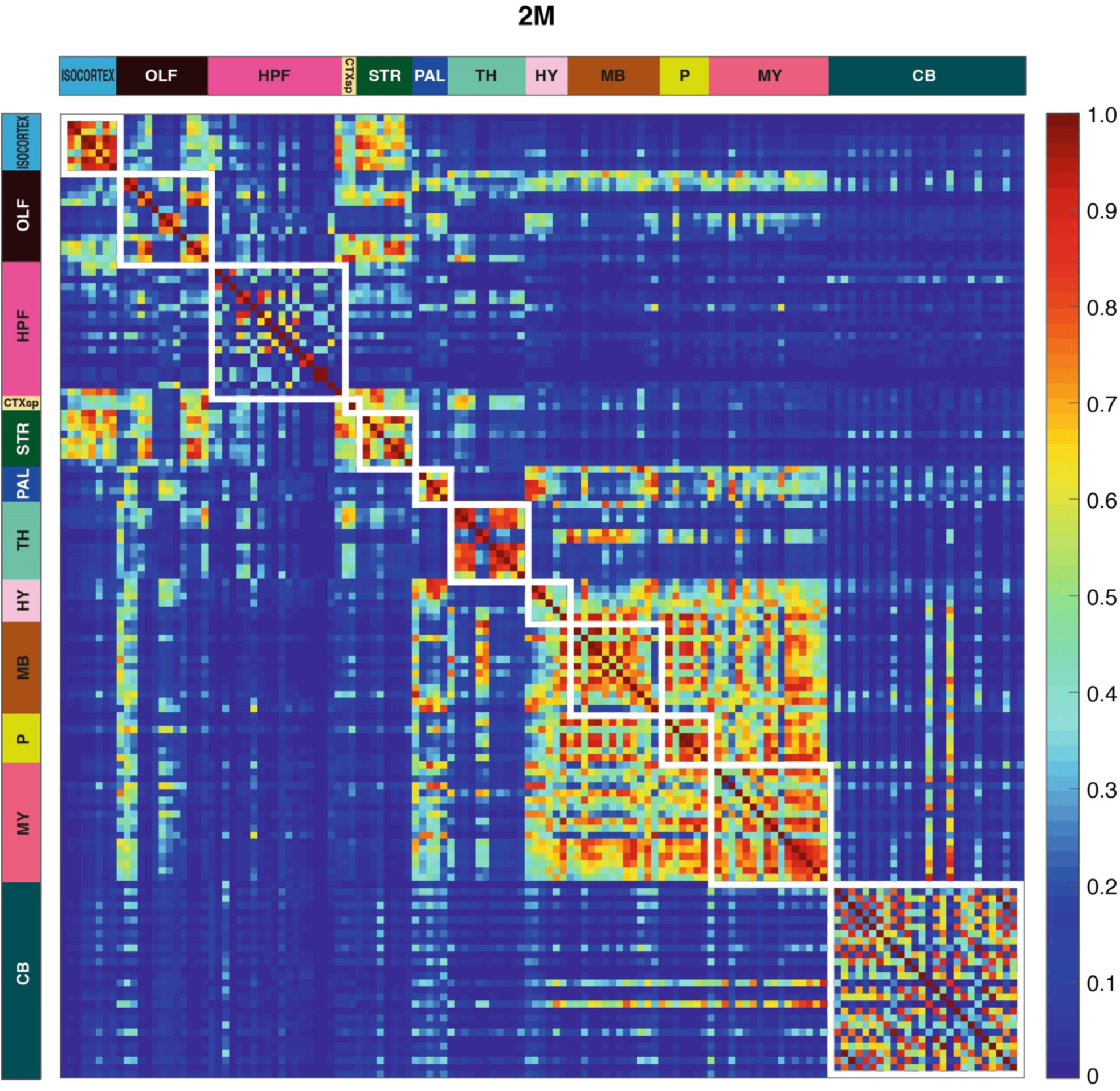
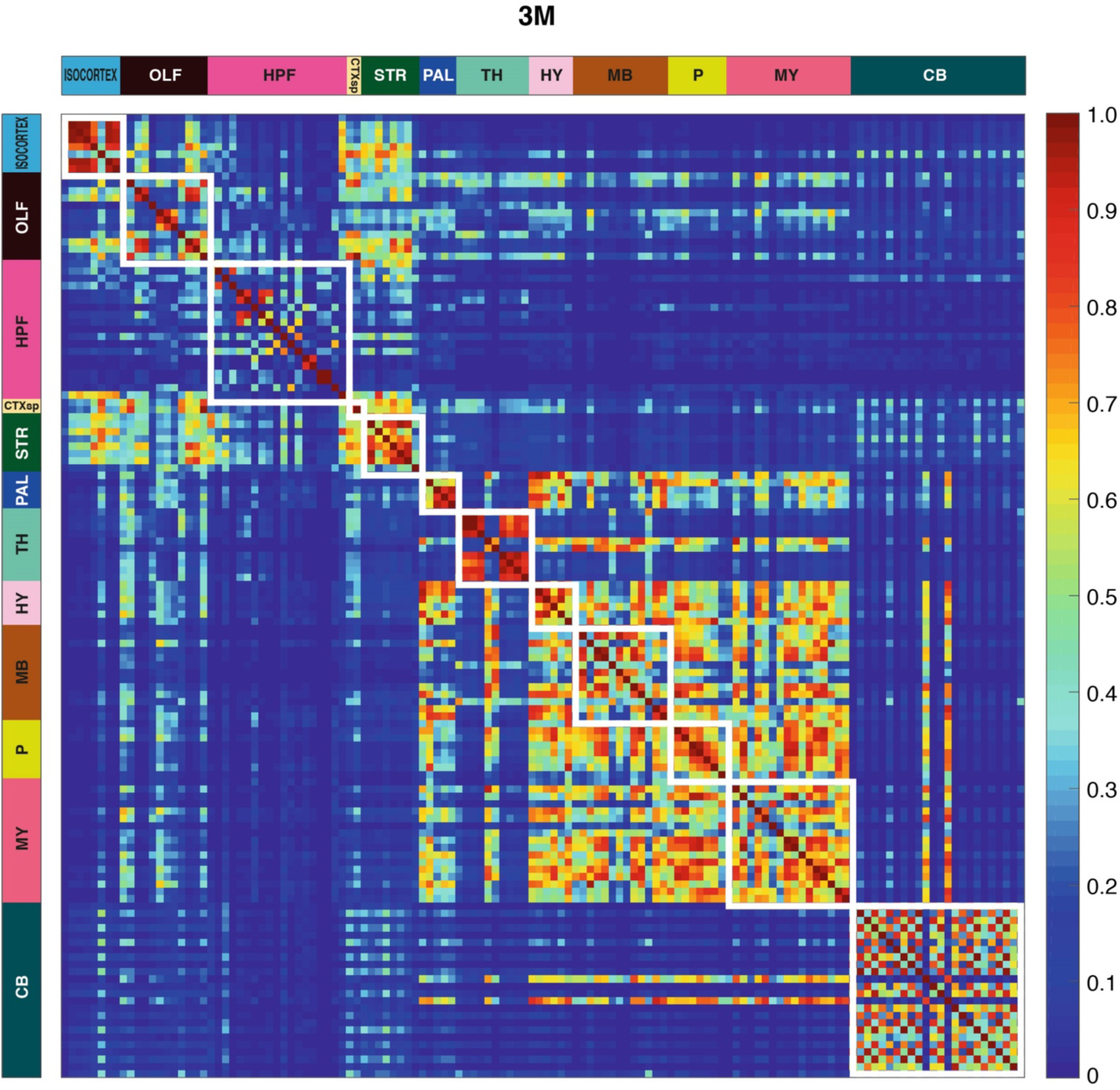
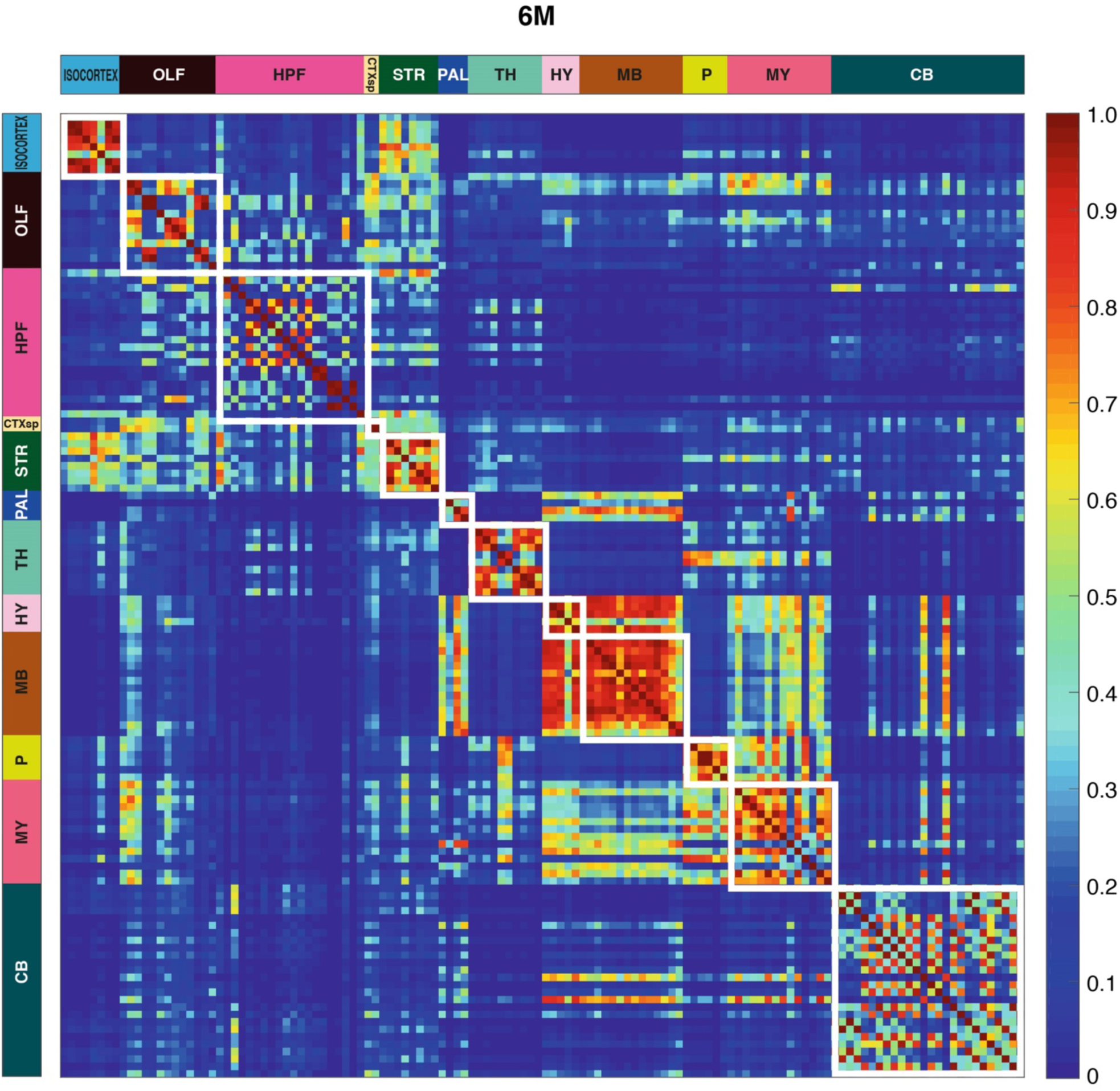
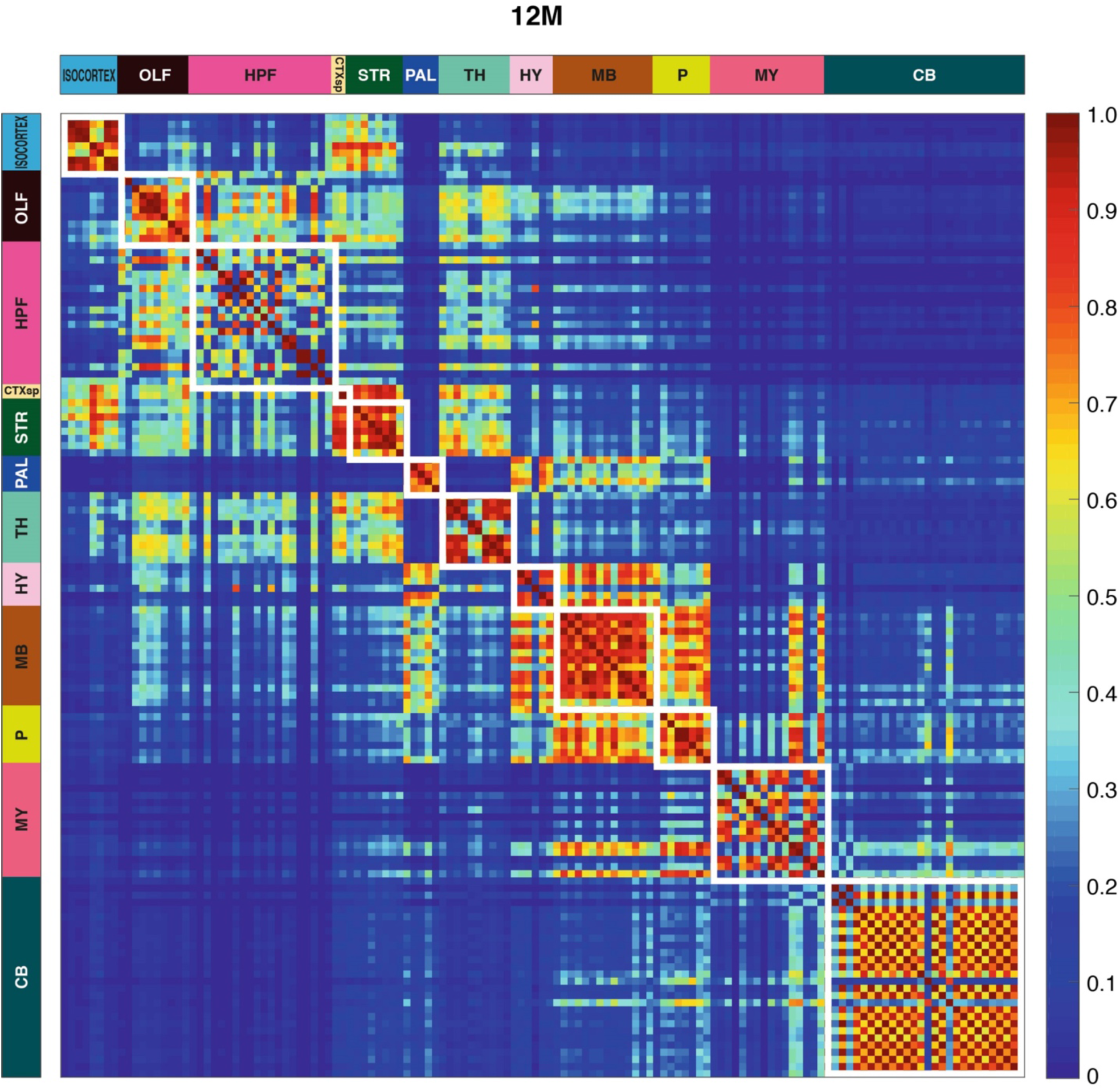
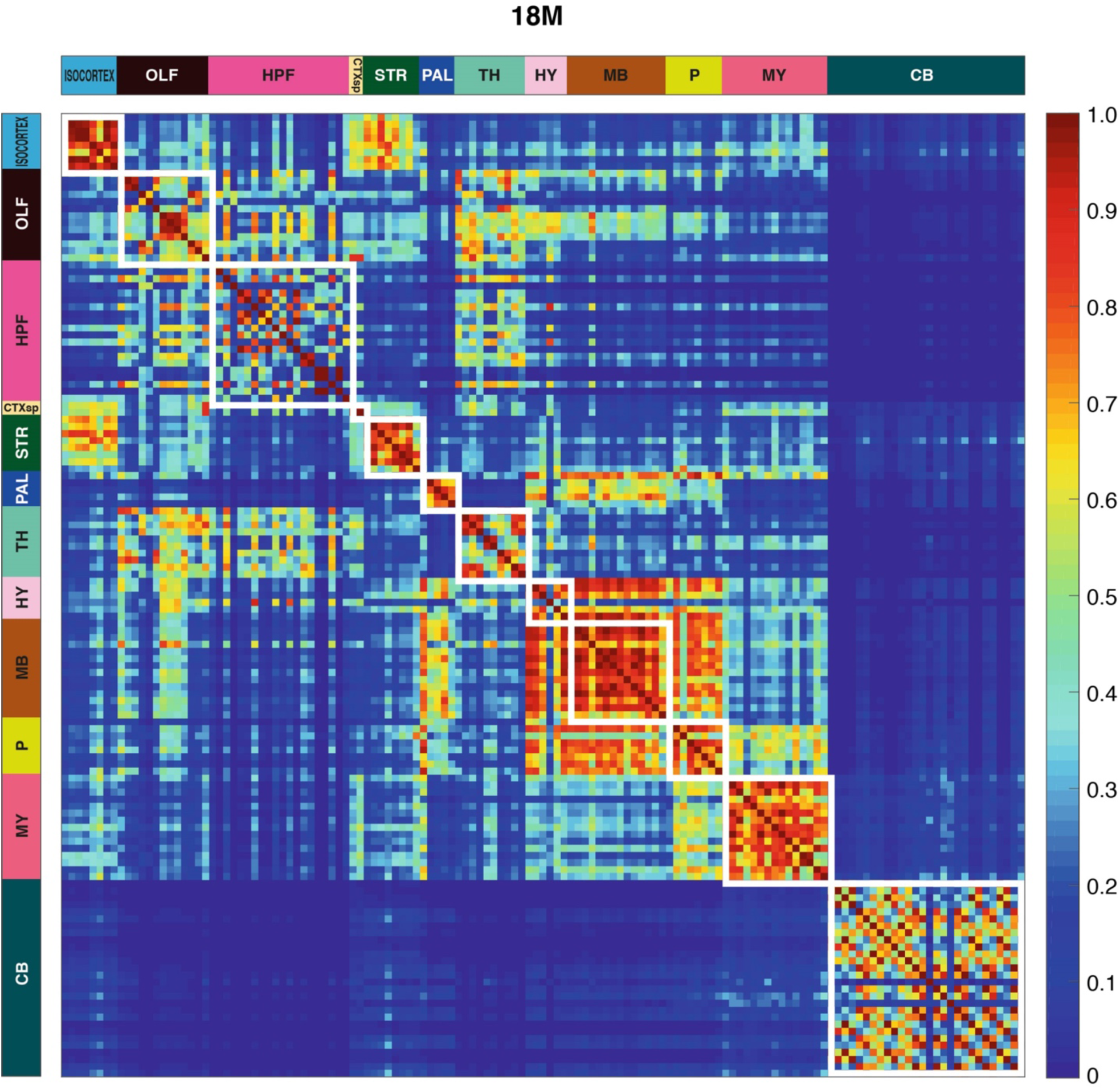
Brain subregion similarity matrices. Matrix of similarities between pairs of subregions (rows and columns) from 1W until 18M (collected and single enlarged matrices shown). White boxes indicate the subregions that belong to the same main brain region (see color code on the left and top). Color scale bar indicates the similarity level ranging from 0 (blue) to 1 (red). The similarity is calculated based on the differences in synaptome parameters between two subregions, details of which can be found in the previous work (15) (see Table S1 for subregion names). The first figure shows the 9 similarity matrices together, then an enlargement of each is shown.

**Fig. S14:**
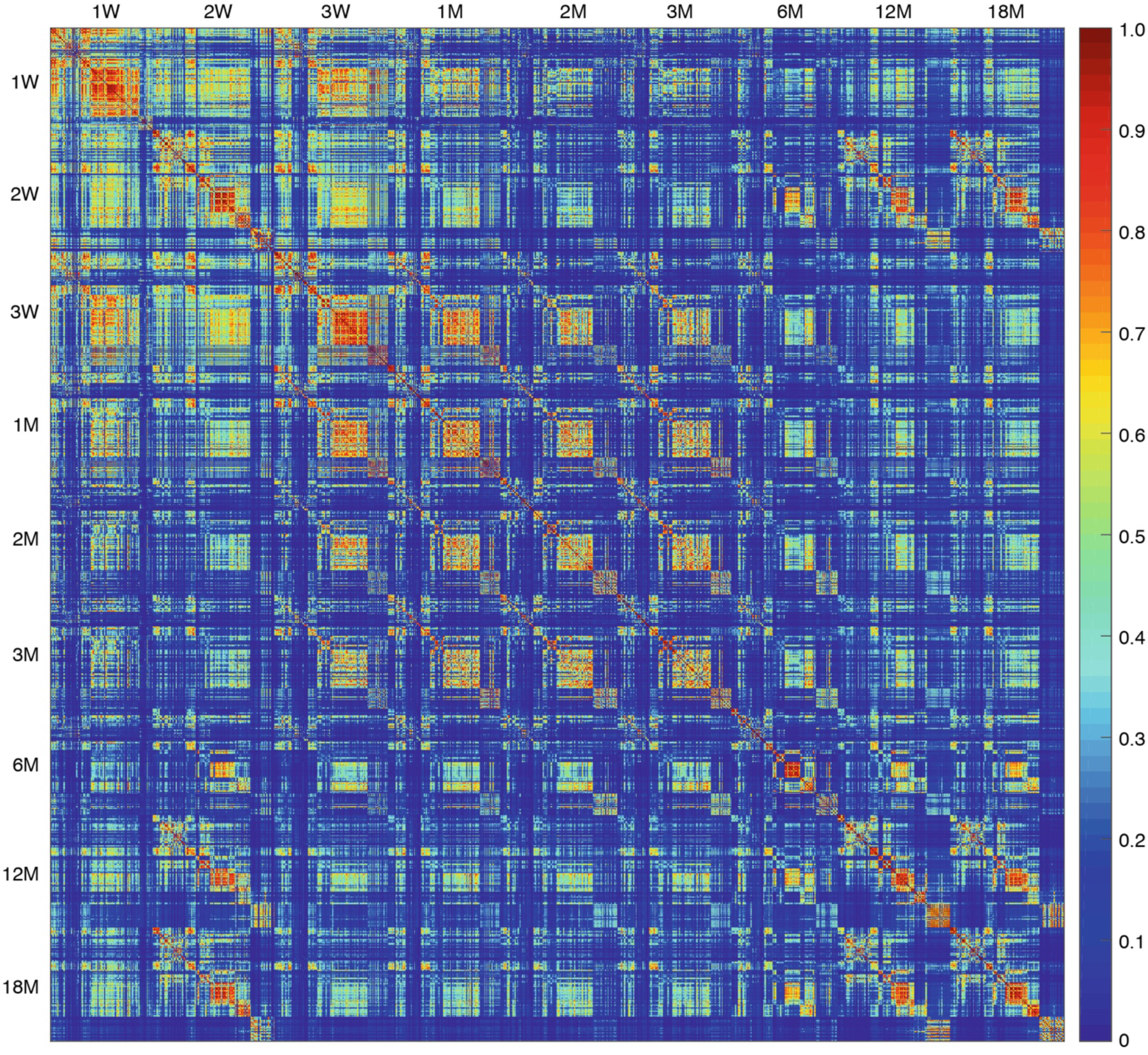
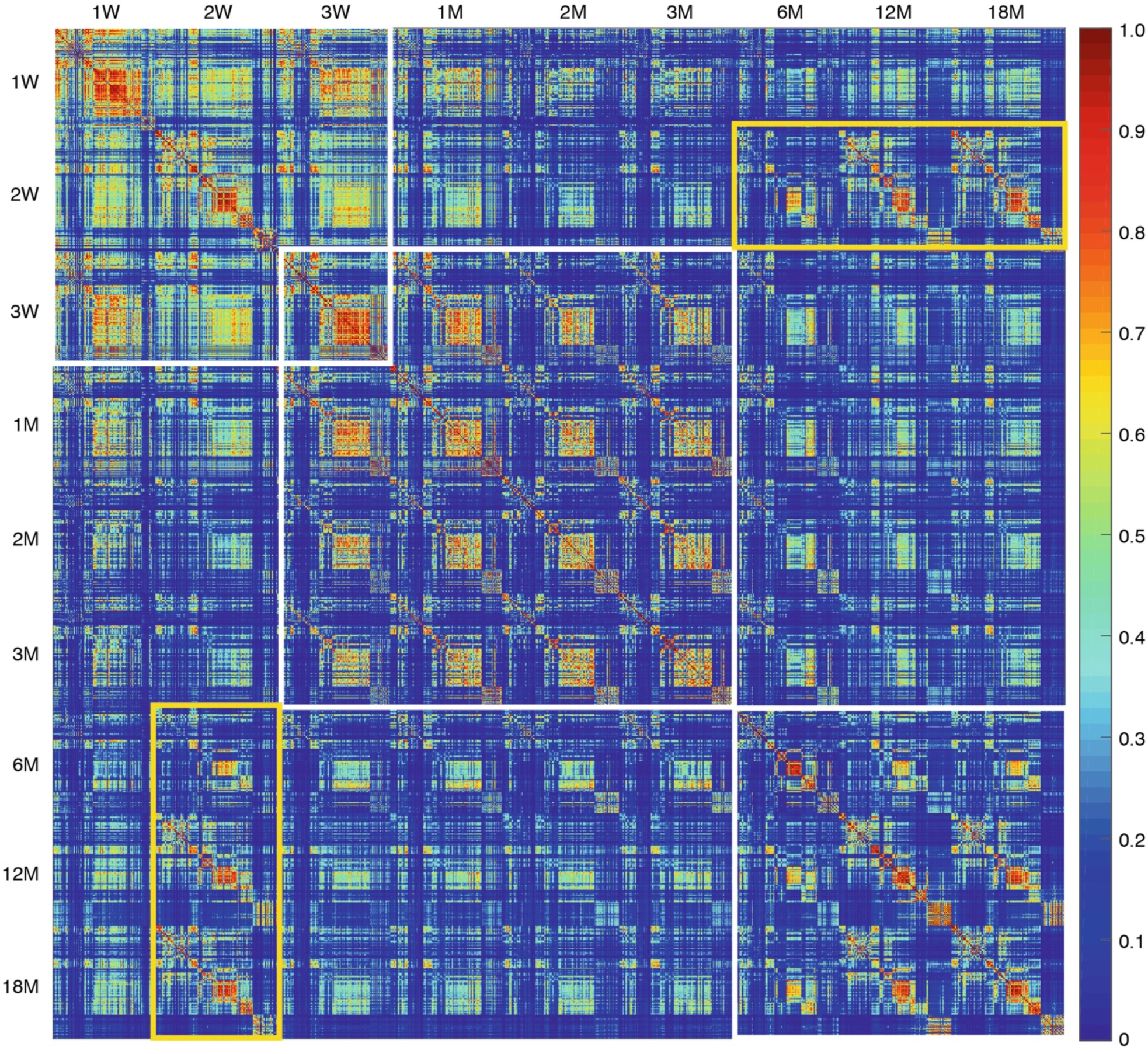
Hypersimilarity matrix comparing all brain subregions across all ages. Enlargement of hypersimilarity matrix in Fig. 3D without superimposed boxes (top) and with boxes (bottom); white boxes indicate the main three clusters, which correspond to LSA-I, -II and -III. Yellow box shows increased similarity of the old brain with young brain.

**Fig. S15:**
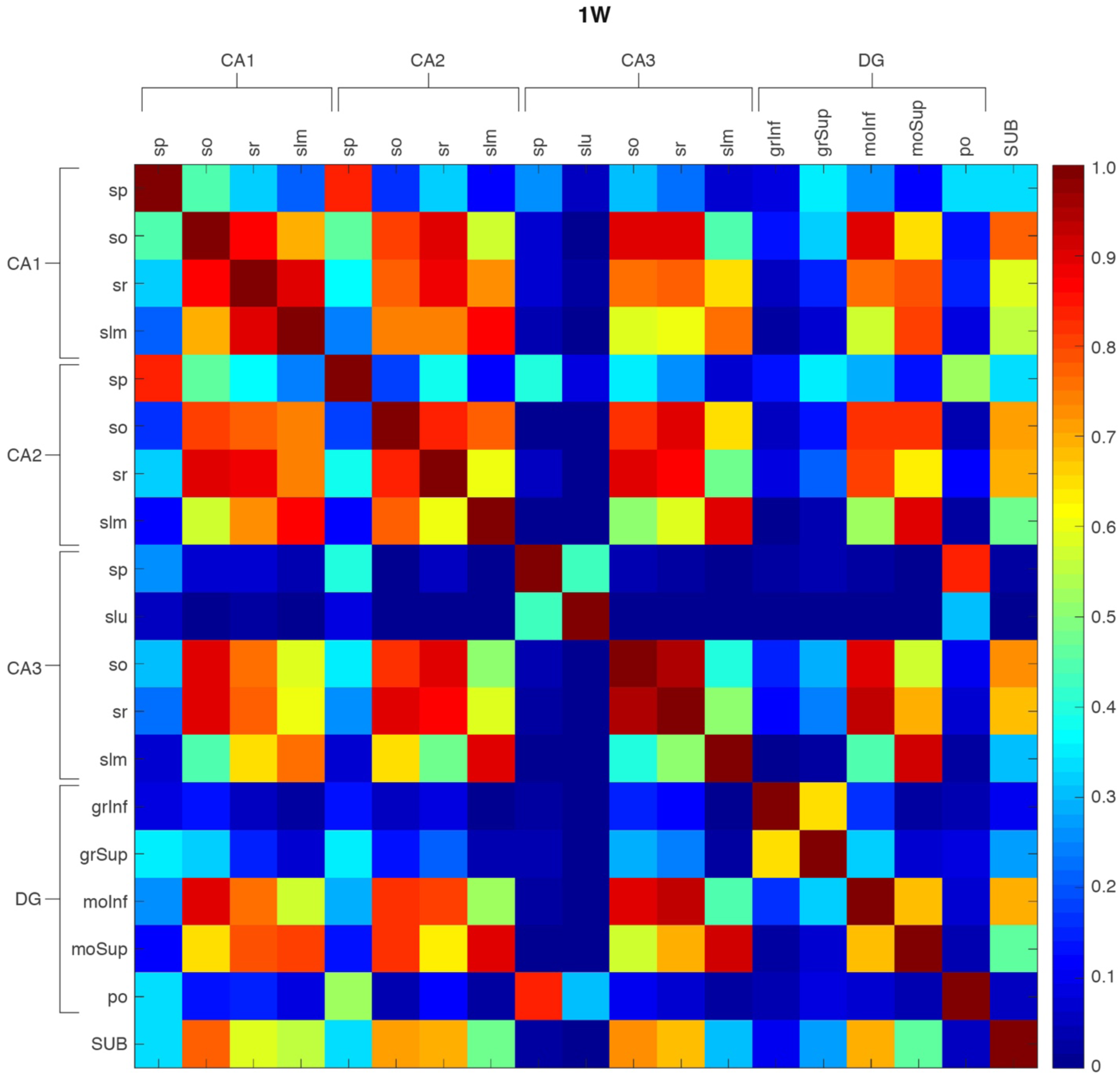
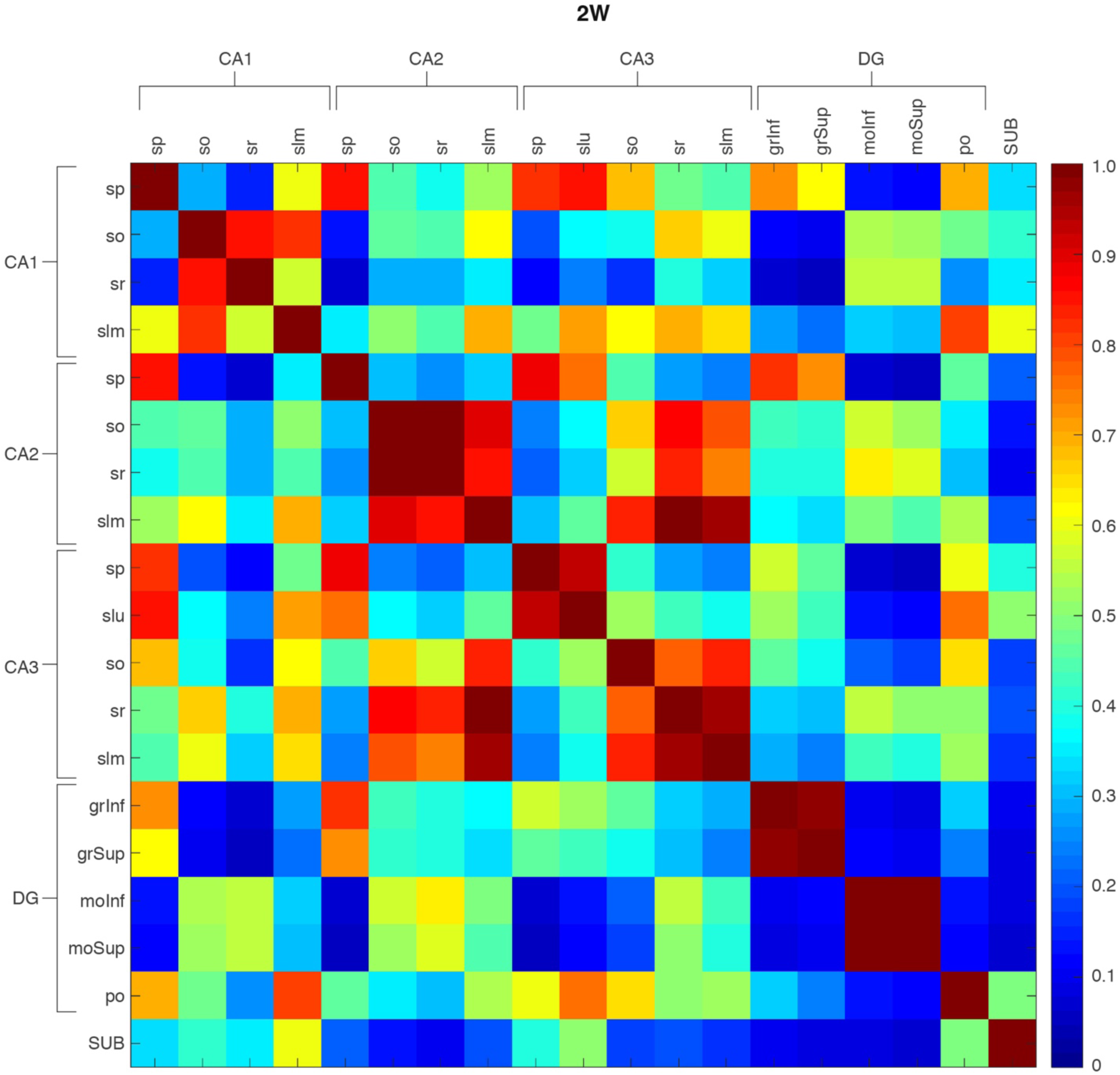
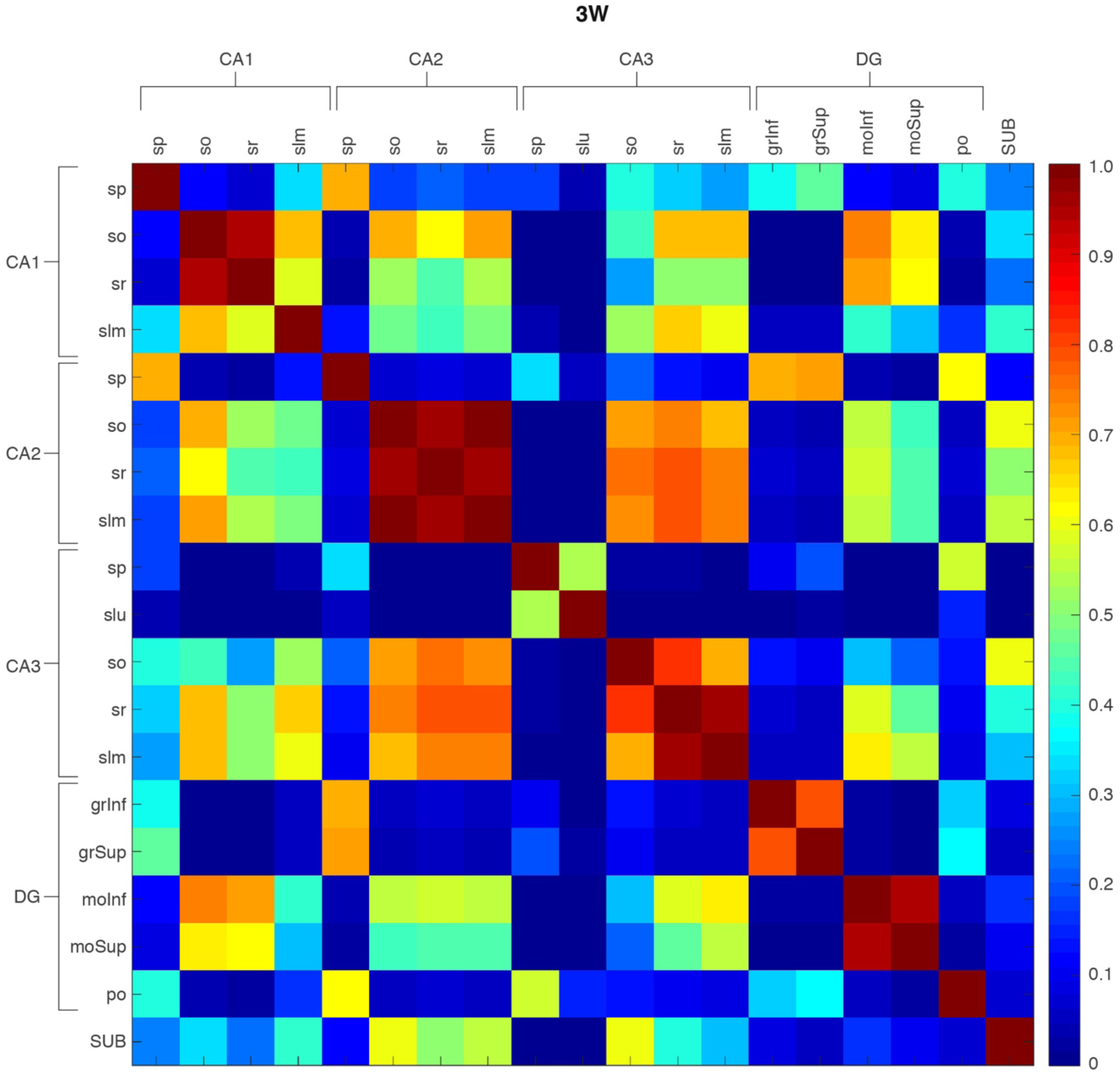
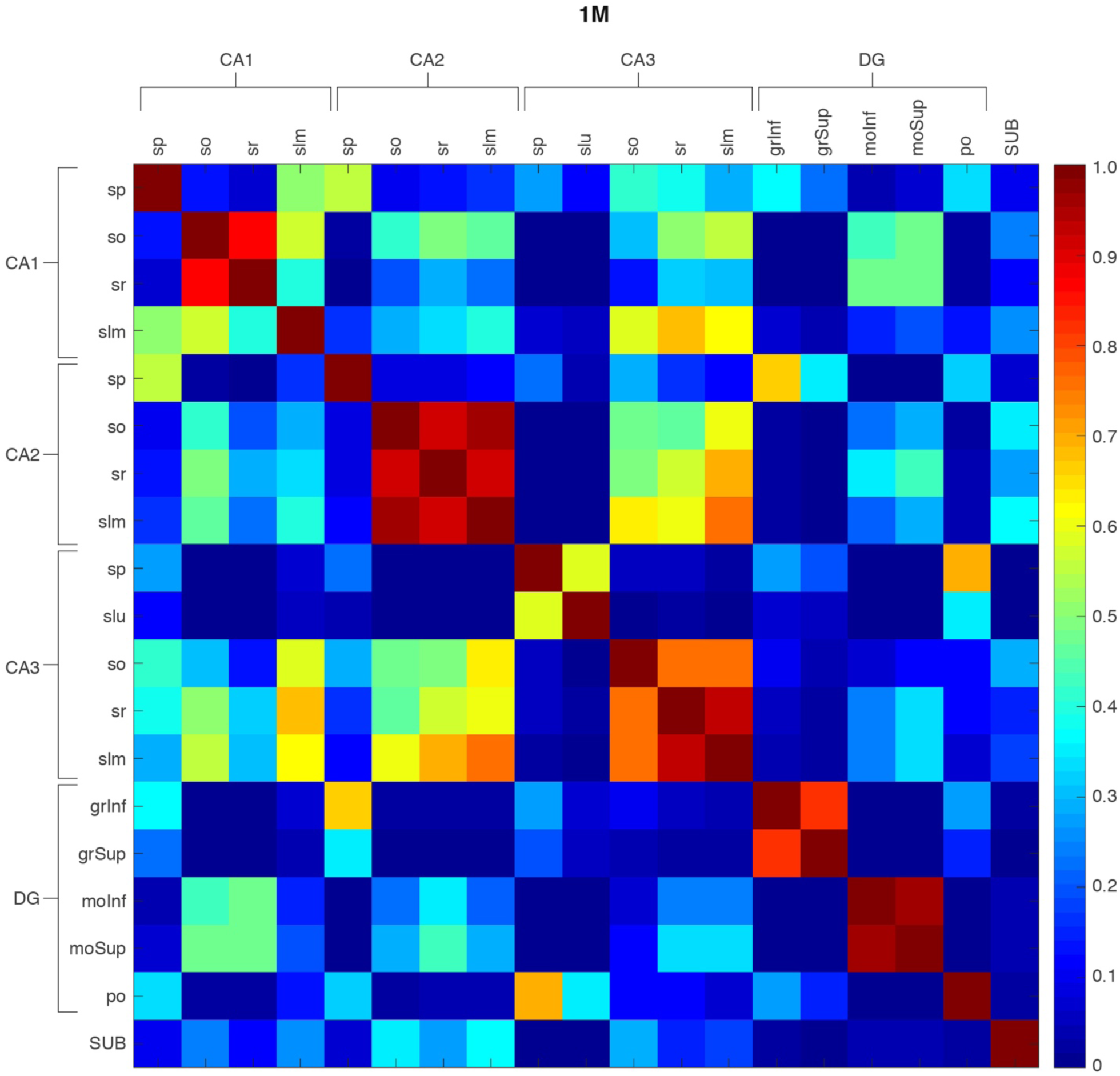
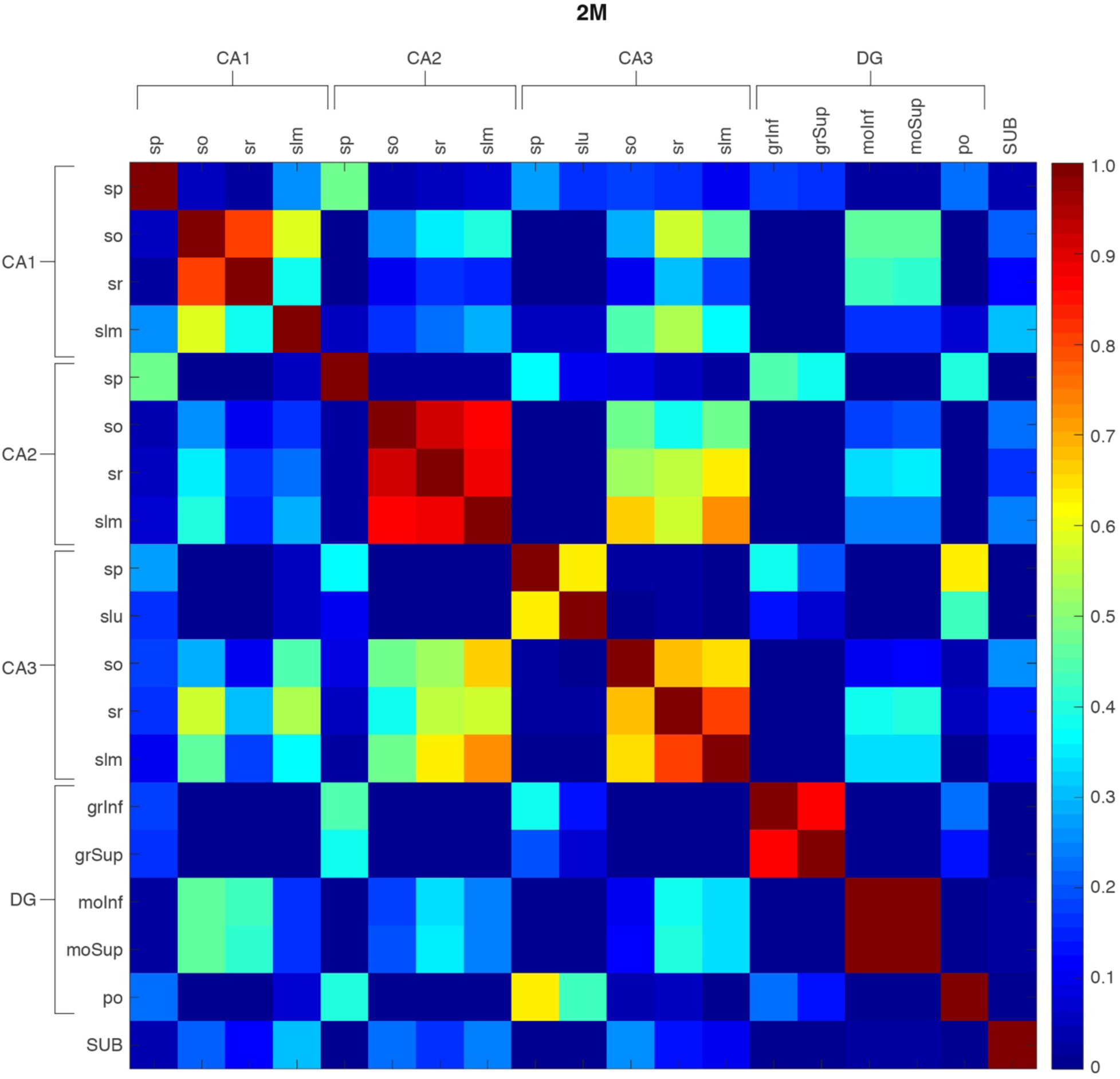
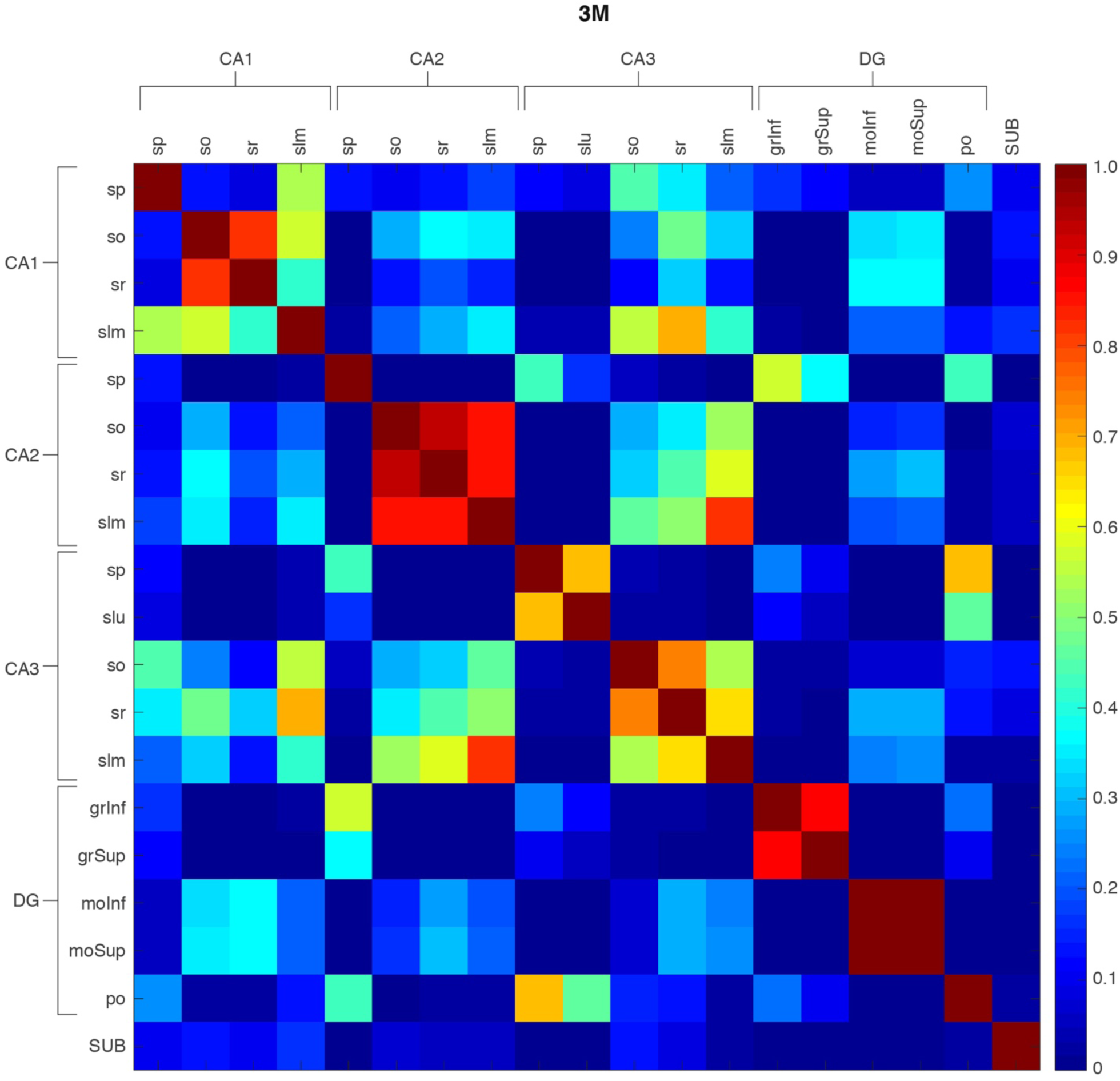
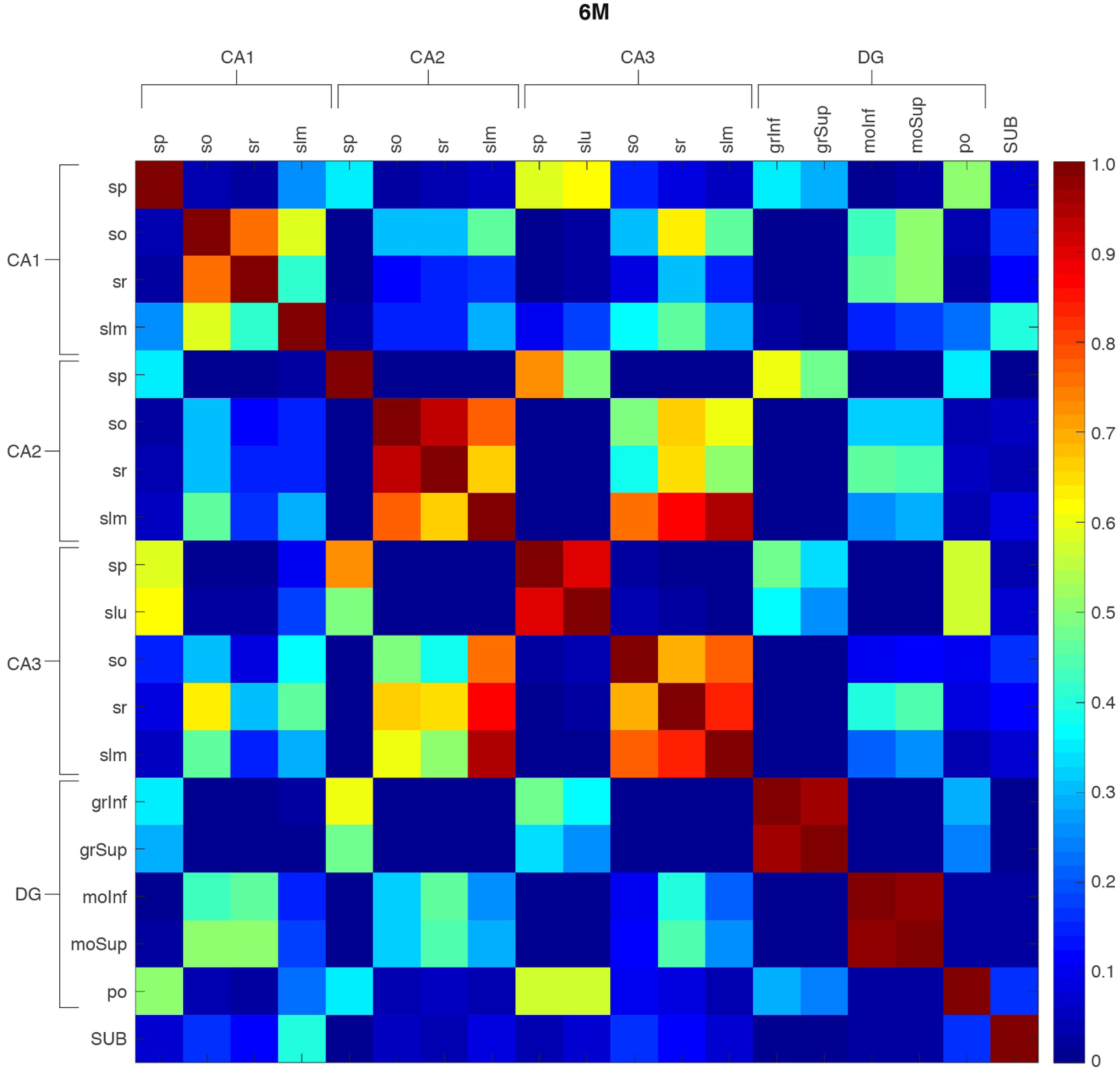
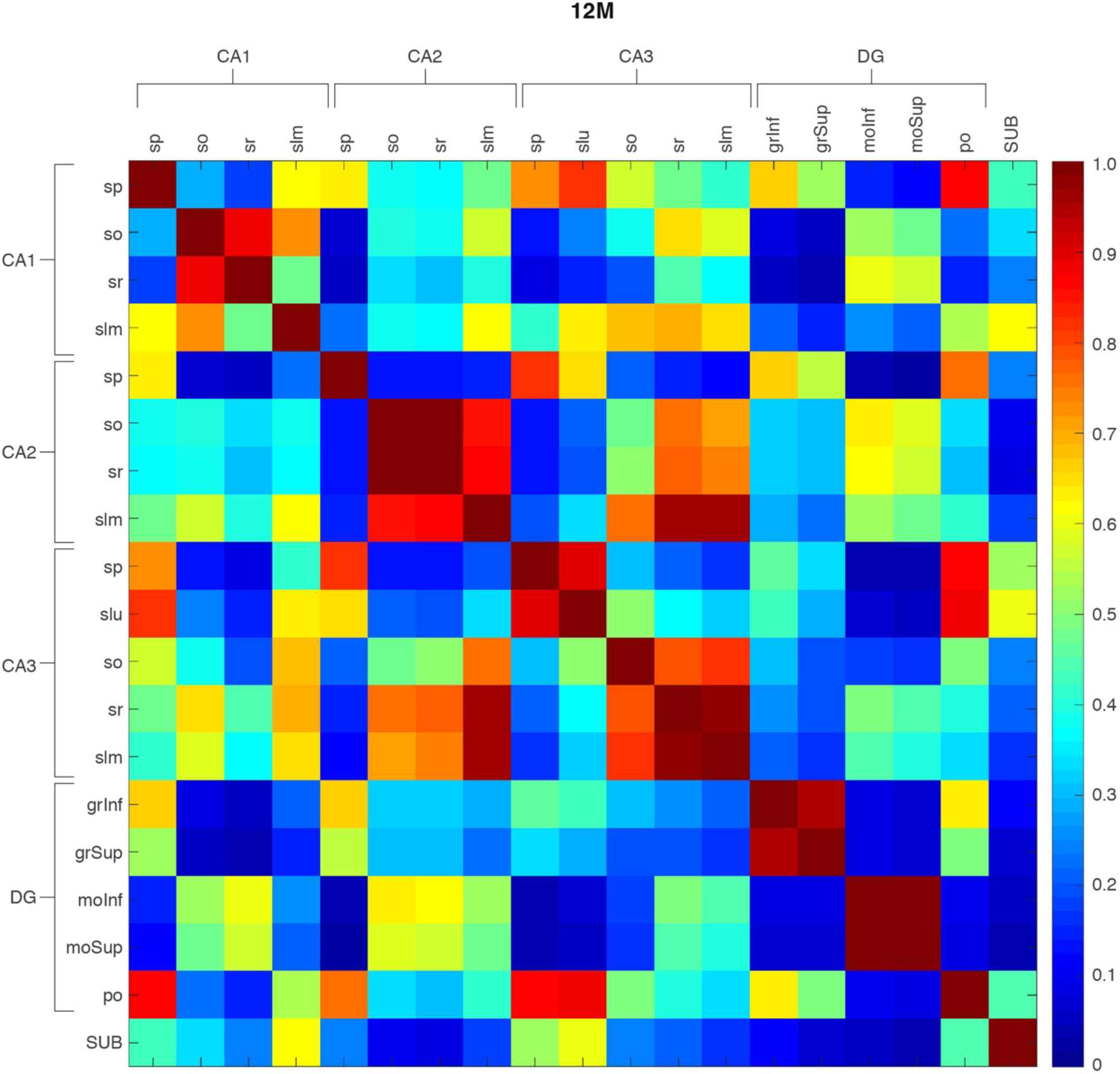
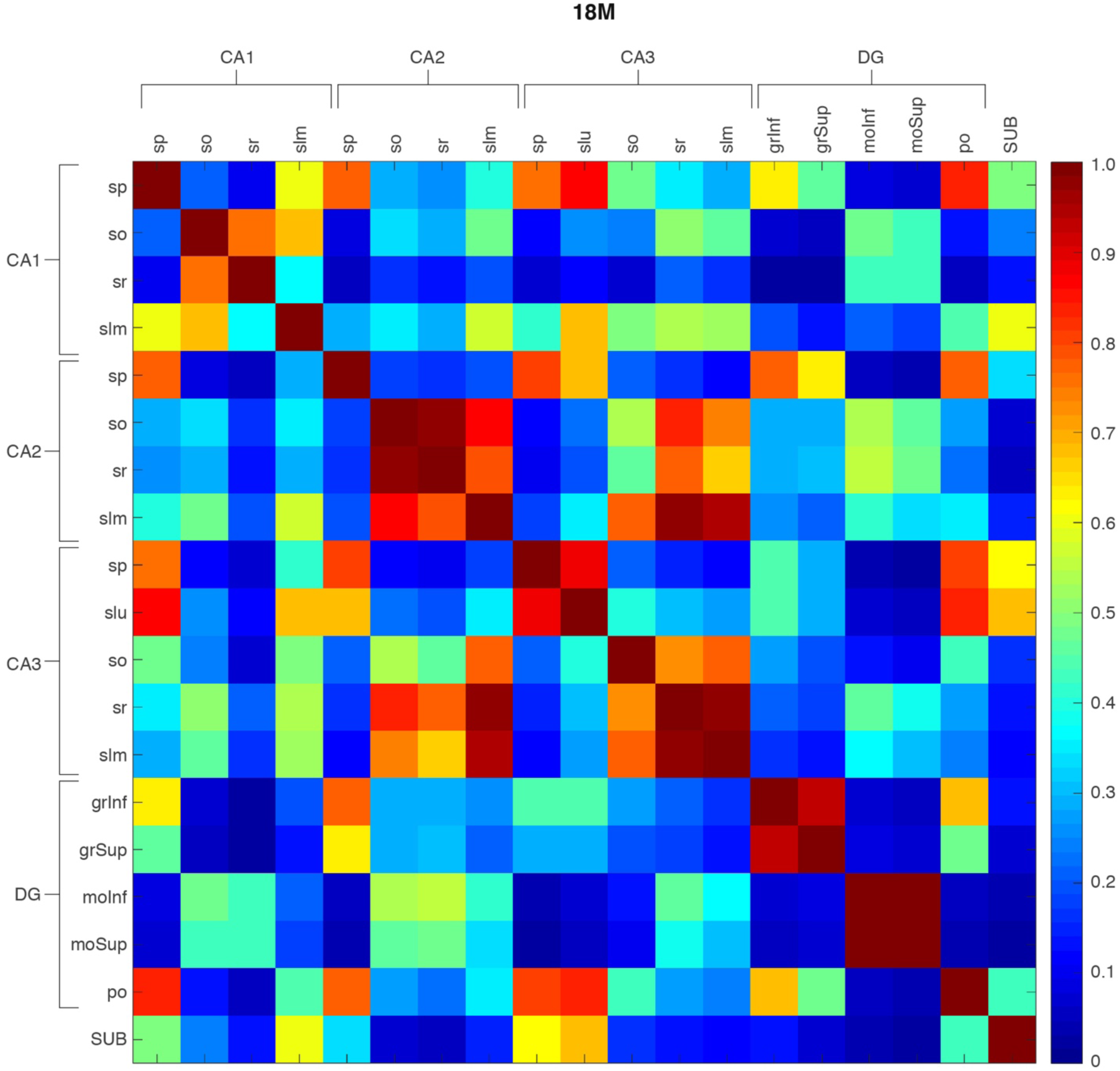
Hippocampal subregion similarity matrices. Matrices of similarities between pairs of subregions (rows and columns) in hippocampal formation from 1W until 18M: 4 subregions from CA1, 4 from CA2, 5 from CA3, 5 from DG, and subiculum. Color scale bar indicates the similarity level ranging from 0 (blue) to 1 (red). The similarity is calculated in the same way as that in Fig. S13. CA1, cornu ammonis 1; CA1so, CA1 stratum oriens; CA1sp, CA1 stratum pyramidale; CA1sr, CA1 stratum radiatum; CA1slm, CA1 stratum lacunosum-moleculare SUB, subiculum. CA2, cornu ammonis 2; CA1so, CA2 stratum oriens; CA2sp, CA2 stratum pyramidale; CA2sr, CA2 stratum radiatum; CA2slm, CA2 stratum lacunosum-moleculare; CA3, cornu ammonis 3; CA3so, CA3 stratum oriens; CA3sp, CA3 stratum pyramidale; CA3slu, CA3 stratum lucidum; CA3 stratum radiatum; CA3slm, CA3 stratum lacunosum-moleculare; DG, dentate gyrus; DGgrInf, DG granular layer, inferior blade; DGgrSup, DG granular layer, superior blade; DGmoInf, DG molecular layer, inferior blade; DGmoSup, DG molecular layer, superior blade; DGpo, DG polymorphic cell layer. SUB, subiculum.

**Fig. S16:**
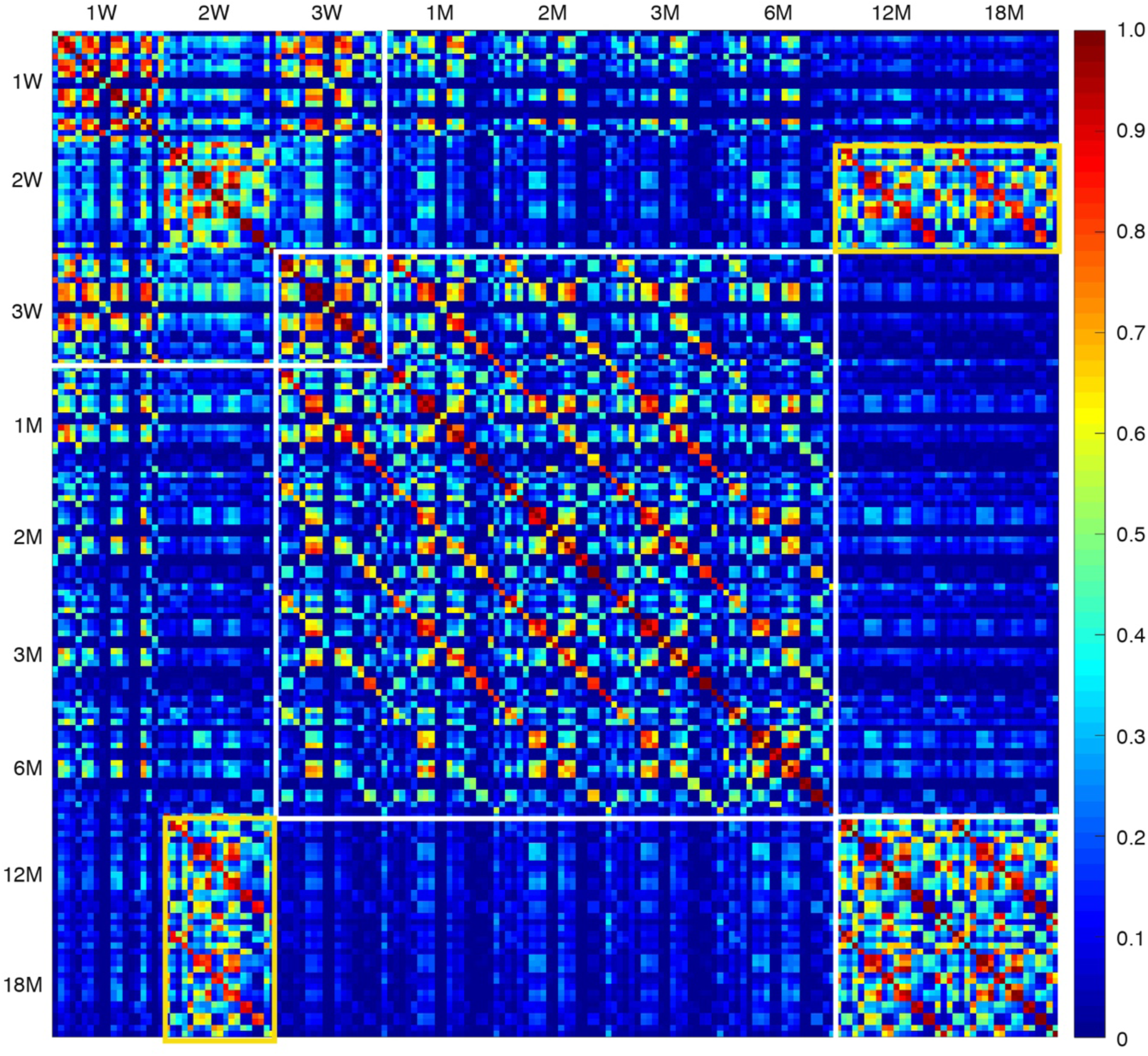
Hypersimilarity matrix comparing all hippocampal subregions across all ages. Enlargement of hypersimilarity matrix in Fig. 3F, where white boxes indicate the main three clusters, which correspond to LSA-I, -II and -III. Yellow box shows increased similarity of the old brain with young brain.

**Fig. S17:**
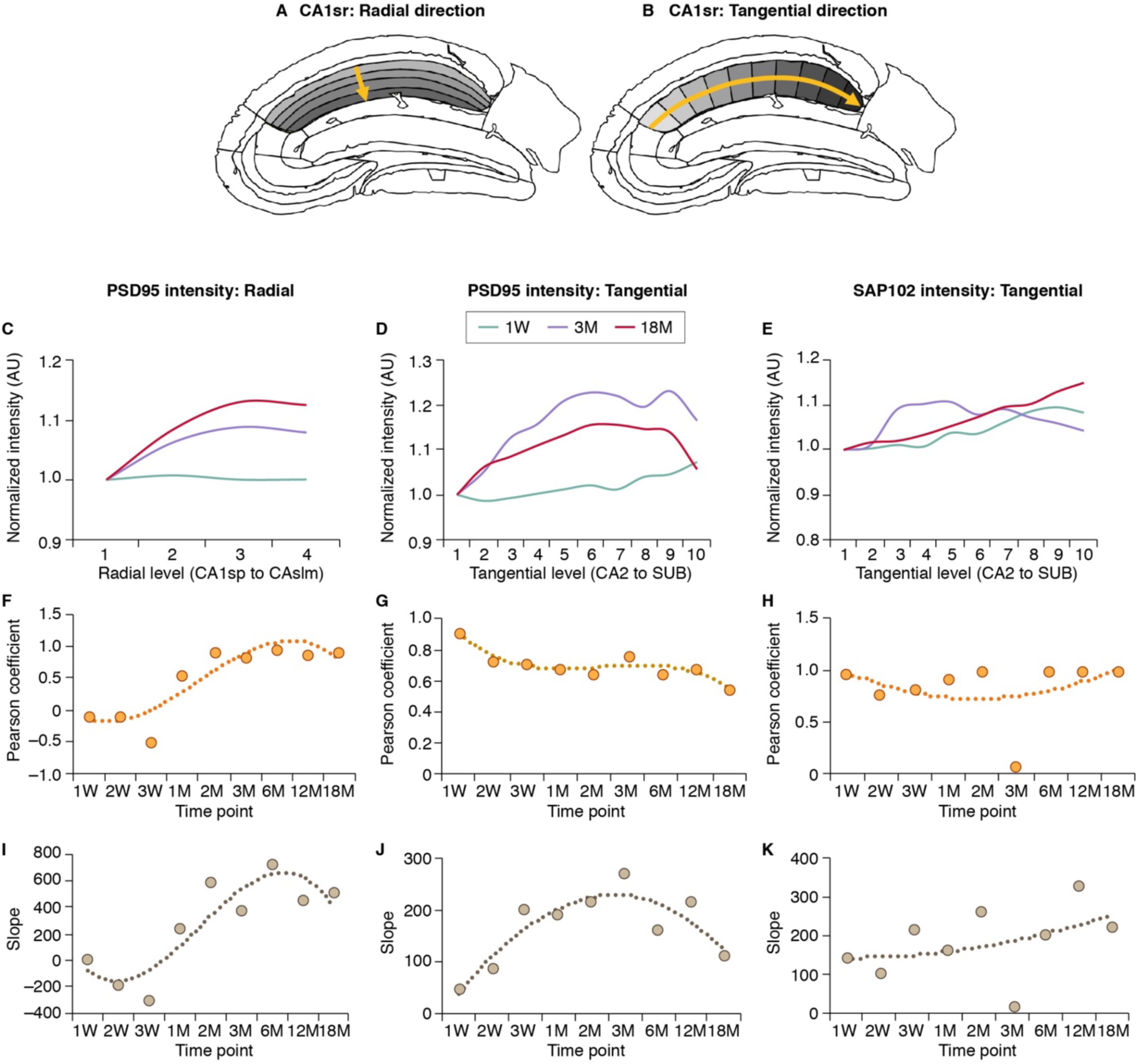
Lifespan trajectories of hippocampal gradients. (A,B) Schematics showing the sub-delineations of CA1sr in the radial (A) and tangential (B) directions used for hippocampal gradient analysis. (C) Smoothed curves showing the PSD95 intensity values along the radial direction of CA1sr (4 levels, CA1sp to CA1slm) at 1W, 3M and 18M (arbitrary units, AU). (D,E) Smoothed curves of PSD95 (D) and SAP102 (E) intensity values along the tangential direction of CA1sr (10 levels, CA2 to SUB) at 1W, 3M and 18M. (F-H) Pearson coefficient of the intensity value versus the gradient level across the lifespan for PSD95 intensity in the radial (F) and tangential (G) directions and for SAP102 intensity in the tangential direction (H). Dashed lines show the polynomial regression curve fitting. (I-K) Slope value of the linear regression curve of the intensity value versus the gradient level across the lifespan for PSD95 intensity in the radial (I) and tangential (J) directions and for SAP102 intensity in the tangential direction (K). Dashed lines show the polynomial regression curve fitting.

**Fig. S18:**
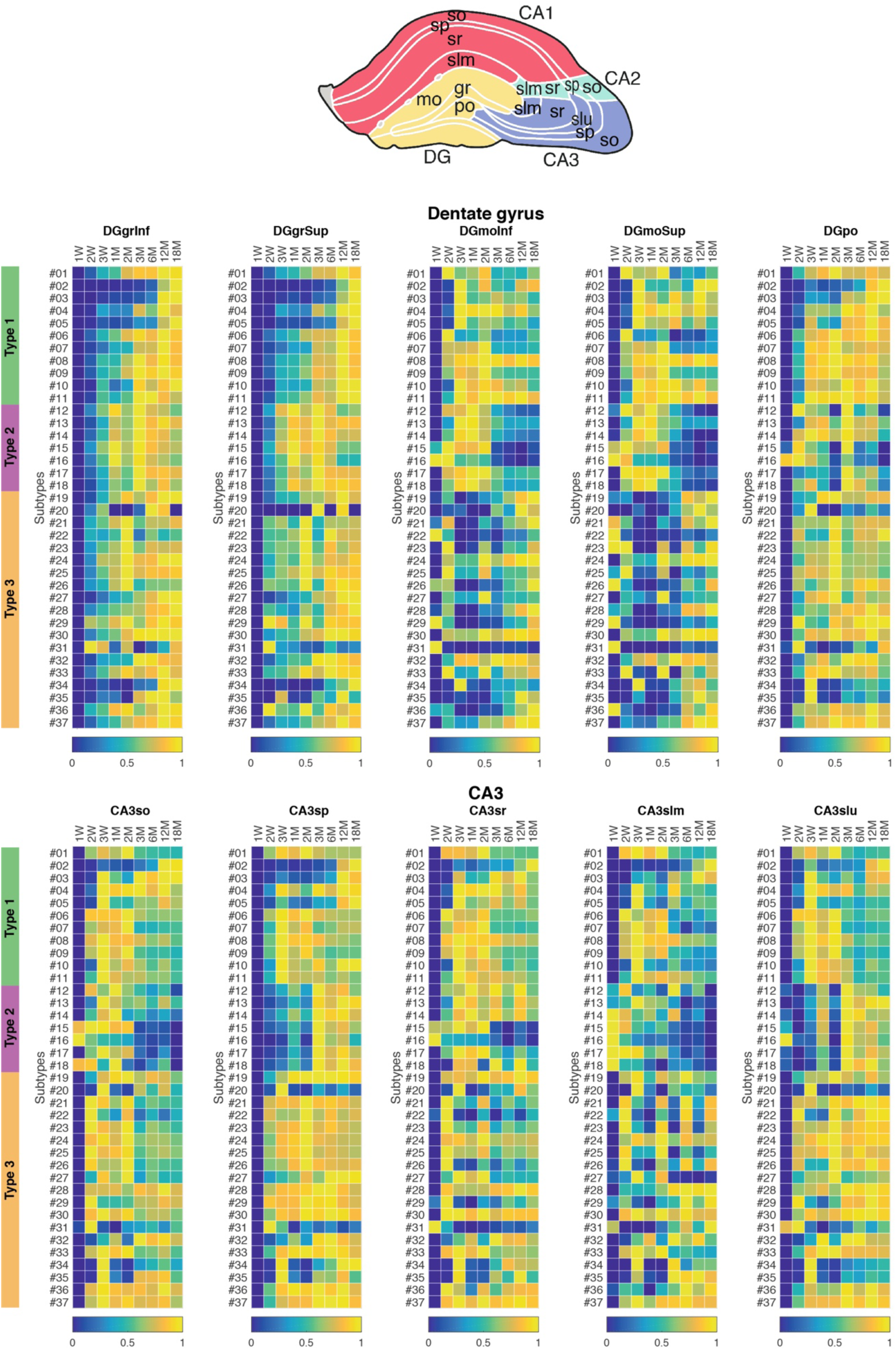
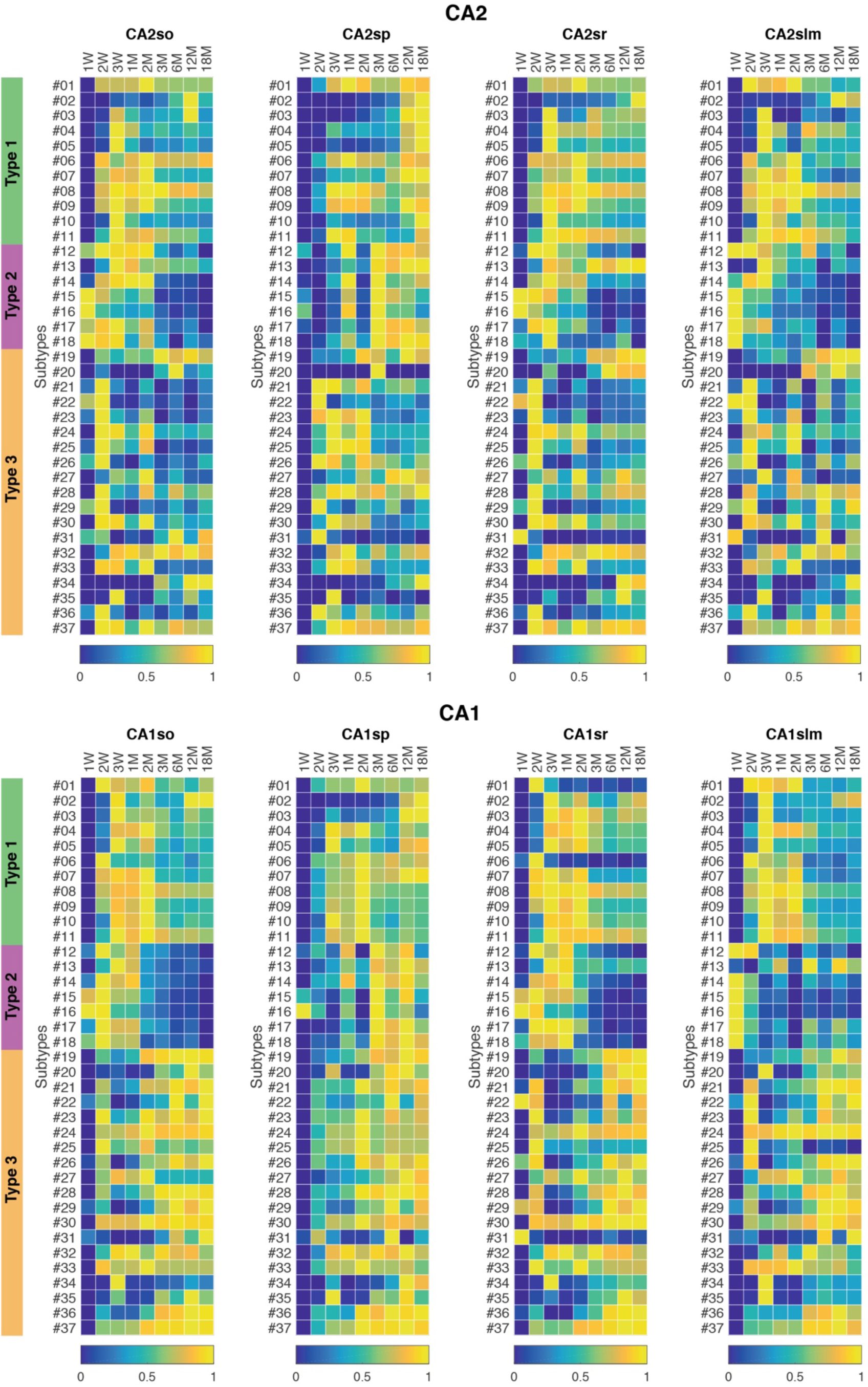
Lifespan trajectories of synapse subtype density in hippocampal subregions. The normalized density of 37 synapse subtypes in subregions of the dentate gyrus, CA3, CA2 and CA1 subregions are shown. Subtype density of each subtype is min-max normalized (0-1) by row to its minimal and maximal density across the lifespan. Color scale bar indicates the normalized subtype densities ranging from 0 (blue) to 1 (yellow). DG, dentate gyrus; DGgrInf, DG granular layer, inferior blade; DGgrSup, DG granular layer, superior blade; DGmoInf, DG molecular layer, inferior blade; DGmoSup, DG molecular layer, superior blade; DGpo, DG polymorphic cell layer. CA3, cornu ammonis 3; CA3so, CA3 stratum oriens; CA3sp, CA3 stratum pyramidale; CA3slu, CA3 stratum lucidum; CA3 stratum radiatum; CA3slm, CA3 stratum lacunosum-moleculare; CA2, cornu ammonis 2; CA1so, CA2 stratum oriens; CA2sp, CA2 stratum pyramidale; CA2sr, CA2 stratum radiatum; CA2slm, CA2 stratum lacunosum-moleculare; CA1, cornu ammonis 1; CA1so, CA1 stratum oriens; CA1sp, CA1 stratum pyramidale; CA1sr, CA1 stratum radiatum; CA1slm, CA1 stratum lacunosum-moleculare;

**Table S1: Names of brain regions and subregions in heatmaps**

The identification of 109 subregions and their order used in heatmaps.

